# Comparative genomics supports that Brazilian bioethanol *Saccharomyces cerevisiae* comprise a unified group of domesticated strains related to cachaça spirit yeasts

**DOI:** 10.1101/2020.12.15.422965

**Authors:** Ana Paula Jacobus, Timothy G. Stephens, Pierre Youssef, Raul González-Pech, Yibi Chen, Luiz Carlos Basso, Jeverson Frazzon, Cheong Xin Chan, Jeferson Gross

## Abstract

Ethanol production from sugarcane is a key renewable fuel industry in Brazil. Major drivers of this alcoholic fermentation are *Saccharomyces cerevisiae* strains that originally were contaminants to the system and yet prevail in the industrial process. Here we present newly sequenced genomes (using Illumina short-read and PacBio long-read data) of two monosporic isolates (H3 and H4) of the *S. cerevisiae* PE-2, a predominant bioethanol strain in Brazil. The assembled genomes of H3 and H4, together with 42 draft genomes of sugarcane-fermenting (fuel ethanol plus cachaça) strains, were compared against those of the reference S288c and diverse *S. cerevisiae*. All genomes of bioethanol yeasts have amplified *SNO2(3)/SNZ2(3)* gene clusters for vitamin B1/B6 biosynthesis, and display ubiquitous presence of *SAM-dependent methyl transferases*, a gene family rare in *S. cerevisiae*. Widespread amplifications of quinone oxidoreductases *YCR102C/YLR460C/YNL134C*, and the structural or punctual variations among aquaporins and components of the iron homeostasis system, likely represent adaptations to industrial fermentation. Interesting is the pervasive presence among the bioethanol/cachaça strains of a five-gene cluster (Region B) that is a known phylogenetic signature of European wine yeasts. Combining genomes of H3, H4, and 195 yeast strains, we comprehensively assessed whole-genome phylogeny of these taxa using an alignment-free approach. The 197-genome phylogeny substantiates that bioethanol yeasts are monophyletic and closely related to the cachaça and wine strains. Our results support the hypothesis that biofuel-producing yeasts in Brazil may have been co-opted from a pool of yeasts that were pre-adapted to alcoholic fermentation of sugarcane for the distillation of cachaça spirit, which historically is a much older industry than the large-scale fuel ethanol production.

## INTRODUCTION

Fermentation of sugarcane extracts to distillate the “cachaça” spirit has been intertwined with the Brazilian culture since the 16th century. In contrast, the production in Brazil of the biofuel ethanol from sugarcane juices and molasses only reached economic prominence in the 1970s, when governmental stimulus propelled the establishment of a nationwide biofuel industry (Basso et al., 2011). As a result, Brazil is currently the world’s second largest producer of ethanol, with the production of ~35.5 billion liters in the 2019/20 period, only behind the United States of America (Brazilian Sugarcane Industry Association, http://www.unicadata.com.br/). Ethyl alcohol is derived from sugarcane sucrose via microbiological fermentation, primarily by *S. cerevisiae* strains. In Brazil, industrial alcoholic fermentation of sugarcane operates in a continuous manner (Basso et al., 2008; Della-Bianca et al., 2013; Lopes et al., 2015). When a fermentation batch is over, yeast cells are centrifuged, washed, treated with sulfuric acid (to hamper bacterial contamination), and then reinoculated for a new round of fermentation (i.e., two per day across a season of up to nine months). During the early period of 1970s-1990s, a common practice in the distilleries was to kick-off a fermentation season with starter cultures consisting of commercial *S. cerevisiae* strains (mainly baker’s yeasts) (Basso et al., 2008; Lopes et al., 2015). However, it was soon observed that the inoculated yeasts were quickly displaced by *S. cerevisiae* strains that intruded the fermentation vats. Those contaminants were often referred to as “indigenous” or “wild-type” yeasts, although no phylogenetic evidence actually corroborated that notion (Basso et al., 2008; Argueso et al., 2009; Antonangelo et al., 2013; Della-Bianca et al., 2013; Lopes et al., 2015). Regardless of their ambiguous provenance, “intruding” strains such as *S. cerevisiae* SA-1, BG-1, CAT-1, VR-1 and PE-2, were systematically isolated, characterized, and reintroduced into distilleries, becoming the standard bioethanol selected strains still in use today (Basso et al., 2008; Lopes et al., 2015). Physiological experiments conducted on these strains revealed excellent fermentation profiles and exceptional capacity to withstand the harsh conditions of industrial fermentation that include high salt and sugar content (18 to 22 % (w/w) total reducing sugars), high temperature (usually reaching 37 °C), presence of bacterial contaminations (e.g., by *Lactobacilli*), high ethanol content (up to 12 % v/v), and low pH (1.8-2.5) due to sulfuric acid treatment (Basso et al., 2008; Della-Bianca et al., 2013; Walker and Basso, 2020). Overall, sugarcane fermentation represents an active system where contaminants (yeasts and bacteria) from the exogenous environment dynamically compete with inoculated strains. Even today, when traditional bioethanol strains (e.g., PE-2, BG-1 and CAT-1) are favored as starter cultures, they are often displaced over the year by exogenous *S. cerevisiae* (Della-Bianca et al., 2013; Lopes et al., 2015). This begs the question about the phylogenetic provenance of strains that enter and prevail through the fermentation season.

Based on comparative genomic hybridization, Roche 454 and emerging Illumina technologies, the genome sequencing and structural analyses of the *S. cerevisiae* PE-2 and CAT-1 strains were reported about ten years ago (Argueso et al., 2009; Stambuk et al., 2009; Babrzadeh et al., 2012). Those studies represent a milestone in our understanding of Brazilian bioethanol strains and substantiated the notion that these yeasts are often heterothallic diploids containing a highly polymorphic set of chromosomes. In particular, subtelomeric regions represent a fluidic part of the genome where frequent ectopic recombination events dynamically alter copy number of genes that encode industrial relevant traits, such as stress response and nutrient acquisition. This is the case for the *SNO2(3)/SNZ2(3)* gene cluster involved in pyridoxal 5’-phosphate (PLP, vitamin B6) and thiamin diphosphate (ThDP, vitamin B1) biosynthesis, which is represented by four copies in the *S. cerevisiae* PE-2 haploid derivative JAY291 (Argueso et al., 2009). Amplifications of the *SNO2(3)/SNZ2(3)* cluster, further observed in the CAT-1, VR-1, BG-1, and SA-1 strains, are thought to confer selective advantages to the yeasts when grown in conditions of thiamin repression under high sugar concentrations, possibly found in the industrial sugarcane fermentations (Stambuk et al., 2009).

However, the *SNO2(3)*/*SNZ2(3)* cluster and other gene families located at the repeat-rich subtelomeric regions cannot be adequately placed in genome assemblies using only short-read sequence data. To overcome this problem, long-read sequence data (e.g., generated using PacBio/Oxford Nanopore technologies) can better resolve these repetitive regions and have been used to improve the chromosomal positioning of telomeres, solo-long terminal repeats (solo LTRs), complete Ty transposable elements, and more importantly, of multi-paralog gene families (McIlwain et al., 2016; Salazar et al., 2017). High-throughput sequencing technologies are also facilitating multi-taxa population genomics studies that are clarifying the evolutionary relationships among key industrial and environmental groups of the *S. cerevisiae sensu stricto* complex (Gallone et al., 2016; Barbosa et al., 2018; Legras et al., 2018; Peter et al., 2018). For example, some studies have hinted at the close evolutionary relationship of Brazilian bioethanol and cachaça strains to European wine yeasts, supporting that two sequential domestication events occurred in the evolutionary lineage of cachaça yeasts; i.e., the primary domestication of wild Mediterranean oak *S. cerevisiae* originating the European wine yeasts (Almeida et al., 2015), and a recent secondary adaptation in Brazil of wine strains for fermenting sugarcane extracts to produce the cachaça spirit (Barbosa et al., 2018). To gain more insights into the evolutionary history of Brazilian sugarcane-fermenting yeasts we sequenced and assembled two haploid genomes derived from the diploid isolate *S. cerevisiae* PE-2. Incorporating 42 other Brazilian biofuel/cachaça strains in a comparative analysis, we systematically assessed gene content, polymorphisms, and copy-number variation (CNV) that represent features conserved among bioethanol yeasts. For the first time using a scalable alignment-free phylogenetic approach, we evaluated the evolutionary relationship among diverse industrial *S. cerevisiae* based on whole-genome sequences from 197 taxa. Our results provide unambiguous support to the notion that Brazilian bioethanol yeasts comprise a uniform group of domesticated and phylogenetically related yeasts.

## MATERIALS AND METHODS

### Genome sequencing of H3 and H4

A frozen stock derived from the original isolate *Saccharomyces cerevisiae* PE-2 was obtained from the collection of Prof. L. C. Basso at the University of São Paulo, Piracicaba, Brazil (Basso et al., 2008). From this source, propagated cultures were sporulated and a single ascus was dissected to isolate four sibling spores. Two of them, denominated haploid H3 (*MATα*) and haploid H4 (*MATα*), were propagated and targeted for DNA extraction (DNeasy Plant Mini Kit, QIAGEN), DNA fragmentation (NEBNext dsDNA Fragmentase, New England BioLabs), and genomic library construction (Illumina TruSeq DNA PCR-Free Low Throughput Library Prep Kit, Illumina). The generated paired-end libraries were quantified by qPCR (KAPA Library Quantification Kit, Roche) and then sequenced at the Federal University of Rio Grande do Sul, Brazil, using the Illumina MiSeq platform (2×1300 paired-end; MiSeq Reagent Kit v3 supporting 600-cycles, Illumina). The resulting sequence reads were filtered, retaining only those with Phred quality scores ≥30 and length ≥75 bases. This resulted in a final set of 11,535,554 paired reads from H3, and 12,366,234 paired reads from H4 for subsequent genome assembly (below). In parallel, genomic DNA samples (isolated from H3 and H4 as described above) were sent for PacBio long-read sequencing using the RS II platform at DNA Link (Seoul, Korea), for one SMRT cell each for H3 and H4. The resulting long-reads were processed using Canu (Koren et al., 2017), resulting in 83,983 and 59,078 corrected reads from H3 and H4, respectively.

### Assembly and manual curation of H3 and H4 genomes

We assembled the H3 and H4 genomes independently using CLC Genomics Workbench version 8.0.1 (Qiagen, https://digitalinsights.qiagen.com), incorporating the Microbial Genome Finishing Module (https://digitalinsights.qiagen.com/plugins/clc-genome-finishing-module/). For each haploid sample, a four-step progressive assembly approach was adopted which included indepth visual inspection and manual curation of each phase of the process. First, Illumina reads were assembled using CLC *de novo* assembler (a de Bruijn graph algorithm), resulting in 384 and 416 contigs for H3 and H4, respectively. Second, the assembled contigs were aligned and superscaffolded against the chromosomal-scale genome sequences of the model reference *S. cerevisiae* S288c using BLASTN (Altschul et al., 1997), as implemented in the “contig aligner tool” of the CLC Genome Finishing Module. Overlapping and redundant contigs were merged and putative chromosomal structures were identified by concatenating contigs following the reference genome. A string of “Ns” was provisionally assigned to represent gaps of unknown length separating adjacent contigs. Third, a manual inspection of sequential contigs that overlaid on S288c genome sequences was conducted, chromosome-by-chromosome. When broken connections between contigs relative to S288c are represented by small (< 500 bp) repetitive sequences (e.g., solo LTRs or regions of genes with highly similar paralogs), we mapped the H3 or H4 reads to the S288c genome to obtain a consensus sequence that bridge those specific gaps [i.e., a reference-guided assembly (Gnerre et al., 2009; Lischer and Shimizu, 2017)]. After joining neighboring contigs, reads were realigned to the scaffold to verify that the number of reads mapped over the proposed connection was uniform (i.e., uniform read coverage, reads fully aligned, and unbroken read pairs). The CLC burrows-wheeler aligner was run at a stringent setting (minimal 0.9 length fraction and 0.9 similarity fraction) throughout the assembly process to ensure high accuracy of the alignments. Fourth, PacBio long-read data were used to resolve larger and/or more complex gapped regions that correspond to complete Ty elements (~6,000 bp), genes with internal repetitions (e.g., the *FLO* family), or repeated subtelomeric regions (Koren and Phillippy, 2015). Corrected long reads or unitigs generated using Canu (Koren et al., 2017) were unambiguously assigned using BLASTN to unique (non-repeated) regions flanking both sides of any gapped regions. By realigning corrected reads and/or unitigs (CLC multiple sequence aligner, slow and accurate mode; gap open cost 10.0, gap extension cost 1.0), consensus sequence that bridge each gapped region was identified. In a similar approach, PacBio long reads and Canu unitigs assigned to chromosome extremities were used to extend core chromosome structures into subtelomeric regions and resolve segmental duplications involving paralog-rich gene families. Only one major Canu unitig (521) related to the H4 genome could not be properly assigned to one of three alternative positions related to the tips of chromosomes XIII, XV, and XVI (See **Figure 1**, bottom). Finally, where necessary, Illumina short reads were used to correct for incorrect bases inherent of PacBio assemblies (Koren and Phillippy, 2015). We tested the resulting H3 and H4 assemblies by thoroughly searching for inconsistencies in Illumina read-mapping profiles over these genomes. Diagnostic tests were implemented by the “analyze contigs tool” of the CLC Genome Finishing Module and included detection of sudden changes in read depth, regions with higher/lower coverage, unaligned read ends, and broken paired-end reads. Those features are usually associated with misassembled regions or the presence of structural polymorphisms (Suzuki et al., 2011; Nagarajan and Pop, 2013; Yang et al., 2019). Only a few problematic regions were retrieved that are related to highly repeated telomeric regions and Ty elements that were assembled relying almost exclusively on PacBio sequence reads of suboptimal quality. In addition, many short (<100 bases) unresolved polymeric structures, which cause errors for both Illumina and PacBio sequencing (Laehnemann et al., 2016), were assigned in our assembly with a string of 20 “N”s. Altogether, quality check reassured that our H3 and H4 genome assemblies generated reliable draft sequences for the 16 yeast chromosomes, mitochondrial genome, 2-micron plasmid, and could assign positions for highly repeated subtelomeric gene families that are key to this study. Genome sequence data for H3 and H4 are available at NCBI under the BioProject accession number PRJEB31792.

**FIGURE 1.**
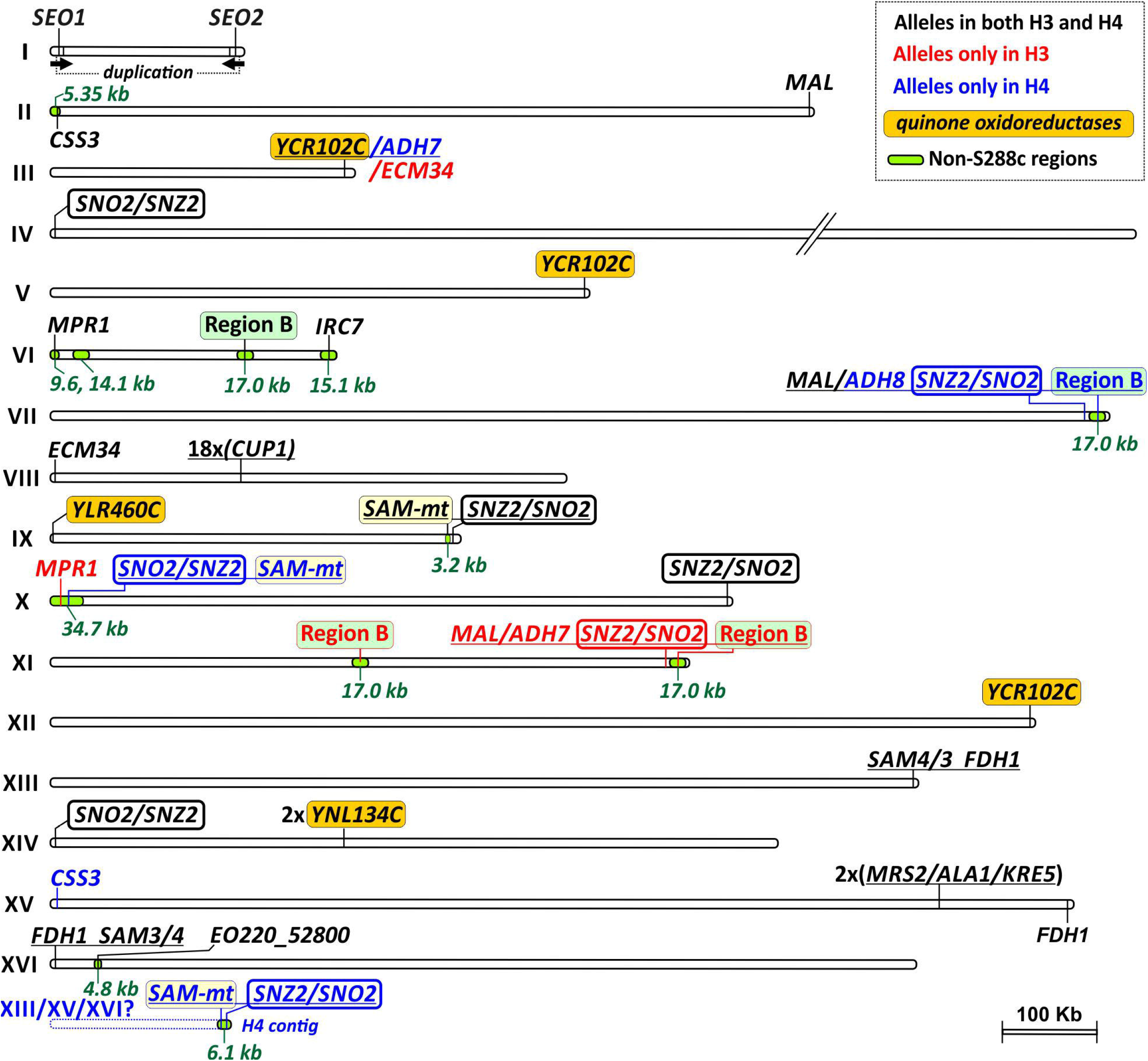
Chromosomic map of H3 and H4 showing regions with increased copy numbers when compared to the reference S288c. *Loci* present in H3 and H4 that are absent from S288c are also displayed as green regions overlaying chromosome structures. The gene content of each labeled region is schematically depicted on the **Figure 2**. The sixteen chromosomes are shown in scale with the exception of a shortened (//) version of the Chr. IV. An unplaced subtelomeric region of the H4 genome (corresponding to a single H4 contig), which contains the *SAM-m*t and *SNZ2*/*SNO2* locus, is depicted below the Chr. XVI and has a putative location downstream the *FDH1* region (its uncertain location at either one of the Chrs. XIII, XV, or XVI is indicated by a question mark). Regions and genes present in both H3 and H4 genomes are labeled in black. Regions and genes present only in H3 have their labels colored in red, while those exclusive to H4 are displayed in blue. Four key multi-copy regions are highlighted by colored boxes framing their labels. They correspond to genes of the *quinone oxireductase* family (orange box), Region B (green box), *SAM-dependent methyltransferases* (*SAM-mt*, yellow), and the cluster *SNO2(3)/SNZ2(3)* (box with thick lines).

### Genome annotation and gene-content comparison against S288c reference

We transferred the annotations *en bloc* from the reference S288c (GenBank accession number GCA_000146045.2) to the assembled H3 and H4 chromosomes using the “annotate from reference” tool from the CLC Genome Finishing Module. In parallel, we generated a genome-wide prediction of all possible coding regions (ORFs) larger than 150 bp. The CLC Graphical interface was used to visually inspect and manually reconcile the two types of information. In addition, ORF predictions allowed us to annotate (and catalogue) genes or regions of H3 and H4 genomes that are not represented in S288c (below). The precise structure of each newly predicted feature was corroborated by comparisons against the S288c syntenic counterparts, specifically to correct any discrepancies in gene length, and identify possible cases of premature stop codons in H3 and H4 genomes. Finally, using custom in-house scripts, protein translations and index tags were assigned to each gene locus, and genomic features were further inspected, and corrected where necessary for any inconsistencies between gene and predicted protein lengths. In our analyses of gene content, we excluded uncharacterized and dubious ORFs (Lin et al., 2013) of short length (≤ 150 bp), and genes related to Ty elements and telomeric regions, given their highly repetitive nature (Engel et al., 2014). Annotation of non-coding RNAs were transferred from the S288c genome and tRNA structures were predicted by using tRNAscan-SE (Lowe and Chan, 2016).

Most cases of non-S288c genes were already discovered during our annotation process within regions in which no S288c annotations were transferred, or in cases where predicted ORFs found no counterpart in S288c. In addition, the H3 and H4 genomes were inspected by TBLASTN for the presence of previously described non-S288c genes (Borneman et al., 2011; McIlwain et al., 2016). Regions potentially introgressed from *Saccharomyces paradoxus* were identified by mapping H3 and H4 reads against the genome of either *S. paradoxus* strain [CBS432 (ASM207905v1) or UFRJ50816 (ASM207914v1)], and simultaneously to S288c. H3/H4 reads that mapped against *S. paradoxus* genome revealed chromosomal segments that are putatively introgressed into the genome *S. cerevisiae* PE-2. Those regions were further queried using BLASTN searches against the nucleotide collection database of GenBank to fulfil the criteria of returning top hits to *S. paradoxus* instead of any *S. cerevisiae* taxa. OrthoFinder (Emms and Kelly, 2019) was used to infer homologous protein families among H3, H4 and S288c, from which orthologs of H3/H4 (i.e. PE-2) not recovered among the S288c predicted proteins, or vice versa, were identified. Genes (and genomic regions) that are absent in PE-2 relative to S288c were alternatively identified by mapping of H3 and H4 Illumina reads against the S288c genome (“coverage analysis”, CLC Genome Finishing Module). Most cases of gene-copy number variation were identified during the earlier assembly of difficult-to-resolve regions, usually related to subtelomeres. For this reason, we used all subtelomeric regions as a query to search, using BLASTN, for further cases of segmental duplications and catalog CNVs between H3, H4 and S288c. In a complementary approach, OrthoFinder (Emms and Kelly, 2019) was used to identify cases where protein families were expanded or contracted. CNVs were further probed based on analysis of read coverage (below).

### Comparative genomic analyses across *S. cerevisiae* strains

For SNP comparison, we focus on H3, H4, JAY291 (another PE-2 haploid derivative), and S288c. Genomes of *S. cerevisiae* S288c (GCA_000146045.2) and JAY291 (ASM18231v2) were downloaded (https://www.ncbi.nlm.nih.gov/assembly/). The first step in this analysis was to mask in the four genomes all repetitive regions such as telomers, transposable elements, rRNA and highly duplicated subtelomeric gene families. Second, unmasked regions corresponding to core chromosomes were aligned using nucmer from the MUMmer 3.0 package (Kurtz et al., 2004). Finally, tools from the same package were used for SNP count and characterization of aligned regions. GFF files were used to distinguish genic and intergenic regions in the analyzed genomes.

For more-comprehensive comparative genomic analysis, we focused on 44 bioethanol and cachaça strains (hereinafter referred to as the Brazilian bioethanol group; **Supplementary Table 1**). They correspond to isolates from ethanol plants, mostly located in the state of São Paulo, Brazil, and include the haploid genomes of H3, H4, JAY291 (Argueso et al., 2009), the diploid genome of BG-1 (Coutoune et al., 2017), five strains sequenced by Gallone et al. (Gallone et al., 2016), plus 35 strains (mostly diploid genomes) from the 1,002 Yeast Genomes Project (Peter et al., 2018). For enrichment analysis of distinct genomic features (including genes and alleles) within those strains, a query specific for each feature was used to search by BLASTN against the genomes of the Brazilian bioethanol group (**Supplementary Table 1**) plus a dataset of 976 strains derived from the 1,002 Yeast Genomes Project (Peter et al., 2018). Query sequences, specific selection criteria, and BLAST settings for this analysis are described in **Supplementary Table 2**. A genetic feature was considered as present even in heterozygosis (i.e., both alternative alleles were present). The statistical significance of the results was assessed using Fisher’s exact test (implemented by GraphPad Prism 7.00, GraphPad Software, La Jolla California USA) comparing bioethanol to non-bioethanol strains and selecting for *p*-values < 0.05 as cut-off.

For analysis of CNVs, we extracted genes/regions of S288c for which copy-numbers were found to be amplified or contracted in H3 and H4, or in genomes of other bioethanol strains (**Supplementary Table 3**). Sequences from H3 and H4 were selected only in cases where no S288c counterpart was available. A nucleotide sequence **(Supplementary Sequence File 1)** was assembled by concatenating these genes/regions. Flanking regions of at least 250 bp (usually 500-1000 bp), corresponding to promotor (upstream) or terminator (downstream) parts of genes, were added to separate the regions and to ensure full alignments of reads at the 5’- or 3 ‘-end of the genes. Illumina reads from 40 Brazilian bioethanol strains (except H3 and H4) were downloaded from The European Nucleotide Archive (ENA, https://www.ebi.ac.uk/ena), filtered by quality, and mapped against the S288c chromosomes to obtain the average genome coverage for each strain (**Supplementary Table 4**). For comparison, we included 10 non-bioethanol strains (**Supplementary Table 5**) randomly picked from the dataset of the 1,002 Yeast Genomes Project (Peter et al., 2018). Read mappings were conducted using a burrows-wheeler aligner (BWA) (Li and Durbin, 2009). After obtaining strain-specific whole genome coverage, reads from each strain were mapped against the concatenated dataset (**Supplementary Sequence File 1**) to interrogate for CNVs over the selected genes/regions. CNVs were calculated by normalizing read coverage of the coding sequence to its length, and to the background read coverage of the genome. As an example, **Supplementary Table 3** describes the numbers and calculations of CNVs for H3. For H3 and H4, we retrieved very similar CNV values estimated with our read depth approach to the numbers derived from our genome assembly (**Supplementary Figure 1**; For H3: Pearson’s *r* = 0.9976, Confidence Interval 95% [0.9956, 0.9987], *p* ≤ 0.0001; For H4: Pearson’s *r* = 0.9978, Confidence Interval 95% [0.9959 to 0.9988], *p* ≤ 0.0001). Absolute copy numbers for probed genes/regions were subtracted from the numbers derived from the S288c genome annotation (GCA_000146045.2) to compose a matrix of differential copy numbers that was transformed in a heatmap representation using the GraphPad Prism 7.00 package (GraphPad Software, La Jolla California USA). A Mann-Whitney U test (implemented by GraphPad Prism 7.00) was applied to infer statistical significance of CNVs found in strains of the bioethanol group when compared to the non-bioethanol control.

### Alignment-free phylogenetic analysis

We adopted an alignment-free (AF) approach (Chan et al., 2014; Bernard et al., 2019) for inferring phylogenetic relationships from whole-genome sequences, based on similarity of shorter, sub-sequences at length *k* (i.e., *k*-mers). For this analysis, we used a total of 197 genomes combining the dataset of bioethanol strains (**Supplementary Table 1**) and 157 genomes from a study of industrial yeasts (Gallone et al. 2016; **Supplementary Table 6**). Genomes of the *Saccharomyces paradoxus* strains CBS432 (ASM207905v1) and UFRJ50816 (ASM207914v1) were included as outgroup. The majority of these genomes are in chromosome-level resolution based on scaffolding against S288c, except for genomes of 35 bioethanol strains obtained from the 1,002 Yeast Genomes Project, and BG-1. These 36-genomes data, although representing draft, fragmented assemblies that could bias enumeration of *k*-mers, were included to allow for comprehensive inclusion of available bioethanol strains. To ensure comparability of these genomes against the others in our dataset, we refined the 36 genome assemblies using Chromosomer (Tamazian et al., 2016) to reconstruct chromosome-level assemblies, with S288c as reference. Where applicable, short (<1 Kb) contigs were removed, and repeats were masked using RepeatMasker (http://www.repeatmasker.org/). Enumeration of *k*-mers for each set of genome sequences was conducted using Jellyfish (Marcais and Kingsford, 2011). The optimal *k* for phylogenetic analysis is based on a comprehensive assessment across all genomes over an increment of *k* (Greenfield and Roehm, 2012), the threshold at which the percentage of unique *k*-mers for each genome reach a plateau (**Supplementary Figure 2**). We chose *k* = 21 based on this assessment, as this value provides a saturated level of unique *k*-mers in each genome while providing sufficient information of shared *k*-mers among the genomes for calculation of pairwise distance. Following Chan et al. (2014), for each possible genome pair, we calculated a pairwise distance based on the 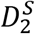 statistic (Wan et al., 2010). The pairwise distance matrix was used to infer a neighbor-joining tree, using *neighbor* implemented in PHYLIP (Felsenstein, 1989).

## RESULTS

### H3, H4 and JAY291 represent alternative polymorphic genomes of the bioethanol strain *S. cerevisiae* PE-2

We isolated two alternative spores derived from a *S. cerevisiae* PE-2 single tetrad. For each haploid, we generated genome assemblies combining Illumina MiSeq short-read and PactBio RS II long-read data, guided by the reference genome of *S. cerevisiae* S288c at chromosomal resolution (see Materials and Methods). We recovered genome sequences for the corresponding chromosomes of H3 and H4 (**Table 1**), including most of the subtelomeric repeats. These new genome assemblies of the Brazilian bioethanol strain *S. cerevisiae* PE-2 represent alternatives to the one from a different haploid isolate (JAY291) that was previously sequenced with Illumina (GAIIx) and Roche 454 technologies (Argueso et al., 2009).

**TABLE 1.**
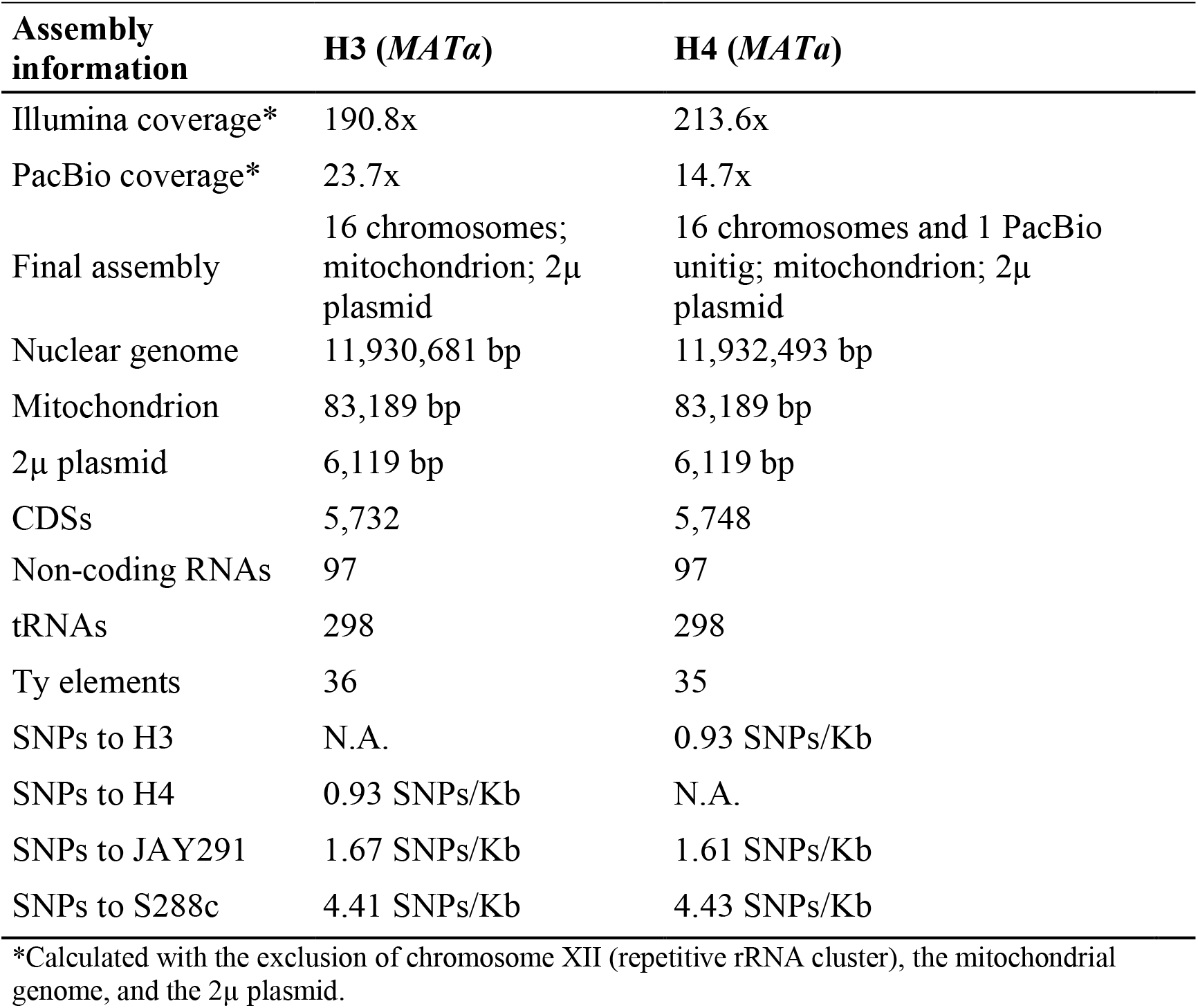
H3 and H4 genome assembly and SNPs numbers.

Direct comparison of non-repetitive genome regions between H3 and H4 estimated on average 0.93 SNPs/Kb (**Table 1** and **Supplementary Table 7**). Loss-of-heterozygosity (LOH) is observed across many chromosomal regions (with only 0.03 SNPs/Kb identified in Chr. VI), consistent with the fact that H3 and H4 originated from sibling spores that were derived via meiosis from the same parental diploid genome. Patterns of LOH across the genome were also previously observed in segregants of JAY270, the parental diploid of JAY291 (Rodrigues-Prause et al., 2018; Sampaio et al., 2019). The higher polymorphism density observed between H3 and H4 along Chrs. IX and XIII (1.95 and 1.88 SNPs/Kb, respectively; **Supplementary Table 7**) is close to an early estimation of about 2.0 SNPs/Kb for the *S. cerevisiae* PE-2 diploid genome (Argueso et al., 2009). H3 and H4 are also more polymorphic when compared to S288c (about 4.42 SNPs/Kb) and to JAY291 (about 1.64 SNPs/Kb) (**Supplementary Tables 8, 9, 10, 11**). Substantial heterozygosity among H3, H4, and JAY291 indicates that these monosporic isolates comprise three distinct genomic sets representing the bioethanol strain *S. cerevisiae* PE-2.

Besides heterozygosity at nucleotide level, H3 and H4 genomes further display substantial structural polymorphisms, specially at chromosome tips. For example, at the Chr. X left-end of H4, there is a 34.7 kb region (spanning 14 genes) that is absent in S288c and H3 (**Figure 1**, **Supplementary Figure 3**). On the other hand, at the corresponding location on Chr. X, H3 carries within a 32.7 kb segment (**Figure 1** and **Figure 2A**) the *AQY3* cluster (*THI5/AAD6/AGP3/AQY3/DAK2/ZNF1/IMA5/HXT8*) and the *MPR1*/C2U11_6360 genes. The same ten-gene region is located at the Chr. VI left-end of both H3 and H4, and represents the long form of Chr. VI previously noticed in *S. cerevisiae* PE-2 (Argueso et al., 2009). Further structural variations at chromosome termini account for differences between H3 and H4 in copy numbers of some key genes (**Figure 1** and **Figure 2**). Some examples are the *SNO2(3)/SNZ2(3)* cluster (five copies in H3 and seven in H4, **Figure 2B**), *SAM-dependent methyl transferases (SAM-mt*, one in H3 and three in H4, **Figure 2B**), Region B (three in H3 and two in H4, **Figure 2C**), and the *MAL* cluster (three in H3 and two in H4, **Figure 2D**). Overall, genome assemblies of H3 and H4 corroborate the notion that in bioethanol strains (sub)telomeres are fluidic genomic regions, accounting for dynamic CNVs of potentially adaptive genes (Argueso et al., 2009).

**FIGURE 2.**
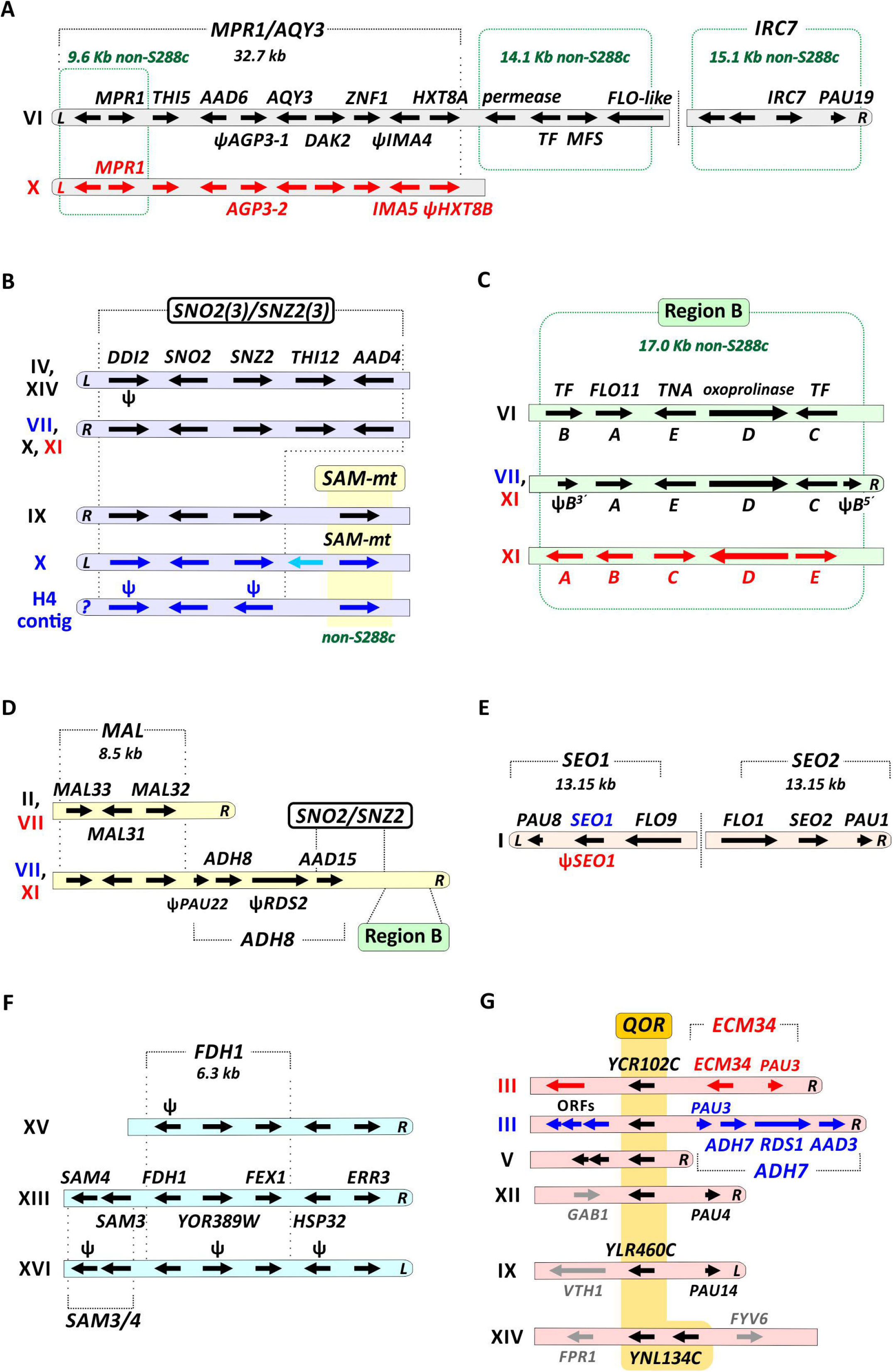
Structural representation of key regions amplified in H3 and H4 genomes. Gene-encoding regions that have increased copy numbers or are not found in S288c are shown. Structural dimensions of genes (arrows) and their chromosomal dispositions are schematic (i.e., not in scale). Chromosome numbers are shown on the left. Rounded chromosome tips indicate that the depicted region has a subtelomeric location at either the left (*L*) or right (*R*) arm of the chromosome; their absence indicates that the respective gene region lays at the chromosome core. Genes and their respective labels that are indicated in black represent alleles found in both H3 and H4. Red represents alleles found only in H3, and those in blue exclusively in H4. Putative pseudogenes (ψ) are also indicated. Non-S288c regions are framed by green dashed lines forming a rectangle. **(A)** The *MPR1/AQY3* region encompasses 10 genes within c. 32.7 kb. It is placed at the left subtelomere of Chr. VI (in H3 and H4) and on the Chr. X (only in H3). A 14.1 Kb non-S288c segment encoding a permease, a transcription factor (TF), a transporter of the major facilitator superfamily (MFS), and a FLO-like protein is located downstream of the *MPR1/AQY3* region. The *IRC7* region is at a distal position at the subtelomeric part of the Chr. VI right arm. **(B)** The *SNO2(3)/SNZ2(3)* cluster is a multi-copy region in the H3 and H4 genomes. On the Chr. IX and X of H4, *SNO2(3)*/*SNZ2(3)* is flanked by genes encoding *SAM-dependent methyltransferases* (*SAM-mt*, yellow), a non-S288c gene family ubiquitous in fuel ethanol strains. The same configuration is found in an unplaced (?) contig of H4. **(C)** The Region B is a mobile five-gene cluster. The gene order (A, B, C, D, E) of the cluster is shifted from chromosome to chromosome (including a split of gene B in H3 Chr. XI and in H4 Chr. VII), denoting differential integration events of a circular episome (Borneman et al., 2011). TF, transcription factor; TNA, putative high affinity nicotinic acid transporter. **(D)** The *MAL* cluster of about 8.5 kb has three copies in H3 and two in H4. The subtelomeric part at the right arm of the Chr. XI (in H3) and Chr. VII (in H4) harbors a multicluster segment encompassing the regions *MAL*, *ADH8*, *SNO2(3)*/*SNZ2(3)*, and Region B. **(E)** The *SEO1* and *SEO2* genes are part of an inverted repeats duplication of c. 13.15 kb found at the tips of the Chr. I left and right arms. **(F)** The *FDH1* cluster is triplicated on H3 and H4 genomes. The *SAM3* and *SAM4* genes are also duplicated. **(G)** The family of quinone oxidoreductase (QOR) encoding genes is amplified in H3 and H4. In both H3 and H4 the locus encoding the *YCR102C* homolog is triplicated (Chrs. III, V, and XII), while one copy related to *YLR460C* is represented on Chr. IX. *YNL134C* is duplicated in tandem on the Chr. XIV. In H4, the subtelomeric region of Chr. III right arm is syntenic to S288c and harbors a segmental duplication of the *ADH7* region [also present on the Chr. VII, panel (**D**)]. In contrast, in H3 a copy of the *ECM34* region is observed at the same part of Chr. III right-end.

### Non-S288c genetic features provide hallmarks of bioethanol strains

In search for genetic features that may be linked to bioethanol production, we first compared H3 and H4 genomes to the reference S288c to identify genes exclusive to the Brazilian strains. We detected 38 and 46 genes in H3 and H4, respectively, that are not present in S288c (**Supplementary Table 13**). Many of them are embedded within gene clusters (**Figure 1**, **Figure 2**, and **Supplementary Table 14**). A typical case of a non-S288c gene is provided by *MRP1* (encoding L-azetidine-2-carboxylic acid acetyltransferases) which is present in H3 and H4 (**Figure 1** and **2A**) and widespread among many yeasts (**Figure 3**). Another excellent example is the cluster of 14 genes (~34.7 kb) located in the H4 genome at the Chr. X left-end (**Figure 1**). This region is rare among *S. cerevisiae* representatives and has been previously observed in JAY291 (Argueso et al., 2009) and in a wild isolate from Costa Rica (Wohlbach et al., 2014). The cluster is also found in *Saccharomyces paradoxus* strains, thus the genes may have been introgressed into the PE-2 genome from this donor species (**Supplementary Figure 3, Supplementary Table 12**). Interestingly, the 34.7 Kb cluster is limited at the right side by the *OPT1* gene which in different strains represent distinct chimeras between the S288c *OPT1* gene and the *S. paradoxus* ortholog, suggesting a recombination hot spot (**Supplementary Figures 3** and **4**). It is worth noting that the presence of *S. paradoxus* related *OPT1* and its neighbor ORF *YJL213W* has been described in Brazilian natural populations of *S. cerevisiae*, while some cachaça strains also encode the introgressed *OPT1* (Barbosa et al., 2016; Barbosa et al., 2018).

**FIGURE 3.**
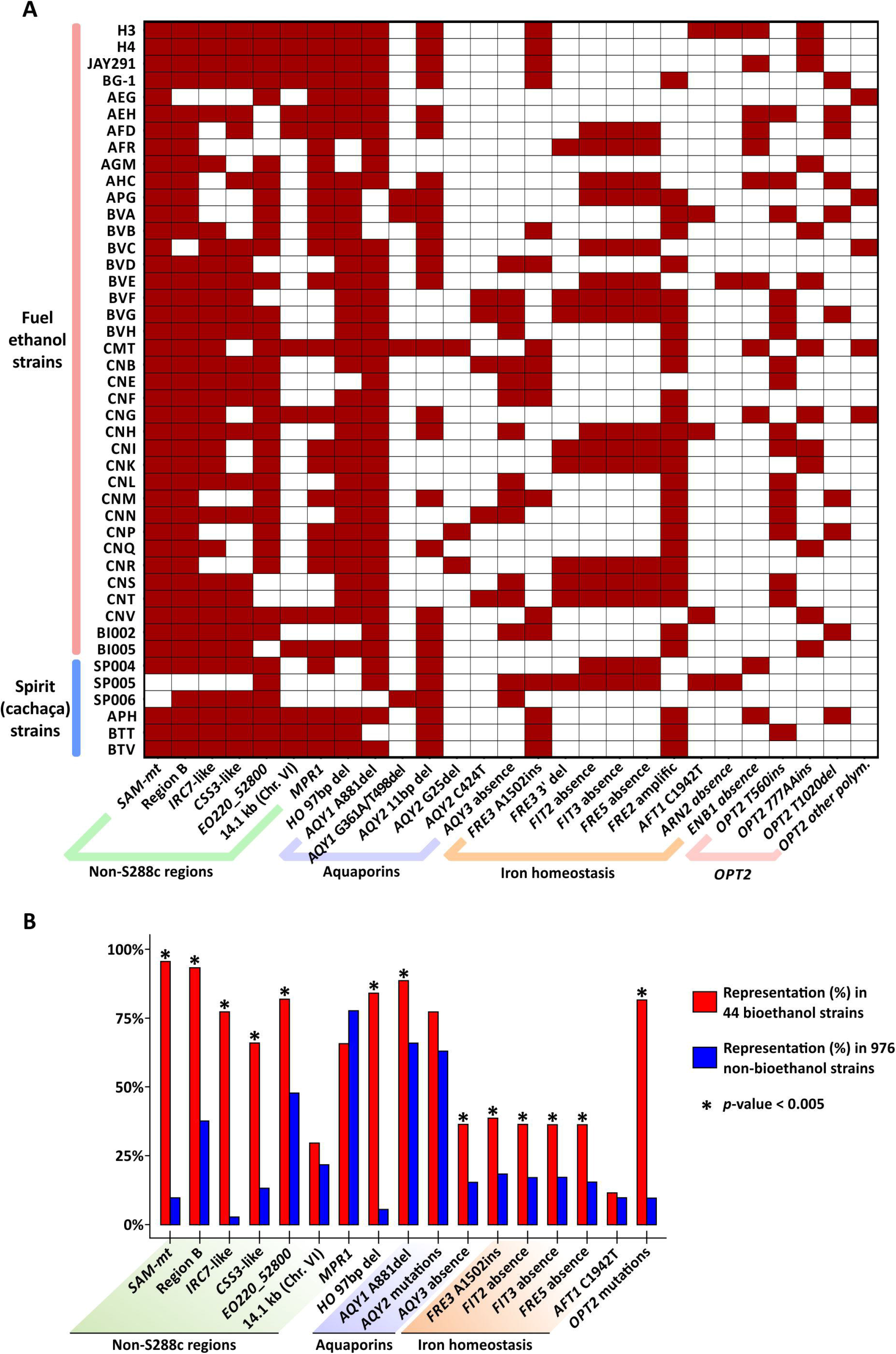
Genetic features enriched among yeasts of the bioethanol group. **(A)** The presence of key genetic features (represented on the x-axis by non-S288c genes/regions, loss-of-function alleles, and absence of specific genes) was determined in 44 strains (y-axis) by BLASTN searches. The color scheme depicts the presence (dark red) or absence (white) of the feature for each strain. *S. cerevisiae* isolates are described in the **Supplementary Table 1**. **(B)** Enrichment of each genetic feature in 44 strains from the bioethanol group (percentage of representation shown in red bars). For comparison, BLASTN searches were also conducted against a dataset of 976 strains and the fraction of presence for each genetic feature is shown in blue bars. *AQY2* and *OPT2* mutations combine the polymorphisms shown in panel (**A**). Asterisks (*) indicate significant enrichment of the feature among yeasts of the bioethanol group (Fisher’s exact test, *p* < 0.005).

Standing out within non-S288c gene families is the one encoding for SAM-dependent methyltransferases (SAM-mt) that is found in H3 and H4 in one and three copies, respectively (**Figure 1**, **Figure 2B, Supplementary Tables 13** and **14**). This gene family is rare among *S. cerevisiae* and uncharacterized so far. Using BLASTN we assessed if *SAM-mts* and six other non-S288c genes/regions found in H3 and H4 genomes (green regions on chromosomes of **Figure 1**) are common among yeasts of the Brazilian bioethanol group (including 38 fuel-producing yeasts plus six cachaça isolates, listed on the **Supplementary Table 1**). For comparison with the Bioethanol group, we extended our BLASTN searches to interrogate a cluster of 976 genomes comprising 34 non-bioethanol clades from the 1,002 Yeast Genomes Project (see **Supplementary Figure 5**). As a result, we found that the *SAM-mt* gene is present in 42 out of 44 yeasts (i.e., 95.45% of taxa) from the Brazilian bioethanol group, including all biofuel strains (it is only absent from two cachaça strains) (**Figure 3A** and **Figure 3B**). Conversely, *SAM-mt* genes are observed in only 95 out of 976 remaining non-bioethanol yeasts (i.e., 9.73% of the strains in the dataset) (**Figure 3B** and **Supplementary Figure 5**). This significant enrichment (*p* < 0.005, Fisher’s exact test) among the Brazilian strains makes the presence of *SAM-mt* a hallmark of bioethanol yeasts.

The Region B (**Figure 1** and **Figure 2C**) encompasses five non-S288c genes that represent a signature of horizontal gene transfer from *Zygosaccharomyces bailii* to *S. cerevisiae*, postulated to have occurred at the basal lineage of the wine yeast clade (Borneman et al., 2011; Almeida et al., 2015). Region B is also enriched (*p* < 0.005) within the group of Brazilian bioethanol yeasts, found in 41 (93.2%) of 44 strains, while only 365 (37.4%) of the 976 remaining (non-bioethanol) strains display the genetic cluster (**Figure 3A** and **3B**, **Supplementary Figure 5**). Another case of non-S288c gene cluster is the previously described (Argueso et al., 2009) four-gene region of about 14.1 Kb, which is located on the Chr. VI, just downstream the *AQY3* cluster (**Figure 1** and **Figure 2A**). This region is not particularly represented within the bioethanol group (29.5%) when compared to the rest of the yeasts (21.8 %) (**Figure 3A** and **3B**). Finally, the annotated ORF *EO220 52800* is also absent from S288c, but present in 36 taxa (82.0%) from the Brazilian bioethanol group and in 464 (47,5%) non-bioethanol strains. This gene is 99% identical to a *S. paradoxus* ortholog and represents a potential case of introgression, enriched (*p* < 0.005) within the bioethanol group.

There are other chromosomal regions in H3 and H4 that slightly differ from S288c (below 93% nucleotide identity), and are instead more similar to regions from *S. paradoxus*. These potentially introgressed regions are considered here as non-S288c features and were also selected for analysis by BLASTN. In H3 and H4, a four-gene region spanning ~15.1 Kb, distally placed on Chr. VI right-end (**Figure 1** and **Figure 2A**) carries an ortholog of the S288c *IRC7* that encodes a beta-lyase involved in the production of thiols (Roncoroni et al., 2011). The Chr. VI-encoded *IRC7* from H3 and H4 has 82.3% nucleotide identity to the S288c counterpart, while carrying 99.8% identity to the *S. paradoxus* homolog. We found that the *S. paradoxus IRC7*-type is particularly associated (*p* < 0.005) to the bioethanol group [present in 34 (77.3%) taxa] when compared to non-bioethanol yeasts [only in 26 (2.66%) strains] (**Figures 3A and 3B**). Some bioethanol strains have both the S288c and *S. paradoxus IRC7* paralogs. As recently reported (D’Angiolo et al., 2020), we observed that the *S. paradoxus IRC7* is also enriched in the Mexican Agave and French Guiana Human populations (**Supplementary Figure 5**). Another case is provided by a ~5.35 Kb region on Chr. II (**Figure 1**) of H3 and H4 that harbors the *CSS3* allele. This gene shares 99.8% identity with the *S. paradoxus* homolog, while having 92.4% identity to the S288c ortholog. The genome of H4 also encodes a paralog on Chr. XV that shares 98.9% identity with the homolog in S288c (**Figure 1**). The *S. paradoxus CSS3*-type is enriched (*p* < 0.005) within the bioethanol group [29 strains (65.9%)] when compared to the remaining yeasts of the non-bioethanol dataset [129 strains (13.2%)] (**Figure 3A and B, Supplementary Figure 5**) and has also been previously observed in cachaça strains of the C1 and C2 clades (Barbosa et al., 2018). Taking together, the presence of the Region B, *SAM-mts, EO220 52800*, and the *S. paradoxus IRC7*- and *CSS3*-types represent distinctive features associated with most Brazilian bioethanol yeasts.

### Possible fingerprints of domestication in bioethanol strains: aquaporins and iron homeostasis related genes

We further compared H3 and H4 genomes against the reference S288c to investigate cases of gene loss or defective alleles associated with bioethanol strains. Based on mapping of short reads, H3 and H4 lack 49 and 35 genes, respectively, that are encoded in the reference genome of S288c (**Supplementary Table 15**). Most of these genes correspond to subtelomeric gene clusters that are duplicated in S288c, but are completely absent or occur in single copies in H3 and/or H4. Some examples are the S288c regions *MPH2/SOR2* (four genes on Chr. IV), *VTH2/MPH3* (five genes on Chr. X), *BDS1* (five genes on Chr. XV), and *ENB1* (six genes on Chr. XV). Read depth analysis of the CNVs across multiple *S. cerevisiae* strains indicates that those regions are also underrepresented among non-bioethanol yeasts (**Supplementary Figure 6**), suggesting that S288c genes not found at the syntenic position on H3 and H4 chromosomes may have arisen from gene gains or duplications specific to the S288c lineage.

We searched for gene models of H3 and H4 with ORFs shorter than their corresponding syntenic homologs in S288c, i.e., potentially having premature stop codons, as a proxy for loss of function. Using this approach, we identified 44 and 40 potential pseudogenes in the H3 and H4 genomes, respectively (**Supplementary Table 16**). Interestingly, we observed many instances of premature stop codons in genes within amplified regions (e.g., *DDI2(3), SNZ2(3), AGP3, SEO1*, and other examples on **Figure 2**), suggesting that the mutated gene is not the one under selection for amplification. Among the remaining putative defective alleles, some may be implicated in adaptation to industrial conditions. For example, most bioethanol strains have been described as heterothallic, as haploid cells cannot switch their mating-type (Della-Bianca et al., 2013). Heterothallism may be advantageous in industrial yeasts to preserve genetic diversity within heterozygous diploid genomes (Argueso et al., 2009). Its genetic basis largely relies on inactivation of the gene encoding HO endonuclease that regulates recombination within the *MAT* locus, leading to mating-type switching (Steensels et al., 2014). The *HO* gene is defective in *S. cerevisiae* PE-2 due to a 97 bp deletion within the region that specifies the DNA-binding domain of the encoded protein (Argueso et al., 2009). We confirmed the presence of this *ho* allele in 37 (84.1%) of 44 Brazilian bioethanol strains, while only 54 (5.53%) of 976 non-bioethanol strains display a similar 97 bp deletion (**Figure 3A** and **3B**). Interestingly, many bioethanol strains are heterozygous with the mutated *ho* allele coexisting with the native form. Nonetheless, the presence of the 97 bp deletion within *HO* in 84.1% of the strains from the Brazilian bioethanol group (including three cachaça strains) is a conserved synapomorphy, indicating that the group consists of a uniform pool of phylogenetic related strains.

Mutations that inactivate aquaporin genes are adaptive for fermentation of sugar-rich substrates in industrial processes because the loss-of-function alleles confer a significant selective advantage to withstand high osmolarity (Bonhivers et al., 1998). Accordingly, defective aquaporin *AQY1* and *AQY2* genes represent a hallmark of domestication among European wine and Brazilian cachaça strains, while both genes remain functional in the related wild Mediterranean and Brazilian *S. cerevisiae*, respectively (Goncalves et al., 2016; Barbosa et al., 2018). The mutation in *AQY1* that is frequently associated with wine and cachaça populations C1 and C2 is a deletion of adenine at position 881 (A881del) (Barbosa et al., 2018). This polymorphism is present in 39 (88.6%) of strains in the Brazilian bioethanol group (**Figure 3A** and **3B**). Two additional bioethanol strains display an alternative substitution (G361A) in *AQY1* leading to a valine-to-methionine (V121M) exchange known to disrupt water transport activity of the encoded protein (Bonhivers et al., 1998), whereas one cachaça strain has a deletion (T498del) causing a frameshift. In total, 42 (95.5%) of 44 strains from the Brazilian bioethanol group have a defective *AQY1* allele. For *AQY2* we observed three frequent loss-of-function alleles affecting 34 (77.3%) strains (**Figure 3**). These include the 11bp deletion, typical of wine and cachaça strains (Goncalves et al., 2016; Barbosa et al., 2018), a guanine deletion at position 25 (G25del), and a premature stop codon at position 424 (C424T). *AQY3* encodes yet a third yeast aquaporin implicated in the regulation of vacuole volume and osmotic responses during hypertonic stress (Patel, 2013). Convergent to the trend of loss-of-function alleles in aquaporins among bioethanol strains, we observed that *AQY3* is missing from the genome of over a third of strains from the Brazilian bioethanol group (36.4%, **Figure 3**), a significant (*p* < 0.005) enrichment compared to the loss of *AQY3* among only 15.4% of the remaining non-bioethanol yeasts (**Figure 3B**). Altogether, the 44 analyzed strains of the bioethanol group have at least one inactive aquaporin allele, 38 have two, and ten have three.

We observed many cases of defective or deleted alleles related to iron homeostasis. Ferric reductases (Fre) act at the cell periphery by reducing soluble or siderophore-bound ferric iron allowing the uptake of the ferrous form through the Fet3/Ftr1 complex (Philpott et al., 2002). We found in H3 and H4 an adenine insertion (frameshift) at position 1502 of the *FRE3* gene that is also enriched among 17 (38.6%) strains of the Brazilian bioethanol group when compared to 178 (18.2%) of non-bioethanol yeasts (*p* < 0.005; **Figure 3**). The *FRE3* gene is located within a cluster of iron regulated genes on the Chr. XV right-end that also harbors the genes *FIT2* and *FIT3* (encoding for mannoproteins that promote retention of siderophore-iron in the cell wall) and the ferric reductase homologue *FRE5*. We observed two types of deletion encompassing *FIT2*, *FIT3* and *FRE5* genes in the genomes of bioethanol strains (**Supplementary Figure 7**). A deletion of ~5.36 Kb is present in 10 strains and has breaking points surrounding the three genes. Interestingly, a second larger deletion that is found in 22 bioethanol strains is reminiscent of the ~14 kb deletion described in flor yeasts (Eldarov et al., 2018). We found that the complete deletion of *FIT2, FIT3* and *FRE5* is enriched (*p* < 0.005) among the Brazilian bioethanol group [15 strains (36.4%)] with respect to non-bioethanol yeasts [161 taxa (16.5%)] (**Figure 3B**). This three-gene deletion was identified *en bloc* among many yeast clades, such as Ale Beer, Wine European Subclass IV, French Guiana Human, African Beer, French Dairy, Alpechin, Mixed Origin, and some Mosaic groups (**Supplementary Figure 5**). The *FIT/FRE* deletion, together with a premature stop codon at position 1942 (C1942T) of the gene encoding the iron-responsive transcriptional factor Aft1, has been associated to flor yeasts and other *S. cerevisiae* strains that are proficient in iron uptake, and thus sensitive to iron when the metal ion is abundant in the environment (Eldarov et al., 2019). The same C1942T mutation in *AFT1* is observed in H3 and in three other bioethanol strains, while the BVA isolate has a premature stop codon at position 1855 (C1855T) (**Figure 3A**). H3 genome also displays a complete deletion of *ARN2* and *ENB1* that encode for iron-siderophores transporters (Philpott et al., 2002). Finally, based on read depth analysis (see next section), we detected an increase in *FRE2* copy number among 28 bioethanol strains (**Figures 3A** and **Figure 4**). This resulted from a ~26.5 kb amplification encompassing seven genes at the Chr. XI left-end (**Supplementary Figure 8**).

**FIGURE 4.**
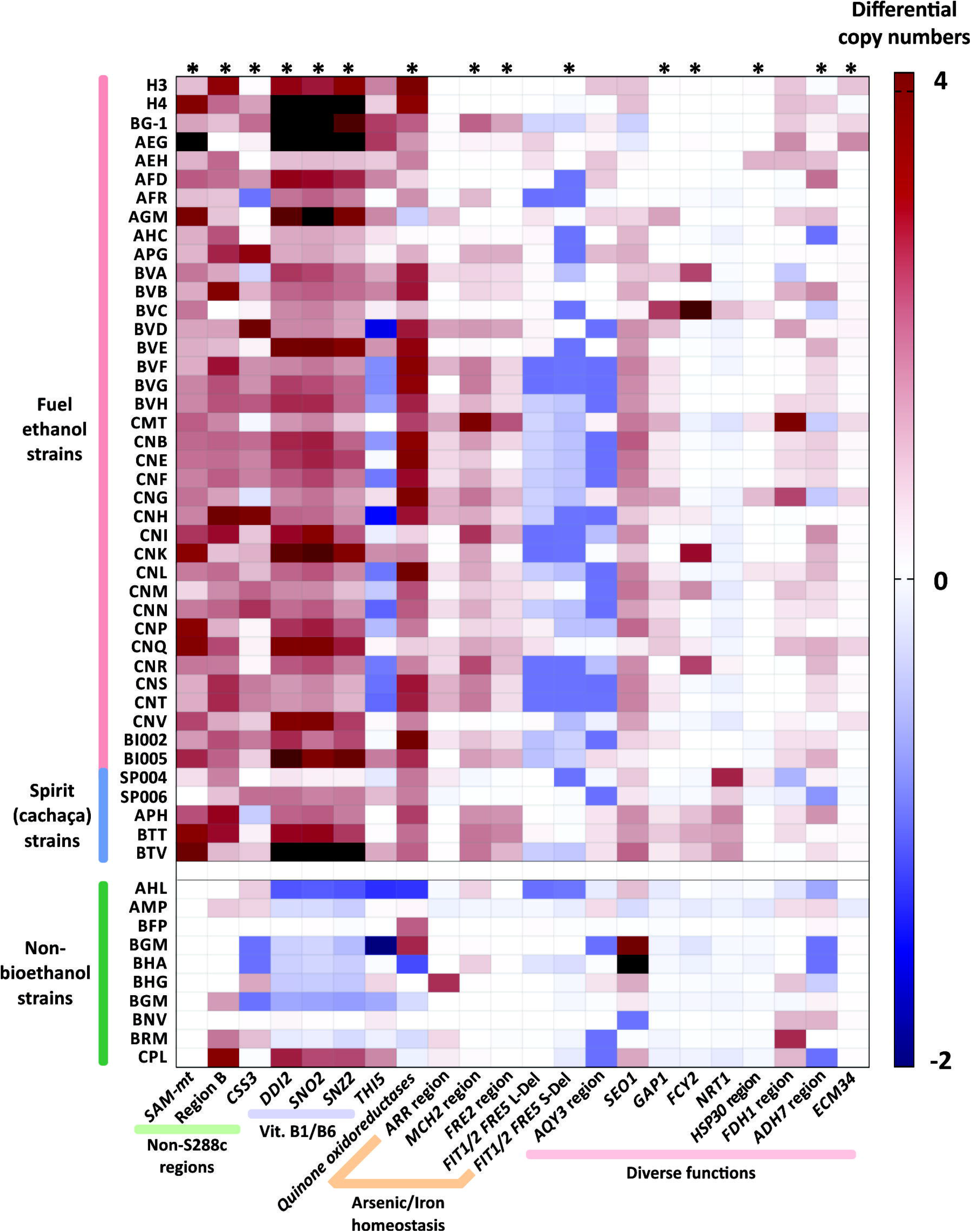
Differential copy numbers of genes/regions in strains of the bioethanol group compared to S288c. Copy numbers of key genes/regions (labeled at the bottom) were measured in bioethanol yeasts by read depth based on Illumina sequence data. The estimated CNVs of probed genes/regions were discounted from the expected copy numbers according to the S288c genome annotation. A heatmap is shown depicting a color scheme in which extra copies in strains of the bioethanol group are quantitatively expressed in a red gradient (up to four copies). Equal number of copies found in both bioethanol and S288c strains is represented in white, while surplus of copy numbers in S288c is displayed as a blue gradient. Note that in some cases, hemizygous patterns account for non-integer gene copy numbers, since multiple diploid genomes are represented within the dataset (Peter et al., 2018). A group of ten randomly chosen non-bioethanol yeasts were included as external controls. Dark red colored boxes outside the red gradient range represent cases in which more than four extra copies of the probed gene/region were counted. Significant enrichment of CNV among the bioethanol clade is indicated with an asterisk (*) above the respective column (*p* < 0.05, Mann-Whitney U test).

*OPT2* (encoding the oligopeptide transporter 2) has been demonstrated to have genetic interactions with components of the iron homeostasis, such as *FET3*, *SIT1*, *GEF1*, and *FRE3* (Elbaz-Alon et al., 2014; Costanzo et al., 2016). Accordingly, the Opt2 protein is involved in the regulation of the peroxisomal, mitochondrial, and cytosolic glutathione redox homeostasis (Elbaz-Alon et al., 2014). We observed three major polymorphisms leading to the loss of function of *OPT2* that are enriched among strains of the bioethanol group (**Figure 3A**). They are T560ins (17 strains), AA777ins (13 yeasts) and T1020del (10 strains). In addition, other mutations (A41del, C1224A, and two 17-bp insertions at positions 1499 and 1519, respectively) affect five additional strains. Altogether, 36 Brazilian bioethanol yeasts (81.8%) have an inactive *OPT2* allele (**Figure 3B**). By searching for the same polymorphisms across the dataset of 976 non-bioethanol yeasts, we estimated that only 9.5% of them carry identical mutations to those found in the Brazilian strains (**Figure 3B**). Although it remains unclear how the presence of these polymorphisms in *OPT2* and in iron homeostasis components impacts yeast physiology, the recurrent incidence of several genetic features shared among bioethanol yeasts suggests common signatures of adaptation to conditions related to the industrial production of ethanol from sugarcane.

### Distinct patterns of gene CNVs unite bioethanol yeasts

Many cases of gene CNVs in H3 and/or H4 genomes are evident relative to the S288c reference (**Figure 1** and **2, Supplementary Tables 14, 15**, and **17**). We further assessed if these CNVs are also observed in other bioethanol yeasts (**Supplementary Table 4**). Genome data (Illumina short reads) from 42 bioethanol strains (including H3 and H4) were independently mapped against a concatenated DNA sequence in which genes/regions probed for CNVs are represented (**Supplementary Sequence File 1**, see Materials and Methods). For each strain, read depth within the analyzed sequence regions was normalized against the read depth over the genome as background (**Supplementary Table 4**), allowing estimation of differential CNVs relative to the S288c genome (**Figure 4** and **Supplementary Figure 6**). CNVs for 10 non-bioethanol strains were also analyzed as external controls. Our analysis revealed that several cases of gene/region CNVs are enriched in the Brazilian bioethanol group (*p* < 0.05, Mann-Whitney U test; **Figure 4**). Remarkably, all bioethanol yeasts have increased copy numbers within the gene cluster for vitamin B1/B6 biosynthesis; the core genes of this cluster comprise *SNO2(3)* and *SNZ2(3)*, and are usually flanked by *DDI2(3), THI5(12), and AAD4*. Based on molecular karyotyping and southern blot hybridization, Argueso et al. (2009) reported four copies of *SNO2(3)*/*SNZ2(3)* in the haploid JAY291 and up to nine copies in its parental diploid. Contrastingly, genome assemblies of H3 and H4 resolved five and seven *SNO2(3)*/*SNZ2(3)* copies, respectively, located at the termini of different chromosomes (**Figure 1** and **Figure 2B, Supplementary Tables 14** and **17**). It is worth noting that CNVs estimations from read depth for the *SNO2(3)*/*SNZ2(3)* cluster and other genes/regions correlate to CNVs observed in our two genome assemblies (**Supplementary Figure 1**; Pearson’s *r* = 0.9976 and 0.9978 for H3 and H4, respectively).

Besides the *SNO2(3)/SNZ2(3)* amplification, the occurrence of other gene CNVs in bioethanol yeasts has been described in previous comparative genomic hybridization analyses (Argueso et al., 2009; Stambuk et al., 2009; Babrzadeh et al., 2012). This is the case of *SEO1*, encoding an allantoate transporter, which is embedded within a ~13.0 Kb inverted duplication on the left- and right-ends of Chr. I (**Figure 1** and **Figure 2E**). This segmental amplification involving *SEO1* appears to be common among bioethanol and non-bioethanol strains (**Figure 4**). Another example is the previously reported *SAM3/SAM4* amplification (Argueso et al., 2009) that was resolved in our H3 and H4 genome assemblies into two copies: one located at the Chr. XIII right-end, and the other at Chr. XVI left-end (**Figures 1** and **2F**). However, we noticed that two copies of *SAM3*/*4* are normal among various *S. cerevisiae* strains (**Supplementary Figure 6**), therefore a single copy in the S288c genome may indicate a lineage-specific gene loss. We observed that the occurrence of many other gene CNVs is not exclusive to bioethanol strains, including gene clusters such as *MAL1x-3x, ENA1/2/5* and *MPR1* (**Supplementary Figure 6**).

Among newly found cases of CNVs in the H3 and H4 genomes, the *CUP1* gene (encoding for a metallothionein) stands out by occurring in tandem repeats with ~18 estimated copies on Chr. VIII of both H3 and H4 (**Supplementary Table 17**, **Supplementary Figures 9** and **10**). *CUP1* is known as a hotspot for adaptive and non-adaptive amplifications on *S. cerevisiae* genomes (Zhao et al., 2014), thus our observation of variable copy numbers of *CUP1* among bioethanol and non-bioethanol yeasts (**Supplementary Figure 10**) is not surprising. Our gene CNVs analysis also confirmed that some of the non-S288c genes, such as the Region B, *SAM-mt*, and *CSS3*, often display more than one copy in strains from the Brazilian bioethanol group (**Figure 4**). For example, the AEG strain has four estimated copies of *SAM-mt* genes, while H4 has three paralogs. Likewise, Region B exists in three copies in H3, two located in the middle of Chrs. VI and XI, respectively, and one at the right tip of Chr. XI. H4 has a copy of Region B syntenic to H3 on the core of Chr. VI, and another at the subtelomeric part of Chr. VII right-end (**Figures 1** and **2C**). These subtelomeric copies of Region B at the Chr. VII of H4 and Chr. XI of H3 are in fact part of a large (~48.9 kb) cluster that also includes the regions *MAL, ADH7(8)*, and *SNO2(3)/SNZ2(3)* (**Figures 1** and **2D**). In H4, the *ADH7(8)* region (*PAU3/ADH7/RDS1/AAD3)* is also duplicated at the Chr. III right-end, while H3 displays at this corresponding position an extra copy of *ECM34* (representing a fusion of the S288c *ECM34* and *YHL042W* homologs) (**Figures 1** and **2G**). We found that amplifications of *ECM34* and the *ADH7* region appear to be recurrent among bioethanol strains (**Figure 4**). Similarly, the *FDH1* region (comprising *FDH1 /YOR389W/FEX1*, **Figures 1** and **2F**) occurs in three copies in the genomes of H3, H4, and other bioethanol strains, and only two in the S288c genome.

Previous comparative genomic hybridization analysis of the strain CAT-1 (Babrzadeh et al., 2012) and current read mapping profiles of bioethanol yeasts other than H3 and H4 reveal further genomic regions with differential CNVs in comparison to S288c. These include the earlier mentioned loci involved in iron homeostasis, such as the *FRE2* cluster (**Supplementary Figure 8**) that is amplified in 27 strains of the bioethanol group, and partial or full deletions of the *FIT1/2/FRE5* region (**Supplementary Figure 7**) in 30 taxa (**Figure 4**). Alongside the *FRE2* amplification, the adjacent *MCH2* region containing the genes *YKL222C* (unknown function) and *MCH2* (specifying a monocarboxylate permease) also shows an increased copy number in many bioethanol yeasts (**Figure 4** and **Supplementary Figure 8**). At the subtelomere of Chr. XVI right-end (**Supplementary Figure 11A**), a ~19 kb-region containing the *ARR1* (encoding a transcriptional activator responsive to arsenic), *ARR2* (arsenate reductase), and *ARR3* (plasma membrane transporter required for detoxification of arsenic compounds) is amplified in 20 strains of the Brazilian bioethanol group (**Figure 4**). CNVs in regions encoding components of arsenic and iron homeostasis suggest that the regulation of mineral nutrition and/or toxicity may be under selection in bioethanol yeasts. Other genes encoding functions related to nutrient acquisition are also found to be amplified among bioethanol yeasts (**Supplementary Figure 12**). *GAP1* (encoding the general amino acid permease) is amplified among 24 strains of the bioethanol group, while *FCY2* (purine-cytosine permease) has multiple copies in some strains. Interestingly, amplification of *NRT1* (transporter for nicotinamide riboside, a NAD^+^ precursor) is a unifying feature of cachaça strains. Finally, lower copy-numbers implicating the *AQY3* region (**Supplementary Figure 11B**) and the amplification of the *HSP30/PMP1* cluster (**Supplementary Figure 11C**) are observed in 18 and 6 strains, respectively, from the Brazilian bioethanol group. These CNVs might be related to stress responses, as *AQY3* deletions potentially counterbalance high osmolarity (Patel, 2013), and overexpression of *HSP30* confers thermotolerance to yeasts (Meena et al., 2011).

Of special interest is the amplification in H3 and H4 genomes of gene homologs annotated as *quinone oxidoreductases* (**Figures 1** and **2G**). They are represented in S288c as *YCR102C* (Chr. III), *YLR460C* (Chr. XII) and *YNL134C* (Chr. XIV), where they are observed as monoallelic. In contrast to S288c, H3 and H4 have three homologs related to *YCR102C* on subtelomeric regions of Chrs. III, V, and XII, and a further paralog more similar to *YLR460C* at the Chr. IX left-end. In addition, H3 and H4 display a tandem duplication of *YNL134C* at the core of Chr. X (**Figures 1** and **2G**), also previously observed in the strain CAT-1 (Babrzadeh et al., 2012). Amplifications of these oxidoreductases are widespread among yeasts in the Brazilian bioethanol group (with the exemption of AGM and AHC strains) (**Figure 4**). Therefore, the functional analysis of this amplified gene family may be important to elucidate molecular mechanisms that underpin stress tolerance in bioethanol yeasts.

### Whole-genome phylogeny reveals a single origin of bioethanol yeasts

To assess the evolutionary origins of bioethanol yeasts in Brazil, we reconstructed a phylogenetic tree using an alignment-free (AF) approach, based on whole-genome sequences from 197 broadly sampled, representative yeast genomes (see Materials and Methods for details). This AF approach, based on analysis of short, sub-sequences of length *k* (*k*-mers), represents a faster, more-scalable alternative to standard phylogenetic inference based on multiple sequence alignment. Bypassing the unrealistic assumption of full-length contiguity among homologous sequences, and the computationally demanding step of multiple sequence alignment, AF approach allows for inferring phylogenetic relationships using whole-genome sequences quickly. Such an approach is known to be robust to distinct evolutionary scenarios, e.g. genetic rearrangement (Bernard et al. 2016; Bernard et al. 2019), and has been used to accurately and quickly infer phylogenetic relationship in thousands of microbial genomes (Bernard et al. 2018; Zielezinski et al. 2019). We calculated the pairwise distances among the 197 genome sequences based on *k*-mers at *k* = 21, from which a neighbor-joining tree is derived. This tree (**Figure 5**) displays a very similar topology to the maximum likelihood tree reconstructed from codon alignments of 2,020 concatenated single-copy nuclear genes (Gallone et al. 2016). We recovered five major clades, consistent with previously reported result (Gallone et al., 2016): Wine (23 taxa), Beer 2 (22 taxa), Mixed (19 taxa), Asia (9 taxa), and Beer 1 (68 taxa) containing Britain (26 taxa), US (10 taxa), and Belgium/Germany (22 taxa) (**Figure 5**). All bioethanol yeasts were grouped in a monophyletic clade in association to the wine supergroup (**Figure 5**). Our result, based on comprehensive wholegenome sequences, provides unequivocal support for a common single origin for bioethanol strains in Brazil, including the cachaça yeasts and isolates from biofuel production plants.

**FIGURE 5.**
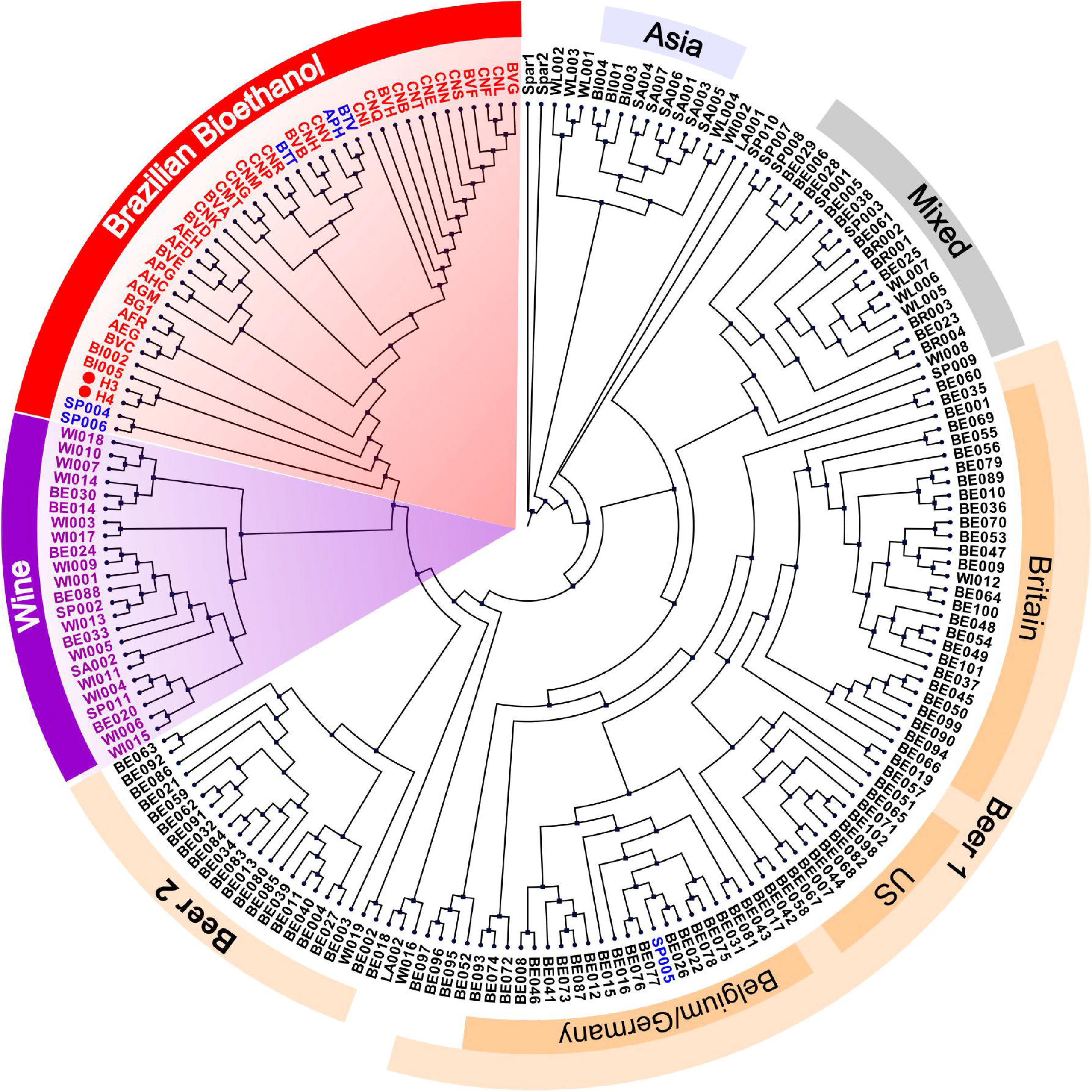
AF-based phylogenetic tree of 197 yeasts genomes. A distance matrix was calculated based on an AF approach and used to build a NJ phylogenetic tree. The Brazilian bioethanol group (red) was clustered in a single clade, with the exemption of the cachaça strain SP005. In addition, five major clades were resolved in accordance to Gallone et al. (2016) and are indicated with different colors. H3 and H4 are indicated with red circles. Cachaça strains are labeled in blue.

## DISCUSSION

### Genetic signatures of bioethanol yeasts and adaptation to industrial fermentation

The present comparative genomics study one of the first attempts to analyze genome structure, gene content, polymorphisms, gene CNVs, and phylogenetic relationships across diverse *S. cerevisiae* used for bioethanol production in Brazil. The Brazilian fuel ethanol production from sugarcane substrates represents a peculiar bioprocess in which yeast cells are recycled and reused in successive fermentation batches. Therefore, throughout the year, bioethanol yeasts are steadily challenged by multiple biotic and abiotic stresses (Basso et al., 2011; Della-Bianca et al., 2013). Little is known about the remarkable genetic features that allow these strains to withstand such pressures while maintaining an excellent fermentation performance. The elucidation of the genomic sequences from the bioethanol strains CAT-1, BG-1, and PE-2 (including JAY291, and our H3 and H4 assemblies) revealed highly heterozygous diploid genomes carrying a rich source of adaptive genetic diversity (Argueso et al., 2009; Babrzadeh et al., 2012; Coutoune et al., 2017). It is suggested that a homeostatic balance between selective pressures in the complex environment of the distillery tends to maintain the heterozygous state of the genome (Sampaio et al., 2019). However, under prevalence of particular selective pressures, mitotic recombination events may lead to allelic loss-of-heterozygosity resulting in the fixation of adaptive haplotypes (Rodrigues-Prause et al., 2018; Sampaio et al., 2019). Another important repository of genetic variability lies within subtelomeric regions (Argueso et al., 2009). The patterns of segmental amplifications or deletions observed here in subtelomeric gene clusters of bioethanol strains are in accordance with the notion that ectopic recombination events at the chromosome tips modulate the dosage of stress and nutrient related genes (Argueso et al., 2009; Babrzadeh et al., 2012). Overall, cryptic genetic variability was consistently observed throughout our survey of 44 bioethanol strains, where potential adaptive SNPs and gene CNVs were often observed in heterozygous and hemizygous states, respectively. More important is that, by scrutinizing such genetic diversity, we uncovered typical genetic features enriched (*p* < 0.05) in Brazilian bioethanol yeasts that comprise: (i) non-S288c marker genes, (ii) regions introgressed from *S. paradoxus*, (iii) synapomorphic deletions, (iv) polymorphisms related to adaptive alleles, and (v) prevalent gene CNVs at subtelomeric regions (**Figure 6**). The identification of specific genetic markers in bioethanol yeasts raises the question of whether these features represent adaptations to industrial fermentation.

**FIGURE 6.**
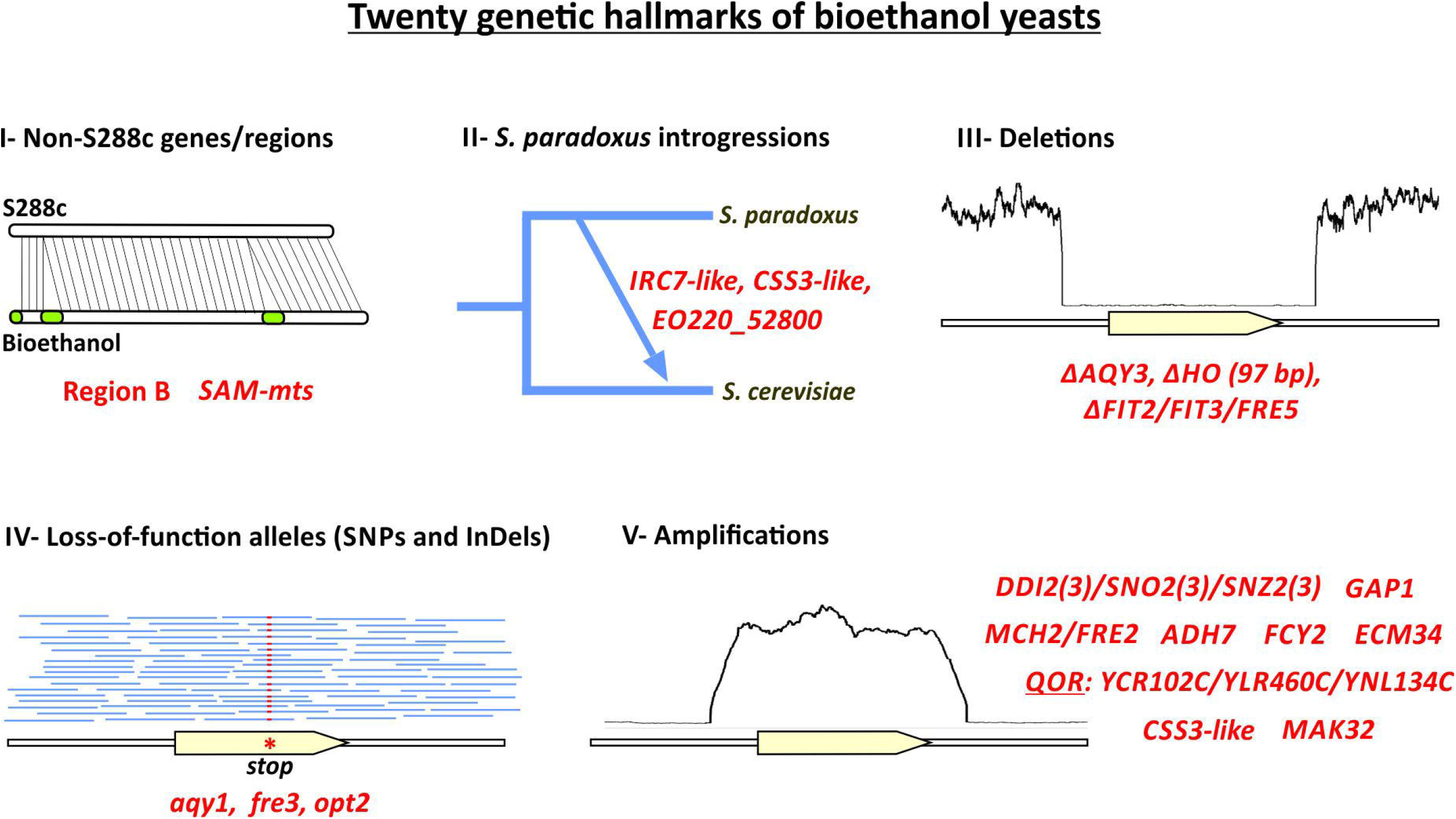
Twenty genetic hallmarks of Brazilian bioethanol yeasts. Genetic features enriched (*p* < 0.05) in bioethanol yeasts are shown in red, including: (i) non-S288c genes, (ii) putative introgressions from *S. paradoxus*, (iii) large deletions, (iv) SNPs and small InDels, (v) amplified regions.

In Brazil, ethanol generation from sugarcane is traditionally coupled with sugar production (Basso et al., 2011; Basso and Lino, 2019). In this industry, after sugarcane stalks are crunched, the resulting juice is concentrated to yield sugar crystals. The by-product of this operational step is the molasse, a remaining viscous phase containing about 45-60% sucrose and 5-20% of glucose/fructose (Basso et al., 2011; Della-Bianca et al., 2013). The actual substrate for yeast fermentation during ethanol production is usually a mixture of molasse with sugarcane juice (or water) in varied proportions. Molasses typically concentrate salts of potassium, calcium and magnesium in amounts high enough to trigger osmotic stress responses in fermenting yeasts (Basso et al., 2011). Our discovery of defective *AQY1/AQY2* alleles and deletions of *AQY3* seems to support this notion. Likely, loss of aquaporins is advantageous to counteract osmotic pressures related not only to sugar concentration, but in most part to the high salinity found in sugarcane molasses (Della-Bianca et al., 2013; Walker and Basso, 2020). Additional types of fermentation inhibitors are also common in molasses. Heating steps (up to 105°C) during sugarcane juice pretreatment can convert sugars into furfurals, formaldehyde, and toxic browning compounds (resulting from reactions between amino acids and reducing sugars) (Basso and Lino, 2019). Other inhibitory compounds, such as phenols from sugarcane plants, and pesticides and fertilizers used in the crop fields, can also be carried over into molasses (Basso et al., 2011). We found that CNVs of some genes involved in detoxification are overrepresented in bioethanol yeasts. The amplification of *DDI2(3)* paralogs occurs in the majority of bioethanol strains as part of the *SNO2(3)*/*SNZ2(3)* cluster. *DDI2(3)* encodes for a hydratase that metabolizes cyanamide, a mildly toxic compound used as a fertilizer, potentially generating pyrimidine intermediates for the synthesis of thiamin (Li et al., 2015). Other detoxification enzymes are encoded by *FDH1* (NAD^+^-dependent formate dehydrogenase), and *ADH7/AAD3* (encoding cinnamyl and aryl alcohol dehydrogenases, respectively). *ADH7/AAD3* are part of a segmental duplication in H4 and other bioethanol strains. In particular, increased expression of *ADH7* conferred furfural resistance in experimental evolution populations of yeasts (Heer et al., 2009). Tolerance to toxic aldehyde compounds has also been linked to the *YNL134C* and *YCR102C* functions, and overexpression of *YCR102C* is known to improve alcoholic fermentation under high acetic acid conditions (Zhao et al., 2015; Chen et al., 2019). These genes, together with *YLR460C*, comprise a family of annotated *quinone oxidoreductases* that is amplified in most bioethanol strains.

Nutrient availability seems to be a key factor for industrial fermentation of sugarcane substrates. In most *S. cerevisiae* strains there are three copies for both *SNO* (encoding glutaminase) and *SNZ* (PLP synthase). These two enzymes respectively control successive catalytic steps for the synthesis of vitamin B6 (PLP) (Rodriguez-Navarro et al., 2002). *SNO2(3)/SNZ2(3)* are located in subtelomeric clusters together with the thiamin biosynthetic gene *THI5(12)*, and their products physically interact in a complex that likely channels the PLP for thiamin production (Rodriguez-Navarro et al., 2002; Paxhia and Downs, 2019). In accordance with previous observations (Argueso et al., 2009; Stambuk et al., 2009), our analysis of gene CNVs revealed that all strains in the bioethanol dataset (exempting one cachaça strain) have increased copies of the *SNO2(3)/SNZ2(3)* region when compared with S288c. This suggests a special need for synthesis of vitamin B1 during industrial alcoholic fermentation. Such a demand may reflect the strict thiamin diphosphate (ThDP) requirement for the committed enzymatic step in the ethanol production; i.e., the conversion of pyruvate to acetaldehyde, controlled by different ThDP-dependent pyruvate decarboxylases in the yeast cell (Brion et al., 2014). We also found in strains of the Brazilian bioethanol group increased copy numbers of the general amino acid permease (*GAP1*), the purine-cytosine permease (*FCY2*), and, in cachaça strains, the transporter for nicotinamide riboside (*NRT1*). These findings suggest that, besides thiamin, yeasts may experience limitation of other nutrients when propagating under harsh industrial conditions (Stambuk et al., 2009; Della-Bianca et al., 2013). For example, the increased copy numbers of *GAP1* in the bioethanol yeasts may represent a compensation for the limited nitrogen content of sugarcane-based substrates in industrial fermentation (Della-Bianca et al., 2013). The amplification of *GAP1* is also interesting because its product has a function in the uptake of polyamines (Uemura et al., 2005), a class of compounds implicated in stress mitigation in alcoholic fermentation (Kim et al., 2015).

Mineral nutrition and/or toxicity may also be important for yeasts during fuel ethanol production, as suggested by amplification of *ARR* cluster for arsenic detoxification we observed in 20 bioethanol strains. The high iron content in molasses possibly represents an additional critical issue (Basso et al., 2011). In yeasts *OPT2* is a genetic interactor of key iron homeostasis components (Elbaz-Alon et al., 2014) and defective *OPT2* alleles are widespread in bioethanol strains. We found that additional genes encoding factors for iron acquisition are also defective (e.g., *FRE3* alleles) or are deleted (e.g., the *FIT2/FIT3/FRE5* cluster) in many bioethanol strains, suggesting a putative downregulation of the ferric iron-siderophore uptake system in face of a high iron content in the environment. However, a similar deletion of the *FIT2/FIT3/FRE5* region in the Chr. XV in flor yeasts was instead linked to an enhanced sensibility to iron (Eldarov et al., 2018; Eldarov et al., 2019). Another contradiction is the *FRE2* amplification in several bioethanol strains, suggesting that probably a more complex remodeling of the iron uptake system may be under selection. This hitherto unknown connection between yeast iron and arsenic homeostasis with sugarcane fermentation emphasizes the need for testing the broad responses of bioethanol strains to mineral nutrition.

Other genetic markers of bioethanol strains could be implicated in varied stress responses. In the Brazilian sugarcane fermentation system temperatures frequently raise above 35°C (Della-Bianca et al., 2013). In at least six bioethanol strains (isolated from five different distilleries) we found amplifications related to a ~17.0 Kb segment on the Chr. III. The region encompasses the genes *HSP30* and *PMP1*, whose products are regulators of the plasma membrane H(+)-ATPase Pma1 (Navarre et al., 1992; Meena et al., 2011). The expression of Hsp30 under heat stress confers thermotolerance by the downregulation of Pma1, sparing ATP during the stress response (Meena et al., 2011). Interestingly, the segmental amplification of the *HSP30/PMP1* region on Chr. III has been linked to thermotolerance in experimental evolution populations of yeasts (Caspeta et al., 2014). While this connection of gene function with a stress response mechanism is possible for *HSP30*/*PMP1*, for some genetic features linked to bioethanol strains such association is not obvious. This is the case of genes whose functions have not been tested so far, such as *ECM34*, *CSS3*, *EO220_52800* and *SAM-mts*. The later corresponds to a multicopy gene family of *SAM-dependent methyltransferases* (Pfam family Methyltransf_11, PF08241) widespread in bioethanol strains. Similarly, the Region B is a non-S288c five-gene cluster that is amplified in bioethanol strains. It encodes typical genes related to stress response and nutrient acquisition (i.e., two transcriptional factors, a FLO-like flocullin, a high affinity nicotinic acid permease, and a 5-oxo-L-prolinase) (Borneman et al., 2011; Galeote et al., 2011). The Region B is a mobile and integrative element in the genome that likely propagates via a circular episomic intermediate (Borneman et al., 2011; Galeote et al., 2011). It is possible that, by mobilization and amplification, the Region B can provide a quick mechanism of adaptation to fluctuating stressful conditions related to industrial fermentation. Altogether, the examples discussed here demonstrate that a detailed knowledge about the key genetic features linked to bioethanol strains may be strategic for the genetic enhancement of yeast fermentation performance.

### Are Brazilian bioethanol yeasts historically derived from cachaça spirit strains?

To understand the evolution of the Brazilian bioethanol strains we performed a phylogenetic analysis including taxa from the major groups of industrial *S. cerevisiae*, encompassing in total 197 yeast genomes. This represents one of the first attempts to use an AF method for inferring phylogenetic relationships among eukaryotic genomes. Our phylogenetic tree, largely topologically congruent with that generated by Gallone et al. (2016), demonstrates the robustness and utility of AF approaches for inferring phylogenetic relationships using whole-genome sequences of eukaryotes, bypassing multiple sequence alignment. In our AF tree, the 42 fuel ethanol and cachaça strains are unified in a single monophyletic clade, sister to wine yeasts. Previous whole-genome phylogenetic and population structure analyses that included JAY291 and/or a few fuel ethanol yeasts already implied the close relationship of Brazilian bioethanol and cachaça strains with yeasts used in industrial wine making (Almeida et al., 2015; Barbosa et al., 2016; Gallone et al., 2016; Barbosa et al., 2018; Legras et al., 2018). For instance, a whole-genome SNP-based NJ tree of 188 taxa including 21 cachaça strains placed the majority of cachaça taxa in two sister clades, C1 and C2, together with the bioethanol strains (CBS7960, JAY291, and BG-1) and at the base of a clade containing the wine group (Barbosa et al., 2018). Our results revealed that cachaça strains and the bioethanol yeasts display typical signatures of domestication shared with Western European wine strains, such as the presence of Region B and deletions in *AQY1* (A881) and *AQY2* (11 bps), lending support to the earlier study (Barbosa et al., 2018). The monophyletic nature of the Brazilian bioethanol strains was also highlighted in the NJ-tree of 1,011 yeast genomes (Peter et al., 2018), in which 35 Brazilian bioethanol strains are clustered in a clade proximal to the wine and other European groups of *S. cerevisiae*. Our phylogenetic study comes therefore in timely manner, as it integrates multiple genome assemblies that had only recently became available, enabling a more-refined phylogenetic inference that provides a clear evidence for a single common origin of Brazilian bioethanol yeasts.

It should be noted that our analyses were largely restricted to *S. cerevisiae* isolates from ethanol plants in the state of São Paulo. Therefore, a broader representation of industrial yeasts from different regions in Brazil is necessary to achieve a more-comprehensive overview of the phylogenetic origin of bioethanol strains. This is particularly relevant giving that some cachaça isolates display different phylogenetic affiliations from most others that are more-closely related to wine and bioethanol yeasts (Barbosa et al., 2018), including strain SP005 that is phylogenetically closely associated with the Beer 1 clade in our tree (**Figure 5**). The origin of bioethanol strains outside the monophyletic clade remains to be investigated with more-broadly sampled data.

Considering our new data, we summarize the putative origin of Brazilian bioethanol *S. cerevisiae* in **Figure 7**. The present-day bioethanol yeasts were derived from “contaminants” that invaded the fermentation process and outcompeted the starter strains (Basso et al., 2008; Basso et al., 2011). A longstanding notion is that the “intruding” yeasts consisted of wild-type *S. cerevisiae* (Basso et al., 2008; Argueso et al., 2009; Antonangelo et al., 2013; Della-Bianca et al., 2013; Lopes et al., 2015). Our results suggest that Brazilian bioethanol yeasts share common genetic traits, are phylogenetically closely related to industrial European strains, and display signs of domestication. Therefore, we found no support for the notion that these strains may have derived multiple times from natural wild populations. Instead, comparative genomics and phylogenetics revealed that the Brazilian natural isolates of *S. cerevisiae* represent separate lineages that are unrelated to bioethanol yeasts, and are devoid of typical domestication marks, such as defective aquaporin alleles (Barbosa et al., 2016; Barbosa et al., 2018).

**FIGURE 7.**
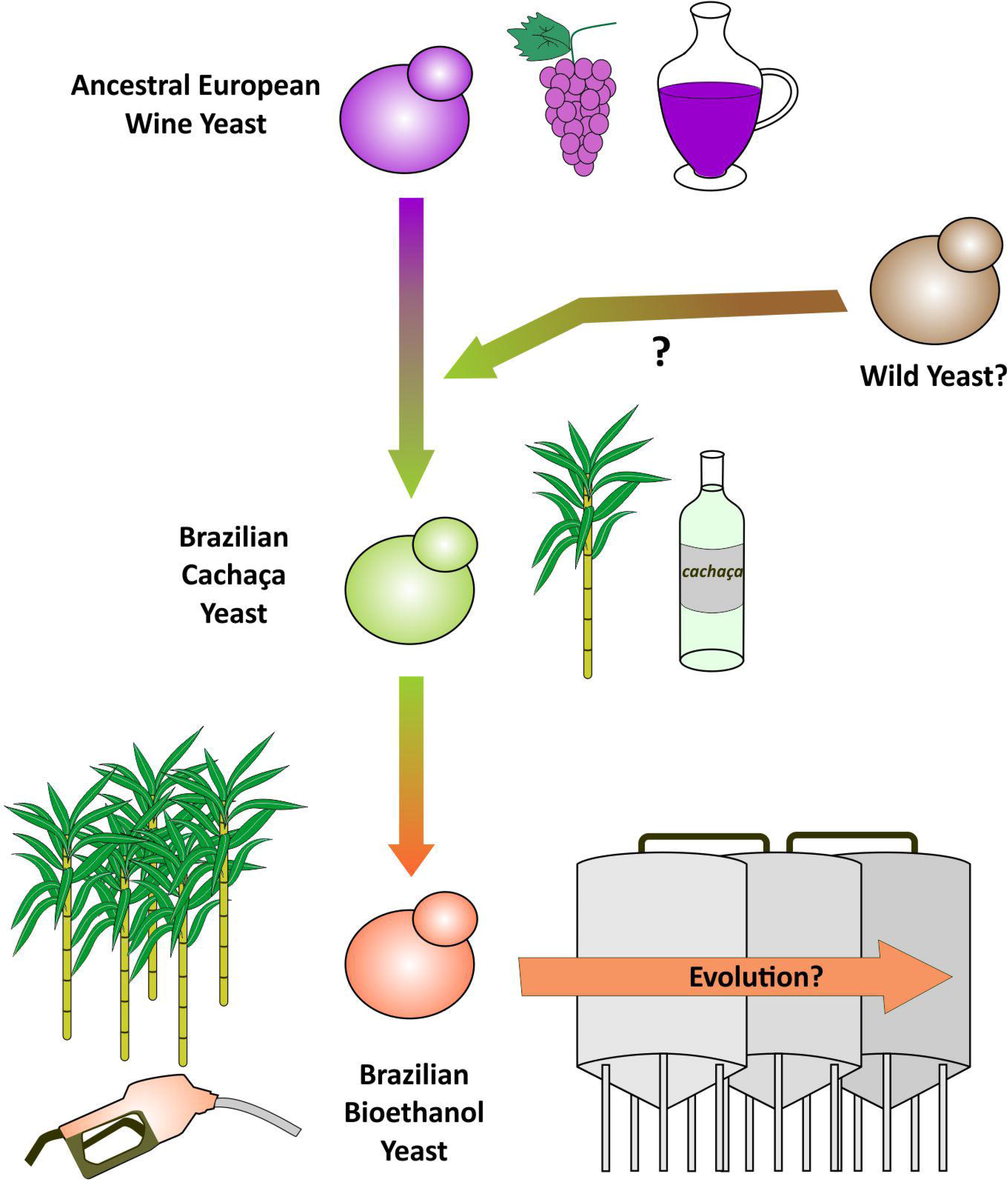
Model for the evolution of Brazilian bioethanol yeasts. Cachaça yeasts probably derive from an ancestral European yeast lineage domesticated for the wine-making process. Since cachaça and bioethanol yeasts have a mosaic genome structure, it is likely that at some point a hybridization event occurred between an European ancestor and a yet-unknown yeast (perhaps from a South American wild population?). Cachaça yeasts were latter co-opted for biofuel production becoming the modern-day bioethanol yeasts. The degree in which bioethanol yeasts may evolve within the process of fermentation is still unknow. This process-related evolution may account for the genetic divergence between cachaça and fuel ethanol strains.

A more likely scenario for the origin of bioethanol strains is through their European ancestry (**Figure 7**). Barbosa et al. (2018) proposed a model supporting that cachaça strains from clades C1 and C2 originated from an European stock related to the wine-making process, possibly following immigration routes during the colonization of Brazil by Portuguese settlers. From this perspective, the adaptation of European strains to sugarcane fermentation for cachaça distillation was facilitated by primary domestication traits already encoded in the genomes of wine strains, such as loss of aquaporins, and presence of Regions B and C. A long period of process-specific adaptation of those early cachaça strains was accompanied by emergence of new traits that improved their capacity to ferment sugarcane substrates. This led to the divergence of cachaça strains from the wine yeasts, representing a secondary domestication event. Among newly selected traits are additional aquaporin defective alleles, acquisition of *RTM1*, *FZF1 C*, and genes for biotin prototrophy, loss of *SSU1-R*, and specific introgressions from *S. paradoxus*. The introgression of exogenous genes suggests that, at some point, wine European strains, or early cachaça lineages, may have bred with natural *S. cerevisiae* or *S. paradoxus* populations (**Figure 7**). In fact, in population structure studies, genomes of bioethanol and cachaça strains are classified as “mosaic” (Gallone et al., 2016; Peter et al., 2018), sharing a strong signal of European wine yeasts while displaying a detectable inheritance from an alternative source(s) (Barbosa et al., 2018; Legras et al., 2018). The possibility of admixture is also consistent with the presence of putative *S. paradoxus* introgressions that are shared among bioethanol, cachaça, and natural Brazilian *S. cerevisiae* populations (Barbosa et al., 2016; Barbosa et al., 2018). Among those genes are *OPT1* and its neighbor ORF *YJL213W*, which are also present in H4, JAY291, CNV, CNP bioethanol strains as part of a large 14-gene cluster found on Chr. X left subtelomere. Interestingly, the presence of this cluster is largely associated with South, Central, and North American *S. cerevisiae* (**Supplementary Figure 3** and **Supplementary Table 12**).

As bioethanol strains are phylogenetically closely related to C1 and C2 cachaça strains (Barbosa et al., 2018), it is logical to propose a common evolutionary history in the wine-cachaça-bioethanol lineage (**Figure 7**). In Brazil, strains for cachaça production have been segregating for centuries along anthropic environments dedicated to sugar milling and cachaça fermentation/distillation, whereas present-day ethanol plants are frequently associated to sugar factories (Basso et al., 2011; Basso and Lino, 2019). Conceivably, when the new technological process of fuel ethanol production was launched in the 20^th^ century, it is possible that physical overlapping (or equipment sharing) between traditional sugarcane processing facilities and the new biofuel plants may have facilitated the cooption of cachaça strains for the emerging fermentation industry. Alternatively, cachaça yeasts may have somehow spread into sugarcane crops, from where they frequently have reentered into the fermentation process for fuel ethanol production. Regardless of the evolutionary path, if Brazilian fuel ethanol yeasts indeed originated from cachaça strains, they should not be regarded as “contaminants”, in a sense that they are actually inherent (i.e., domesticated) to the process and environment of sugarcane fermentation.

Finally, the degree to which bioethanol strains are evolving within the fermentation process should also be investigated by using genomic approaches (**Figure 7**). The industrial fermentation technology in Brazil is based on yeast biomass recycling and reuse in a continuous manner throughout the year, setting up an evolutionary experiment in which new mutations may arise and be selected according to their fitness contribution to withstand harsh industrial conditions. In fact, the frequently observed shifts in yeast karyotype patterns during industrial fermentation of sugarcane substrates indicate dynamic occurrences of chromosomal structural variations (Basso et al., 2008; Lopes et al., 2015). Moreover, recent mechanization of sugarcane harvesting has altered the biomass composition and processing, resulting in a higher input of toxic compounds into molasses (e.g., phenols) (Basso et al., 2019). This is aggravated by raises in sugar commodity prices, which have stimulated increase in production via reiterated cycles of crystal sugar extraction, resulting in exhausted molasses enriched in fermentation inhibitors (Basso et al., 2019). Therefore, new factors are further intensifying selective pressures over bioethanol yeasts, with consequences for their evolution. Such putative process-associated evolution may constitute yet a tertiary domestication event in the wine-cachaça-bioethanol lineage, in which bioethanol strains are dynamically acquiring adaptive genetic variations specific to the biofuel industry.

## Supporting information

Supplementary Sequence File 1

## DATA AVAILABILITY STATEMENT

Genome sequence data for H3 and H4 are available at NCBI (https://www.ncbi.nlm.nih.gov/) under the BioProject accession number PRJEB31792. Accession numbers from other genomes analyzed in this work are available in Supplementary Tables 1, 4, 5, and 6.

## AUTHOR CONTRIBUTIONS

APJ, JG, CXC: Project conceptualization; LCB: Strain/spore isolation characterization; APJ, LCB, JF: DNA prep, Genome library prep, Genome sequencing; APJ, JG, Genome Assembly; APJ, JG, TGS, PY, RGP, YC, CXC: Genome annotation, diverse bioinformatic analyses, statistical analyses; APJ, CXC, JG: Manuscript writing.

## ACKNOWLEDGMENTS

This work is supported by the computational resources of the Australian National Computational Infrastructure (NCI) National Facility systems through the NCI Merit Allocation Scheme (Project d85) awarded to CXC.

## SUPPLEMENTARY MATERIAL

The Supplementary Material for this article can be found online at:

