## Supplementary Figures for "Comparative genomics supports that Brazilian bioethanol *Saccharomyces cerevisiae* comprise a unified group of domesticated strains related to cachaça spirit yeasts"

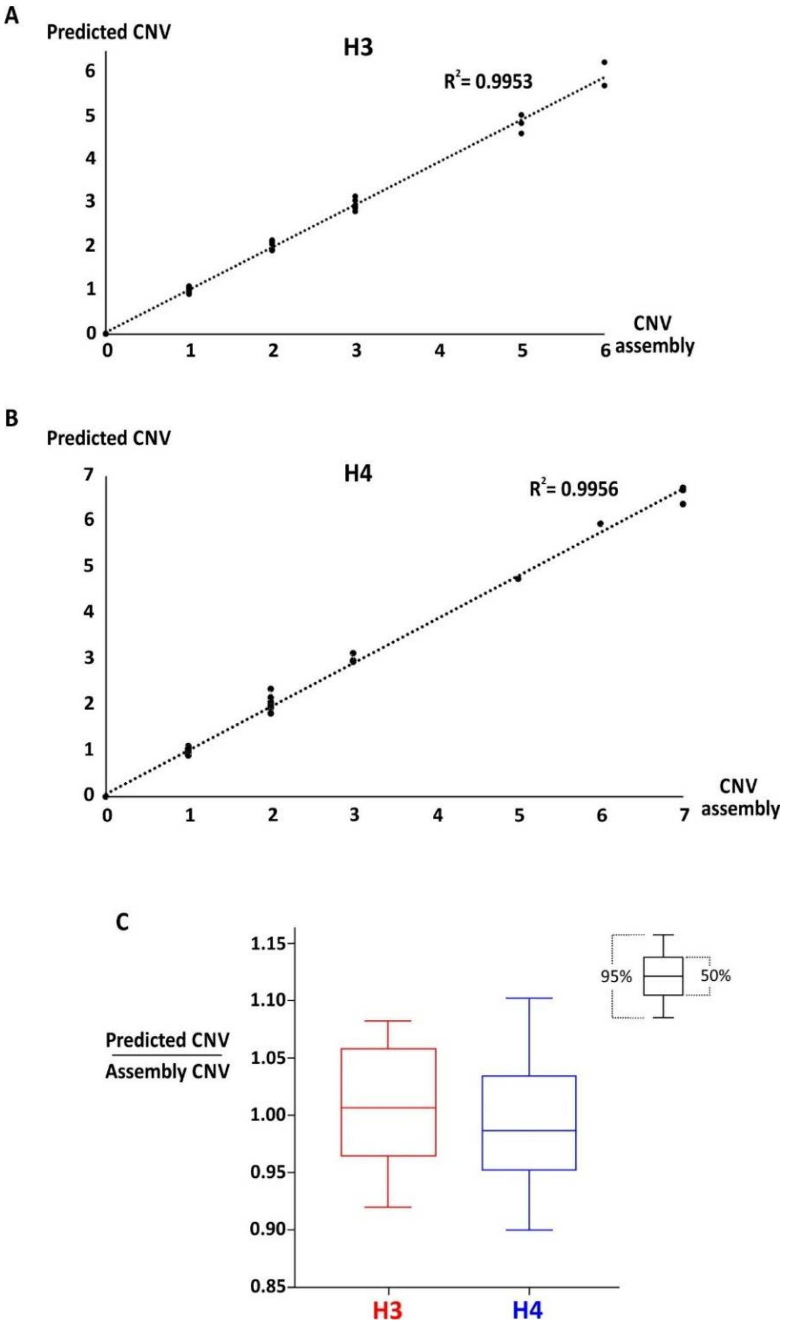


**SUPPLEMENTARY FIGURE 1|** Validation of the procedure for counting genes/regions copy numbers based on read depth. For H3 **(A)** and H4 **(B)** a strict correlation was found by counting copy numbers of 43 genes/regions from Illumina read depth and from the expected numbers based on the genome assembly. For H3: Pearson's $r = 0.9976$, Confidence Interval 95% (0.9956, 0.9987), $p$-value $\leq 0.0001$; For H4: Pearson's $r = 0.9978$, Confidence Interval 95% (0.9959 to 0.9988), $p$-value $\leq 0.0001$. **(C)** Box plot correlating estimation of copy numbers by read depth with actual genome assembly numbers for 37 (H3) and 39 (H4) genes/regions, respectively. For H3 all predictions and for H4 most of the estimations fell within a window of over than 90% accuracy from the actual numbers captured by genome assembly.

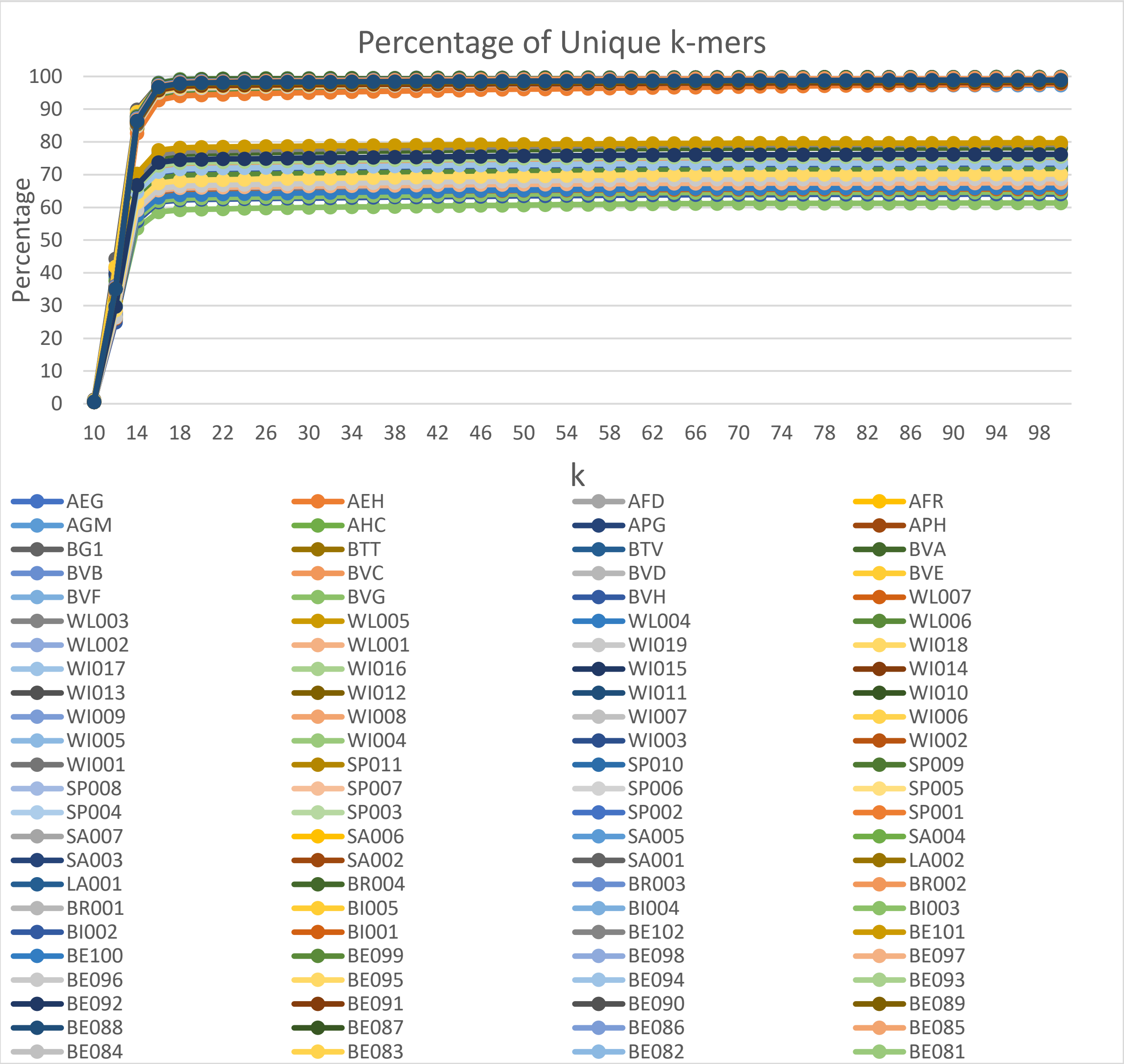


**SUPPLEMENTARY FIGURE 2|** Percentage of unique *k*-mers in representative strains of our dataset. Y-axis represents the percentage of unique *k*-mers and x-axis displays the size of *k*-mers (*k*). Trends of unique *k*-mers percentage, obtained for each strain, collectively indicate *k* = 18 as the minimum *k* to be used for deriving $D_2^S$ distances between all possible genome-pairs. At the lower part of the figure, a color scheme identifies the strains plotted in the graph.

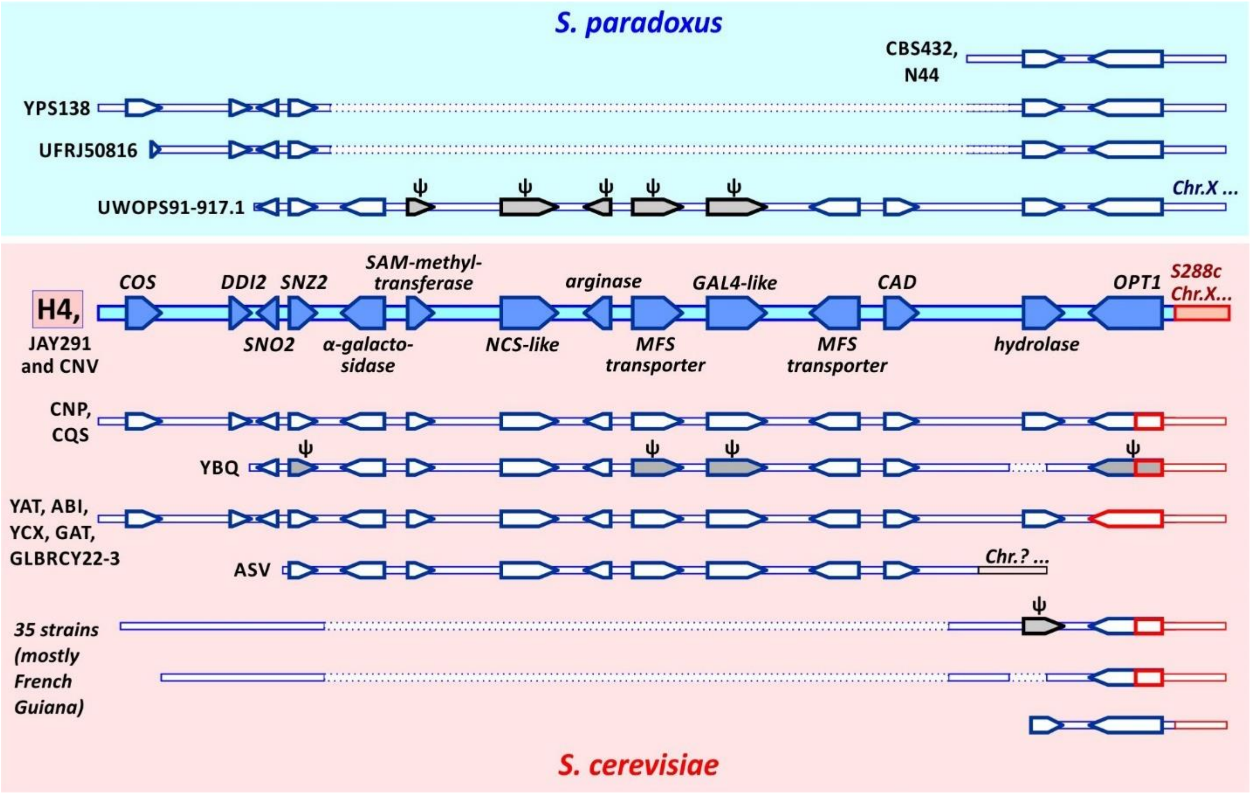


**SUPPLEMENTARY FIGURE 3|** Syntenic conservation of the H4 34.7 kb region at the left-end of Chr. X. BLASTN searches against the dataset of the 1,002 yeasts genome project and to GenBank (https://blast.ncbi.nlm.nih.gov/Blast.cgi) did identify a few strains that have the gene cluster (see **Supplementary Table 12**). The corresponding structures are represented in synteny to the H4 region, for which structural genes are labeled and highlighted at the center of the figure. The region has been potentially introgressed from *S. paradoxus*, and structures found at the Chr. X of *S. paradoxus* taxa are depicted at the upper part (blue background) of the figure, while at the lower part are displayed clusters of genes observed in *S. cerevisiae* strains (reddish background). The *OPT1* gene represents the potential spot where *S. paradoxus* genes (blue) have recombined with the *S. cerevisiae* Chr. X (red) (See below **Supplementary Figure 4**). This might explain why in some *S. cerevisiae* strains the *OPT1* is chimeric, having part of the *S. paradoxus* gene (blue) and a complement of the S288c-type gene (red). Putative pseudogenes (ψ) are colored in grey. Deletions are represented by dot lines and white background separating syntenic chromosome structures. Strains and gene labels are described on the **Supplementary Table 12**.

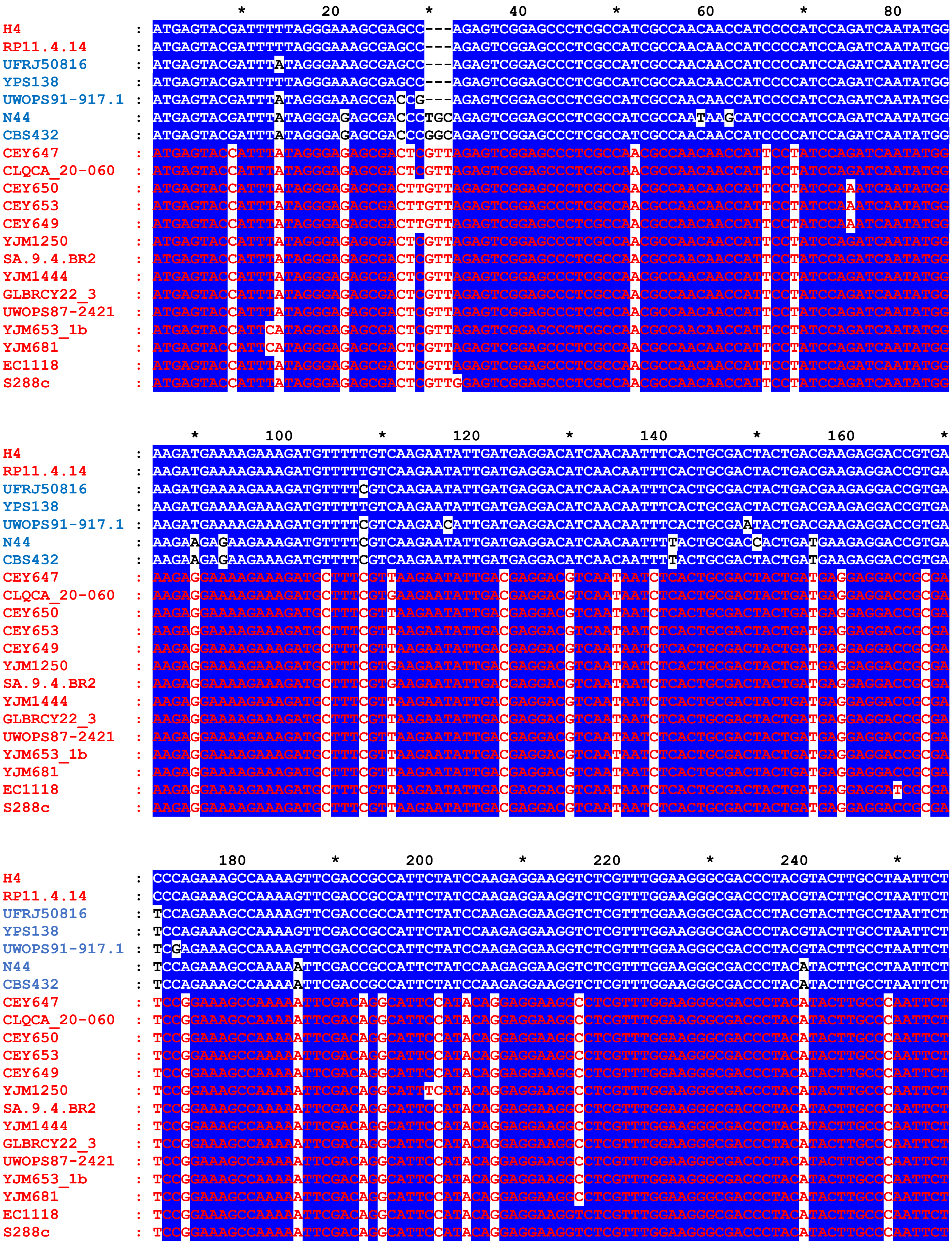
H4
RP11.4.14
UFRJ50816
YPS138
UWOPS91-917.1
N44
CBS432
CEY647
CLQCA_20-060
CEY650
CEY653
CEY649
YJM1250
SA.9.4.BR2
YJM1444
GLBRCY22_3
UWOPS87-2421
YJM653_1b
YJM681
EC1118
S288c
20
40
60
80
100
120
140
160
180
200
220
240

```
                     260       *       280       *       300       *       320       *       340
H4              : CCATATCCCGAAGTTAGGTCTGCGGTCTCCATTGAGGATGATCCTACCATCCGTCTCAACCACTGGAGGACTTGGTTCTTAACCA
RP11.4.14       : CCATATCCCGAAGTTAGGTCTGCGGTCTCCATTGAGGATGATCCTACCATCCGTCTCAACCACTGGAGGACTTGGTTCTTAACCA
UFRJ50816       : CCATATCCCGAAGTTAGGTCTGCGGTCTCCATTGAGGATGATCCTACCATTCGCCTCAACCACTGGAGGACTTGGTTCTTAACCA
YPS138          : CCATATCCCGAAGTTAGGTCTGCGGTCTCCATTGAGGATGATCCTACCATCCGCCTCAACCACTGGAGGACTTGGTTCTTAACCA
UWOPS91-917.1   : CCATATCCCGAAGTTAGGTCTGCGGTCTCCATTGAGGATGATCCTACCATCCGCCTCAACCACTGGAGGACTTGGTTCTTAACCA
N44             : CCATATCCCGAAGTGAGGTCTGCGGTCTCCATTGAGGATGATCCCACCATCCGCCTCAACCACTGGAGGACTTGGTTCTTAACCA
CBS432          : CCATATCCCGAAGTGAGGTCTGCGGTCTCCATTGAGGATGATCCCACCATCCGCCTCAACCACTGGAGGACTTGGTTCTTAACCA
CEY647          : CCATATCCTGAAGTGAGATCGGCGGTGTCCATCGAGGATGACCCCACCATCCGCCTCAACCACTGGAGAACCTGGTTCTTGACCA
CLQCA_20-060    : CCATATCCTGAAGTGAGATCGGCGGTGTCCATCGAGGATGACCCCACCATCCGCCTCAACCACTGGAGAACCTGGTTCTTGACCA
CEY650          : CCATATCCTGAAGTGAGATCGGCGGTGTCCATCGAGGATGACCCCACCATCCGCCTCAACCACTGGAGAACCTGGTTCTTGACCA
CEY653          : CCATATCCTGAAGTGAGATCGGCGGTGTCCATCGAGGATGACCCCACCATCCGCCTCAACCACTGGAGAACCTGGTTCTTGACCA
CEY649          : CCATATCCTGAAGTGAGATCGGCGGTGTCCATCGAGGATGACCCCACCATCCGCCTCAACCACTGGAGAACCTGGTTCTTGACCA
YJM1250         : CCATATCCTGAAGTGAGATCGGCGGTGTCCATCGAGGATGACCCCACCATCCGCCTCAACCACTGGAGAACCTGGTTCTTGACCA
SA.9.4.BR2      : CCATATCCTGAAGTGAGATCGGCGGTGTCCATCGAGGATGACCCCACCATCCGCCTCAACCACTGGAGAACCTGGTTCTTGACCA
YJM1444         : CCATATCCTGAAGTGAGATCGGCGGTGTCCATCGAGGATGACCCCACCATCCGCCTCAACCACTGGAGAACCTGGTTCTTGACCA
GLBRCY22_3      : CCATATCCTGAAGTGAGATCGGCGGTGTCCATCGAGGATGACCCCACCATCCGCCTCAACCACTGGAGAACCTGGTTCTTGACCA
UWOPS87-2421    : CCATATCCTGAAGTGAGATCGGCGGTGTCCATCGAGGATGACCCCACCATCCGCCTCAACCACTGGAGAACCTGGTTCTTGACCA
YJM653_1b       : CCATATCCTGAAGTGAGATCGGCGGTGTCCATCGAGGATGACCCCACCATCCGCCTCAACCACTGGAGAACGTGGTTCTTGACCA
YJM681          : CCATATCCTGAAGTGAGATCGGCGGTGTCCATCGAGGATGACCCCACCATCCGCCTCAACCACTGGAGAACGTGGTTCTTGACCA
EC1118          : CCATATCCTGAAGTGAGATCGGCGGTGTCCATCGAGGATGACCCCACCATCCGCCTCAACCACTGGAGAACCTGGTTCTTGACCA
S288c           : CCATATCCTGAAGTGAGATCGGCGGTGTCCATCGAGGATGACCCCACCATCCGCCTCAACCACTGGAGAACGTGGTTCTTGACCA

                            *       360       *       380       *       400       *       420
H4              : CAATATTTGTGGTAGTTTTCGCCGGGGTCAATCAGTTCTTTTCCCTAAGATATCCATCTTTAGAGATCAATTTTCTTGTCGCACA
RP11.4.14       : CAATATTTGTGGTAGTTTTCGCCGGGGTCAATCAGTTCTTTTCCCTAAGATATCCATCTTTAGAGATCAATTTTCTTGTCGCACA
UFRJ50816       : CGATATTTGTGGTAGTTTTCGCCGGGGTCAATCAGTTCTTTTCCCTAAGATATCCATCTTTAGAGATCAATTTTCTTGTCGCACA
YPS138          : CGATATTTGTGGTAGTTTTCGCCGGGGTCAATCAGTTCTTTTCCCTAAGATATCCATCTTTAGAGATCAATTTTCTGGTCGCACA
UWOPS91-917.1   : CGATATTCGTGGTAGTTTTCGCCGGGGTCAATCAGTTCTTTTCCCTAAGATATCCATCTTTAGAGATCAATTTTCTTGTCGCACA
N44             : CGATATTTGTGGTAGTTTTCGCCGGTGTCAATCAGTTCTTTTCCCTAAGATATCCATCTTTAGAGATCAATTTCCTTGTCGCACA
CBS432          : CGATATTTGTGGTAGTATTCGCCGGTGTCAATCAGTTCTTTTCCCTGAGATATCCATCTTTAGAGATCAATTTCCTTGTCGCACA
CEY647          : CGGTGTTTGTGGTAGTTTTCGCCGGTGTTAATCAATTTTTTTCCCTGAGATATCCATCGCTAGAGATCAACTTCCTTGTTGCACA
CLQCA_20-060    : CGGTGTTTGTGGTAGTTTTCGCCGGTGTTAATCAATTTTTTTCCCTGAGATATCCATCGCTAGAGATCAACTTCCTTGTTGCACA
CEY650          : CGGTGTTTGTGGTAGTTTTCGCCGGTGTTAATCAATTTTTTTCCCTGAGATATCCATCGCTAGAGATCAACTTCCTTGTTGCACA
CEY653          : CGGTGTTTGTGGTAGTTTTCGCCGGTGTTAATCAATTTTTTTCCCTGAGATATCCATCGCTAGAGATCAACTTCCTTGTTGCACA
CEY649          : CGGTGTTTGTGGTAGTTTTCGCCGGTGTTAATCAATTTTTTTCCCTGAGATATCCATCGCTAGAGATCAACTTCCTTGTTGCACA
YJM1250         : CGGTGTTTGTGGTAGTTTTCGCCGGTGTTAATCAATTTTTTTCCCTGAGATATCCATCGCTAGAGATCAACTTCCTTGTCGCACA
SA.9.4.BR2      : CGGTGTTTGTGGTAGTTTTCGCCGGTGTTAATCAATTTTTTTCCCTGAGATATCCATCGCTAGAGATCAACTTCCTTGTCGCACA
YJM1444         : CGGTGTTTGTGGTAGTTTTCGCCGGTGTTAATCAATTTTTTTCCCTGAGATATCCATCGCTAGAGATCAACTTCCTTGTTGCACA
GLBRCY22_3      : CGGTGTTTGTGGTAGTTTTCGCCGGTGTTAATCAATTTTTTTCCCTGAGATATCCATCGCTAGAGATCAACTTCCTTGTTGCACA
UWOPS87-2421    : CGGTGTTTGTGGTAGTTTTCGCCGGTGTTAATCAATTTTTTTCCCTGAGATATCCATCGCTAGAGATCAACTTCCTTGTTGCACA
YJM653_1b       : CGGTGTTTGTGGTAGTTTTCGCCGGTGTTAATCAATTTTTTTCCCTGAGATATCCATCGCTAGAGATCAACTTCCTTGTTGCACA
YJM681          : CGGTGTTTGTGGTAGTTTTCGCCGGTGTTAATCAATTTTTTTCCCTGAGATATCCATCGCTAGAGATCAACTTCCTTGTTGCACA
EC1118          : CGGTGTTTGTGGTAGTTTTCGCCGGTGTTAATCAATTTTTTTCCCTGAGATATCCATCGCTAGAGATCAACTTCCTTGTTGCACA
S288c           : CGGTGTTTGTGGTAGTTTTCGCCGGTGTTAATCAATTTTTTTCCCTGAGATATCCATCGCTAGAGATCAACTTCCTTGTTGCACA

                       *       440       *       460       *       480       *       500       *
H4              : GGTTGTTTGCTATCCAATAGGTAGAGTGCTGGCTCTCTTGCCCGACTGGAAGTGTCCCAAAGTACCATTTTTTGATTTGAACCCG
RP11.4.14       : GGTTGTTTGCTATCCAATAGGTAGAGTGCTGGCTCTCTTGCCCGACTGGAAGTGTCCCAAAGTACCATTTTTTGATTTGAACCCG
UFRJ50816       : GGTTGTTTGCTATCCAATAGGTAGAGTGCTGGCTCTCTTGCCCGACTGGAAGTGTCCCAAAGTACCATTTTTTGATTTGAACCCG
YPS138          : GGTTGTTTGCTATCCAATAGGTAGAGTACTGGCTCTCTTGCCCGACTGGAAGTGTCCCAAAGTACCATTTTTTGATTTGAACCCG
UWOPS91-917.1   : GGTTGTTTGCTATCCAATAGGTAGAGTGCTGGCTCTCTTGCCCGACTGGAAGTGTCCCAAAGTACCATTTTTTGATTTGAACCCG
N44             : GGTTGTTTGCTATCCAATAGGTAGAGTGCTGGCTCTCTTGCCCGACTGGAAGTGTCCCAAAGTACCATTCTTTGATTTGAACCCG
CBS432          : GGTTGTTTGCTATCCAATAGGTAGAGTGCTGGCTCTCTTGCCCGACTGGAAGTGTCCCAAAGTACCATTCTTTGATTTGAACCCG
CEY647          : AGTTGTTTGCTACCCAATTGGTAGGATACTGGCTCTCTTGCCCGACTGGAAGTGTTCTAAAGTGCCATTTTTCGATTTAAACCCG
CLQCA_20-060    : AGTTGTTTGCTACCCAATTGGTAGGATACTGGCTCTCTTGCCCGACTGGAAGTGTTCTAAAGTGCCATTTTTCGATTTAAACCCG
CEY650          : AGTTGTTTGCTACCCAATTGGTAGGATACTGGCTCTCTTGCCCGACTGGAAGTGTTCTAAAGTGCCATTTTTCGATTTAAACCCG
CEY653          : AGTTGTTTGCTACCCAATTGGTAGGATACTGGCTCTCTTGCCCGACTGGAAGTGTTCTAAAGTGCCATTTTTCGATTTAAACCCG
CEY649          : AGTTGTTTGCTACCCAATTGGTAGGATACTGGCTCTCTTGCCCGACTGGAAGTGTTCTAAAGTGCCATTTTTCGATTTAAACCCG
YJM1250         : AGTTGTTTGCTATCCAATTGGTAGGATACTGGCTCTCTTGCCCGACTGGAAGTGTTCTAAAGTGCCATTTTTTGATTTAAACCCG
SA.9.4.BR2      : AGTTGTTTGCTATCCAATTGGTAGGATACTGGCTCTCTTGCCCGACTGGAAGTGTTCTAAAGTGCCATTTTTTGATTTAAACCCG
YJM1444         : AGTTGTTTGCTACCCAATTGGTAGGATACTGGCTCTCTTGCCCGACTGGAAGTGTTCTAAAGTGCCATTTTTCGATTTAAACCCG
GLBRCY22_3      : AGTTGTTTGCTACCCAATTGGTAGGATACTGGCTCTCTTGCCCGACTGGAAGTGTTCTAAAGTGCCATTTTTCGATTTAAACCCG
UWOPS87-2421    : AGTTGTTTGCTACCCAATTGGTAGGATACTGGCTCTCTTGCCCGACTGGAAGTGTTCTAAAGTGCCATTTTTCGATTTAAACCCG
YJM653_1b       : AGTTGTTTGCTACCCAATTGGTAGGATACTGGCTCTCTTGCCCGACTGGAAGTGTTCTAAAGTGCCATTTTTCGATTTAAACCCG
YJM681          : AGTTGTTTGCTACCCAATTGGTAGGATACTGGCTCTCTTGCCCGACTGGAAGTGTTCTAAAGTGCCATTTTTCGATTTAAACCCG
EC1118          : AGTTGTTTGCTACCCAATTGGTAGGATACTGGCTCTCTTGCCCGACTGGAAGTGTTCTAAAGTGCCATTTTTCGATTTAAACCCG
S288c           : AGTTGTTTGCTACCCAATTGGTAGGATACTGGCTCTCTTGCCCGACTGGAAGTGTTCTAAAGTGCCATTTTTCGATTTAAACCCG
```

```
                                   520         *         540         *         560         *         580         *
H4                 : GGACCATTCACCAAAAAGGAACACGCCGTCGTTACTATTGCCGTGGCGCTCACTTCTTCCACTGCATACGCTATGTACATTTTGA
RP11.4.14          : GGACCATTCACCAAAAAGGAACACGCCGTCGTTACTATTGCCGTGGCGCTCACTTCTTCCACTGCATACGCTATGTACATTTTGA
UFRJ50816          : GGACCATTCACCAAAAAGGAACACGCCGTCGTTACTATTGCCGTGGCGCTCACTTCTTCCACTGCATACGCTATGTACATTTTGA
YPS138             : GGACCATTCACCAAAAAGGAACACGCCGTCGTTACTATTGCCGTGGCGCTCACTTCTTCCACTGCATACGCTATGTACATTTTGA
UWOPS91-917.1      : GGACCATTCACCAAAAAGGAACACGCCGTCGTTACTATTGCCGTGGCGCTCACTTCTTCCACTGCATACGCTATGTACATTTTGA
N44                : GGACCATTCACCAAAAAGGAACACGCCGTCGTTACTATTGCCGTGGCGCTCACTTCTTCCACTGCATACGCTATGTACATTTTGA
CBS432             : GGACCATTCACCAAAAAGGAACATGCCGTCGTTACTATTGCCGTGGCGCTCACTTCTTCCACTGCATACGCTATGTACATTTTGA
CEY647             : GGCCCATTTACCAAAAAGGAACACGCCGTGGTCACAATTGCCGTGGCGCTTACTTCCTCCACTGCATACGCTATGTACATTTTGA
CLQCA_20-060       : GGCCCATTTACCAAAAAGGAACACGCCGTGGTCACAATTGCCGTGGCGCTTACTTCCTCCACTGCATACTCTATGTACATTTTGA
CEY650             : GGCCCATTTACCAAAAAGGAACACGCCGTGGTCACAATTGCCGTGGCGCTTACTTCCTCCACTGCATACGCTATGTACATTTTGA
CEY653             : GGCCCATTTACCAAAAAGGAACACGCCGTGGTCACAATTGCCGTGGCGCTTACTTCCTCCACTGCATACGCTATGTACATTTTGA
CEY649             : GGCCCATTTACCAAAAAGGAACACGCCGTGGTCACAATTGCCGTGGCGCTTACTTCCTCCACTGCATACGCTATGTACATTTTGA
YJM1250            : GGCCCATTTACCAAAAAGGAACACGCCGTGGTCACAATTGCCGTGGCGCTTACTTCCTCCACTGCATACGCTATGTACATTTTGA
SA.9.4.BR2         : GGCCCATTTACCAAAAAGGAACACGCCGTGGTCAGGATTGCCGTGGCGCTTACTTCCTCCACTGCATACGCTATGTACATTTTGA
YJM1444            : GGCCCATTTACCAAAAAGGAACACGCCGTGGTCACAATTGCCGTGGCGCTTACTTCCTCCACTGCATACGCTATGTACATTTTGA
GLBRCY22_3         : GGCCCATTTACCAAAAAGGAACACGCCGTGGTCACAATTGCCGTGGCGCTTACTTCCTCCACTGCATACGCTATGTACATTTTGA
UWOPS87-2421       : GGCCCATTTACCAAAAAGGAACACGCCGTGGTCACAATTGCCGTGGCGCTTACTTCCTCCACTGCATACGCTATGTACATTTTGA
YJM653_1b          : GGCCCATTTACCAAAAAGGAACACGCCGTGGTCACAATTGCTGTGGCGCTTACTTCCTCCACTGCATACGCTATGTACATTTTGA
YJM681             : GGCCCATTTACCAAAAAGGAACACGCCGTGGTCACAATTGCTGTGGCGCTTACTTCCTCCACTGCATACGCTATGTACATTTTGA
EC1118             : GGCCCATTTACCAAAAAGGAACACGCCGTGGTCACAATTGCCGTGGCGCTTACTTCCTCCACTGCATACGCTATGTACATTTTGA
S288c              : GGCCCATTTACCAAAAAGGAACACGCCGTGGTCACAATTGCCGTGGCGCTTACTTCCTCTACTGCATACGCTATGTACATTTTGA

                          600         *         620         *         640         *         660         *         680
H4                 : ACGCTCAAGGAAGTTTCTACAATATGAAGCTAAACGTCGGATATCAGTTCTTGTTGGTTTGGACATCACAGATGATTGGTTATGG
RP11.4.14          : ACGCTCAAGGAAGTTTCTACAATATGAAGCTAAACGTCGGATATCAGTTCTTGTTGGTTTGGACATCACAGATGATTGGTTATGG
UFRJ50816          : ACGCTCAAGGAAGTTTCTACAATATGAAGCTAAACGTCGGATATCAGTTCTTGTTGGTTTGGACATCACAGATGATTGGTTATGG
YPS138             : ACGCTCAAGGAAGTTTCTACAATATGAAGCTAAACGTCGGATATCAGTTCTTGTTGGTTTGGACATCACAGATGATTGGTTATGG
UWOPS91-917.1      : ACGCTCAAGGAAGTTTCTACAATATGAAGCTAAACGTCGGATATCAGTTCTTGTTGGTTTGGACATCACAGATGATTGGTTATGG
N44                : ACGCTCAAGGAAGTTTCTACAATATGAAGCTAAACGTCGGATATCAGTTCTTGTTGGTTTGGACATCACAGATGATTGGCTATGG
CBS432             : ACGCTCAAGGAAGTTTCTACAATATGAAGCTAAACGTCGGATATCAGTTCTTGTTGGTTTGGACATCACAGATGATTGGTTATGG
CEY647             : ACGCCCAGGGAAGCTTTTACAATATGAAACTGAATGTCGGATATCAGTTCTTGTTGGTTTGGACATCTCAAATGATTGGTTATGG
CLQCA_20-060       : ACGCCCAGGGAAGCTTTTACAATATGAAACTGAATGTCGGATATCAGTTCTTGTTGGTTTGGACATCACAGATGATTGGTTATGG
CEY650             : ACGCCCAGGGAAGCTTTTACAATATGAAACTGAATGTCGGATATCAGTTCTTGTTGGTTTGGACATCTCAAATGATTGGTTATGG
CEY653             : ACGCCCAGGGAAGCTTTTACAATATGAAACTGAATGTCGGATATCAGTTCTTGTTGGTTTGGACATCTCAAATGATTGGTTATGG
CEY649             : ACGCCCAGGGAAGCTTTTACAATATGAAACTGAATGTCGGATATCAGTTCTTGTTGGTTTGGACATCTCAAATGATTGGTTATGG
YJM1250            : ACGCCCAGGGAAGCTTTTACAATATGAAACTGAATGTCGGATATCAGTTCTTGTTGGTTTGGACATCTCAAATGATTGGTTATGG
SA.9.4.BR2         : ACGCCCAGGGAAGCTTTTACAATATGAAACTGAATGTCGGATATCAGTTCTTGTTGGTTTGGACATCTCAAATGATTGGTTATGG
YJM1444            : ACGCCCAGGGAAGCTTTTACAATATGAAACTGAATGTCGGATATCAGTTCTTGTTGGTTTGGACATCTCAAATGATTGGTTATGG
GLBRCY22_3         : ACGCCCAGGGAAGCTTTTACAATATGAAACTGAATGTCGGATATCAGTTCTTGTTGGTTTGGACATCTCAAATGATTGGTTATGG
UWOPS87-2421       : ACGCCCAGGGAAGCTTTTACAATATGAAACTGAATGTCGGATATCAGTTCTTGTTGGTTTGGACATCTCAAATGATTGGTTATGG
YJM653_1b          : ACGCCCAGGGAAGCTTTTACAACATGAAATTGAATGTCGGATATCAGTTCTTGTTGGTTTGGACATCTCAAATGATTGGTTATGG
YJM681             : ACGCCCAGGGAAGCTTTTACAACATGAAATTGAATGTCGGATATCAGTTCTTGTTGGTTTGGACATCTCAAATGATTGGTTATGG
EC1118             : ACGCCCAGGGAAGCTTTTACAATATGAAATTGAATGTCGGATATCAGTTCTTGTTGGTTTGGACATCTCAAATGATTGGTTATGG
S288c              : ACGCCCAGGGAAGCTTTTACAACATGAAACTTAATGTCGGATATCAGTTCTTGTTGGTTTGGACATCTCAAATGATTGGTTATGG

                                 *         700         *         720         *         740         *         760
H4                 : TGCTGCAGGTCTGACTAGAAGATGGGTTGTCAACCCTGCGAGCTCCGTCTGGCCTCAAACTCTAATTTCAGTGTCATTGTTTGAT
RP11.4.14          : TGCTGCAGGTCTGACTAGAAGATGGGTTGTCAACCCTGCGAGCTCCGTCTGGCCTCAAACTCTAATTTCAGTGTCATTGTTTGAT
UFRJ50816          : TGCTGCAGGTCTGACCAGAAGATGGGTTGTCAACCCTGCAAGCTCCGTCTGGCCTCAAACTCTAATTTCAGTGTCATTGTTTGAT
YPS138             : TGCTGCAGGTCTGACCAGAAGATGGGTTGTCAACCCTGCAAGCTCCATCTGGCCTCAAACTCTAATTTCAGTGTCATTGTTTGAT
UWOPS91-917.1      : TGCTGCAGGTCTGACCAGAAGATGGGTTGTCAACCCTGCAAGCTCCGTCTGGCCTCAAACTCTAATTTCAGTGTCATTGTTTGAT
N44                : TGCTGCGGGTCTGACCAGAAGATGGGTCGTCAACCCTGCAAGCTCCATCTGGCCTCAAACTCTAATTTCAGTGTCATTGTTTGAT
CBS432             : TGCTGCGGGTCTGACCAGAAGATGGGTCGTCAACCCTGCAAGCTCCATCTGGCCTCAAACTCTAATTTCAGTGTCATTGTTTGAT
CEY647             : TGCTGCAGGTCTGACCAGAAGATGGGTTGTCAACCCTGCAAGCTCCATCTGGCCTCAAACTCTAATTTCAGTGTCATTGTTTGAT
CLQCA_20-060       : TGCTGCAGGTCTGACCAGAAGATGGGTTGTCAACCCTGCAAGCTCCATCTGGCCTCAAACTCTAATTTCAGTGTCATTGTTTGAT
CEY650             : TGCTGCAGGTCTTACCAGAAGATGGGTCGTCAACCCTGCAAGCTCCATCTGGCCTCAAACTTTAATTTCAGTGTCATTGTTTGAT
CEY653             : TGCTGCAGGTCTTACCAGAAGATGGGTCGTCAACCCTGCAAGCTCCATCTGGCCTCAAACTTTAATTTCAGTGTCATTGTTTGAT
CEY649             : TGCTGCAGGTCTTACCAGAAGATGGGTCGTCAACCCTGCAAGCTCCATCTGGCCTCAAACTTTAATTTCAGTGTCATTGTTTGAT
YJM1250            : TGCTGCAGGTCTTACCAGAAGATGGGTCGTCAACCCTGCAAGCTCCATCTGGCCTCAAACTTTAATTTCAGTGTCATTGTTTGAT
SA.9.4.BR2         : TGCTGCGGGTCTTACCAGAAGATGGGTCGTCAACCCTGCAAGCTCTATCTGGCCTCAGACTTTAATTTCAGTGTCCTTGTTTGAT
YJM1444            : TGCTGCAGGTCTTACCAGAAGATGGGTCGTCAACCCTGCAAGCTCTATCTGGCCTCAAACTTTAATTTCAGTGTCATTGTTTGAT
GLBRCY22_3         : TGCTGCAGGTCTTACCAGAAGATGGGTCGTCAACCCTGCAAGCTCTATCTGGCCTCAAACTTTAATTTCAGTGTCATTGTTTGAT
UWOPS87-2421       : TGCTGCAGGTCTTACCAGAAGATGGGTCGTCAACCCTGCAAGCTCTATCTGGCCTCAAACTTTAATTTCAGTGTCATTGTTTGAT
YJM653_1b          : TGCTGCAGGTCTTACCAGAAGATGGGTCGTCAACCCTGCAAGCTCCATCTGGCCTCAAACTTTAATTTCAGTGTCATTGTTTGAT
YJM681             : TGCTGCAGGTCTTACCAGAAGATGGGTCGTCAACCCTGCAAGCTCCATCTGGCCTCAAACTTTAATTTCAGTGTCATTGTTTGAT
EC1118             : TGCTGCAGGTCTTACCAGAAGATGGGTCGTCAACCCTGCAAGCTCTATCTGGCCTCAGACTTTAATTTCAGTGTCCTTGTTTGAT
S288c              : TGCTGCAGGTCTTACCAGAAGATGGGTCGTCAACCCTGCAAGCTCTATCTGGCCTCAGACTTTAATTTCAGTGTCCTTGTTTGAT
```

```
                         *       780         *       800         *       820         *       840         *
H4                 : TCGCTGCACTCAAGAAAAGTTGAAAAGACAGTCGCAAACGGTTGGACGATGCCTCGTTACAGGTTTTTCTTGATTGTTCTTATCG
RP11.4.14          : TCGCTGCACTCAAGAAAAGTTGAAAAGACAGTCGCAAACGGTTGGACGATGCCTCGTTACAGGTTTTTCTTGATTGTTCTTATCG
UFRJ50816          : TCGCTGCACTCAAGAAAAGTTGAAAAGACAGTCGCAAACGGTTGGACGATGCCTCGTTACAGGTTTTTCTTGATTGTTCTTATCG
YPS138             : TCGCTGCACTCAAGAAAAGTTGAAAAGACAGTCGCAAACGGTTGGACGATGCCTCGTTACAGGTTTTTCTTGATTGTTCTTATCG
UWOPS91-917.1      : TCGCTGCACTCAAGAAAAGTTGAAAAGACAGTCGCAAACGGTTGGACGATGCCTCGTTACAGGTTTTTCTTGATTGTTCTTATCG
N44                : TCGCTGCACTCAAGAAAAGTTGAAAAGACAGTCGCAAACGGTTGGACGATGCCCCGTTACAGGTTTTTCTTGATTGTTCTTATCG
CBS432             : TCGCTGCACTCAAGAAAAGTTGAAAAGACAGTCGCAAACGGTTGGACGATGCCCCGTTACAGGTTTTTCTTGATTGTTCTTATCG
CEY647             : TCGCTGCACTCAAGAAAAGTTGAAAAGACAGTCGCAAACGGTTGGACGATGCCTCGTTACAGGTTTTTCTTGATTGTTCTTATCG
CLQCA_20-060       : TCGCTGCACTCAAGAAAAGTTGAAAAGACAGTCGCAAACGGTTGGACGATGCCTCGTTACAGGTTTTTCTTGATTGTTCTTATCG
CEY650             : TCGTTGCACTCGAGAAAAGTTGAAAAGACAGTCGCAAATGGTTGGACGATGCCCCGTTACAGGTTCTTCTTAATCGTTCTTATCG
CEY653             : TCGTTGCACTCGAGAAAAGTTGAAAAGACAGTCGCAAATGGTTGGACGATGCCCCGTTACAGGTTCTTCTTAATCGTTCTTATCG
CEY649             : TCGTTGCACTCGAGAAAAGTTGAAAAGACAGTCGCAAATGGTTGGACGATGCCCCGTTACAGGTTCTTCTTAATCGTTCTTATCG
YJM1250            : TCGTTGCACTCGAGAAAAGTTGAAAAGACAGTCGCAAATGGTTGGACGATGCCCCGTTACAGGTTCTTCTTAATCGTTCTTATCG
SA.9.4.BR2         : TCGTTGCACTCGAGAAAAGTTGAAAAGACAGTCGCAAATGGTTGGACGATGCCCCGTTACAGGTTCTTCTTAATCGTCCTTATCG
YJM1444            : TCGTTGCACTCGAGAAAAGTTGAAAAGACAGTCGCAAATGGTTGGACGATGCCCCGTTACAGGTTCTTCTTAATCGTTCTTATCG
GLBRCY22_3         : TCGTTGCACTCGAGAAAAGTTGAAAAGACAGTCGCAAATGGTTGGACGATGCCCCGTTACAGGTTCTTCTTAATCGTTCTTATCG
UWOPS87-2421       : TCGTTGCACTCGAGAAAAGTTGAAAAGACAGTCGCAAATGGTTGGACGATGCCCCGTTACAGGTTCTTCTTAATCGTTCTTATCG
YJM653_1b          : TCGTTGCACTCGAGAAAAGTTGAAAAGACAGTCGCAAATGGTTGGACGATGCCCCGTTACAGGTTCTTCTTAATCGTCCTTATCG
YJM681             : TCGTTGCACTCGAGAAAAGTTGAAAAGACAGTCGCAAATGGTTGGACGATGCCCCGTTACAGGTTCTTCTTAATCGTCCTTATCG
EC1118             : TCGTTGCACTCGAGAAAAGTTGAAAAGACAGTCGCAAATGGTTGGACGATGCCCCGTTACAGGTTCTTCTTAATCGTCCTTATCG
S288c              : TCGTTGCACTCGAGAAAAGTTGAAAAGACAGTCGCAAATGGTTGGACGATGCCCCGTTACAGGTTCTTCTTAATCGTCCTTATCG

                            860         *       880         *       900         *       920         *
H4                 : GATCCTTCGTCTGGTATTGGGTACCTGGATTCCTCTTCACTGGCCTGTCCTATTTCAACGTCATTTTATGGGGGTCGAAGACAAG
RP11.4.14          : GATCCTTCGTCTGGTATTGGGTACCTGGATTCCTCTTCACTGGCCTGTCCTATTTCAACGTCATTTTATGGGGGTCGAAGACAAG
UFRJ50816          : GATCCTTCGTCTGGTATTGGGTACCTGGATTCCTCTTCACTGGCCTGTCCTATTTCAACGTCATTTTATGGGGGTCGAAGACAAA
YPS138             : GATCCTTCGTCTGGTATTGGGTACCTGGATTCCTCTTCACTGGCCTGTCCTATTTCAACGTCATTTTATGGGGGTCGAAGACAAG
UWOPS91-917.1      : GATCCTTCGTCTGGTATTGGGTACCTGGATTCCTCTTCACTGGCCTGTCCTATTTCAACGTCATTTTATGGGGGTCGAAGACAAG
N44                : GATCCTTCGTCTGGTATTGGGTACCAGGATTCCTCTTCACTGGCCTGTCCTATTTCAACGTCGTTTTATGGGGGTCAAAGACAAG
CBS432             : GATCCTTCGTCTGGTATTGGGTACCAGGATTCCTCTTCACTGGCCTGTCCTATTTCAACGTCGTTTTATGGGGGTCGAAGACAAG
CEY647             : GATCCTTCGTCTGGTATTGGGTACCTGGATTCCTCTTCACTGGCCTGTCCTATTTCAACGTCATTTTATGGGGGTCGAAGACAAG
CLQCA_20-060       : GATCCTTCGTCTGGTATTGGGTACCTGGATTCCTCTTCACTGGCCTGTCCTATTTCAACGTCATTTTATGGGGGTCGAAGACAAG
CEY650             : GATCGTTCATCTGGTATTGGGTACCTGGATTCCTCTTTACCGGTCTGTCCTATTTCAACGTTATCCTATGGGGGTCGAAGACAAG
CEY653             : GATCGTTCATCTGGTATTGGGTACCTGGATTCCTCTTTACCGGTCTGTCCTATTTCAACGTTATCCTATGGGGGTCGAAGACAAG
CEY649             : GATCGTTCATCTGGTATTGGGTACCTGGATTCCTCTTTACCGGTCTGTCCTATTTCAACGTTATCCTATGGGGGTCGAAGACAAG
YJM1250            : GATCGTTCATCTGGTATTGGGTACCTGGATTCCTCTTTACCGGTCTGTCCTATTTCAACGTTATCCTATGGGGGTCGAAGACAAG
SA.9.4.BR2         : GATCGTTCATCTGGTATTGGGTACCTGGATTCCTCTTTACCGGTCTGTCCTATTTCAACGTTATCCTATGGGGGTCGAAGACAAG
YJM1444            : GATCGTTCATCTGGTATTGGGTACCTGGATTCCTCTTTACCGGTCTGTCCTATTTCAACGTTATCCTATGGGGGTCGAAGACAAG
GLBRCY22_3         : GATCGTTCATCTGGTATTGGGTACCTGGATTCCTCTTTACCGGTCTGTCCTATTTCAACGTTATCCTATGGGGGTCGAAGACAAG
UWOPS87-2421       : GATCGTTCATCTGGTATTGGGTACCTGGATTCCTCTTTACCGGTCTGTCCTATTTCAACGTTATCCTATGGGGGTCGAAGACAAG
YJM653_1b          : GATCGTTCATCTGGTATTGGGTACCTGGATTCCTCTTTACCGGTCTGTCCTATTTCAACGTTATCCTATGGGGGTCGAAGACAAG
YJM681             : GATCGTTCATCTGGTATTGGGTACCTGGATTCCTCTTTACCGGTCTGTCCTATTTCAACGTTATCCTATGGGGGTCGAAGACAAG
EC1118             : GATCGTTCATCTGGTATTGGGTACCTGGATTCCTCTTTACCGGCCTGTCCTATTTCAACGTTATCCTATGGGGGTCGAAGACAAG
S288c              : GATCGTTCATCTGGTATTGGGTACCTGGATTCCTCTTTACCGGTCTGTCCTATTTCAACGTTATCCTATGGGGGTCGAAGACAAG

                     940         *       960         *       980         *      1000         *      1020
H4                 : ACACAATTTCATTGCTAACACAATCTTCGGTACTCAAAGTGGGCTCGGTGCGTTGCCAATCACATTCGACTACACGCAAGTTTCC
RP11.4.14          : ACACAATTTCATTGCTAACACAATCTTCGGTACTCAAAGTGGGCTCGGTGCGTTGCCAATCACATTCGACTACACGCAAGTTTCC
UFRJ50816          : ACACAATTTCATTGCTAACACAATCTTCGGTACTCAAAGTGGGCTCGGTGCGTTGCCAATCACATTCGACTACACGCAAGTTTCC
YPS138             : ACACAATTTCATTGCTAACACAATCTTCGGTACTCAAAGTGGGCTCGGTGCGTTGCCAATCACATTCGACTACACGCAAGTTTCC
UWOPS91-917.1      : ACACAATTTCATTGCTAACACAATCTTCGGTACTCAAAGTGGGCTCGGTGCGTTGCCAATCACATTCGACTACATGCAAGTTTCC
N44                : ACACAATTTCATTGCTAACACAATCTTCGGTACTCAAAGTGGGCTCGGTGCGTTGCCAATCACATTCGACTACACGCAAGTTTCC
CBS432             : ACACAATTTCATTGCTAACACAATCTTCGGTACTCAAAGTGGGCTCGGTGCGTTGCCAATCACATTCGACTACACGCAAGTTTCC
CEY647             : ACACAATTTCATTGCTAACACAATCTTCGGCACTCAAAGTGGACTCGGTGCGTTGCCAATCACATTCGACTACACGCAAGTTTCC
CLQCA_20-060       : ACACAATTTCATTGCTAACACAATCTTCGGTACTCAAAGTGGGCTCGGTGCGTTGCCAATCACATTCGACTACACGCAAGTTTCC
CEY650             : ACACAATTTCATCGCTAACACAATCTTTGGTACTCAAAGTGGTCTCGGTGCCTTGCCAATTACGTTTGACTACACCCAGGTTTCC
CEY653             : ACACAATTTCATCGCTAACACAATCTTTGGTACTCAAAGTGGTCTCGGTGCCTTGCCAATTACGTTTGACTACACCCAGGTTTCC
CEY649             : ACACAATTTCATCGCTAACACAATCTTTGGTACTCAAAGTGGTCTCGGTGCCTTGCCAATTACGTTTGACTACACCCAGGTTTCC
YJM1250            : ACACAATTTCATCGCTAACACAATCTTTGGTACTCAAAGTGGTCTCGGTGCCTTGCCAATTACGTTTGACTACACCCAGGTTTCC
SA.9.4.BR2         : ACACAATTTCATCGCTAACACAATCTTTGGTACTCAAAGTGGTCTCGGTGCCTTGCCAATTACGTTTGACTACACCCAGGTTTCC
YJM1444            : ACACAATTTCATCGCTAACACAATCTTTGGTACTCAAAGTGGTCTCGGTGCCTTGCCAATTACGTTTGACTACACCCAGGTTTCC
GLBRCY22_3         : ACACAATTTCATCGCTAACACAATCTTTGGTACTCAAAGTGGTCTCGGTGCCTTGCCAATTACGTTTGACTACACCCAGGTTTCC
UWOPS87-2421       : ACACAATTTCATCGCTAACACAATCTTTGGTACTCAAAGTGGTCTCGGTGCCTTGCCAATTACGTTTGACTACACCCAGGTTTCC
YJM653_1b          : ACACAATTTCATCGCTAACACAATCTTTGGTACTCAAAGTGGTCTCGGTGCGTTGCCAATTACATTTGACTACACCCAGGTTTCC
YJM681             : ACACAATTTCATCGCTAACACAATCTTTGGTACTCAAAGTGGTCTCGGTGCGTTGCCAATTACATTTGACTACACCCAGGTTTCC
EC1118             : ACACAATTTCATCGCTAACACAATCTTTGGTACTCAAAGTGGTCTCGGTGCGTTGCCAATTACATTTGACTACACCCAGGTTTCT
S288c              : ACACAATTTCATCGCTAACACAATCTTTGGTACTCAAAGTGGTCTCGGTGCGTTGCCAATTACATTTGACTACACCCAGGTTTCC
```

```
                              *        1040          *        1060          *        1080          *        1100
H4                 : CAAGCTATGTCCGGTTCAGTATTCGCTACACCATTCTACGTTTCTGCTAACACGTATGCATCAGTGTTAATATTTTTTGTTATAG
RP11.4.14          : CAAGCTATGTCCGGTTCAGTATTCGCTACACCATTCTACGTTTCTGCTAACACGTATGCATCAGTGTTAATATTTTTTGTTATAG
UFRJ50816          : CAAGCTATGTCCGGTTCAGTATTCGCTACACCATTCTACGTTTCTGCTAACACGTATGCATCAGTGTTAATATTTTTTGTTATAG
YPS138             : CAAGCTATGTCCGGTTCAGTATTCGCTACACCATTCTACGTTTCTGCTAACACATATGCATCAGTGTTAATATTTTTTGTTATTG
UWOPS91-917.1      : CAAGCTATGTCCGGTTCAGTATTTGCTACACCATTCTACGTTTCTGCTAACACGTATGCATCAGTGTTAATATTTTTTGTTATAG
N44                : CAAGCCATGTCCGGTTCTGTATTCGCTACACCATTCTACGTCTCTGCCAACACCTATGCATCAGTGTTAATATTTTTTGTCATAG
CBS432             : CAAGCCATGTCCGGCTCCGTATTCGCTACACCATTCTACGTCTCTGCCAACACCTATGCATCAGTGTTAATATTCTTTGTCATAG
CEY647             : CAAGCTATGTCCGGTTCAGTATTCGCTACACCATTCTACGTTTCTGCTAACACGTATGCATCAGTGTTAATATTTTTTGTTATAG
CLQCA_20-060       : CAAGCTATGTCCGGTTCAGTATTCGCTACACCATTCTACGTTTCTGCTAACACGTATGCATCAGTGTTAATATTTTTTGTTATAG
CEY650             : CAAGCTATGTCCGGTTCAGTATTCGCTACACCATTCTACGTTTCTGCTAACACGTATGCATCAGTGTTAATATTTTTTGTTATAG
CEY653             : CAAGCTATGTCCGGTTCAGTATTCGCTACACCATTCTACGTTTCTGCTAACACGTATGCATCAGTGTTAATATTTTTTGTTATAG
CEY649             : CAAGCTATGTCCGGTTCAGTATTCGCTACACCATTCTACGTTTCTGCTAACACGTATGCATCAGTGTTAATATTTTTTGTTATAG
YJM1250            : CAAGCTATGTCCGGTTCAGTATTCGCTACACCATTCTACGTTTCTGCTAACACGTATGCATCAGTGTTAATATTTTTTGTTATAG
SA.9.4.BR2         : CAAGCTATGTCCGGTTCAGTATTCGCTACACCATTCTACGTTTCTGCTAACACGTATGCATCAGTGTTAATATTTTTTGTTATAG
YJM1444            : CAAGCCATGTCCGGCTCTGTTTTCGCCACACCATTCTACGTCTCCGCCAACACCTATGCATCAGTGTTGATATTCTTCGTCATAG
GLBRCY22_3         : CAAGCCATGTCCGGCTCTGTTTTCGCCACACCATTCTACGTCTCCGCCAACACCTATGCATCAGTGTTGATATTCTTCGTCATAG
UWOPS87-2421       : CAAGCCATGTCCGGCTCTGTTTTCGCCACACCATTCTACGTCTCCGCCAACACCTATGCATCAGTGTTGATATTCTTCGTCATAG
YJM653_1b          : CAAGCCATGTCCGGCTCTGTTTTCGCCACACCATTCTACGTCTCCGCCAACACCTATGCATCAGTGTTGATATTCTTCGTCATAG
YJM681             : CAAGCCATGTCCGGCTCTGTTTTCGCCACACCATTCTACGTCTCCGCCAACACCTATGCATCAGTGTTGATATTCTTCGTCATAG
EC1118             : CAAGCCATGTCCGGCTCTGTTTTCGCCACACCATTCTACGTCTCCGCCAACACCTATGCATCAGTGTTGATATTCTTCGTCATAG
S288c              : CAAGCCATGTCCGGCTCTGTTTTCGCCACACCATTCTACGTCTCCGCCAACACCTATGCATCAGTGTTGATATTCTTCGTCATAG

                          *        1120          *        1140          *        1160          *        1180          *
H4                 : TGCTACCATGCCTCTATTTCACCAATACCTGGTATGCCAAATACATGCCTGTCATTTCAGGTTCCACTTATGACAACACTCAAAA
RP11.4.14          : TGCTACCATGCCTCTATTTCACCAATACCTGGTATGCCAAATACATGCCTGTCATTTCAGGTTCCACTTATGACAACACTCAAAA
UFRJ50816          : TGCTACCATGCCTCTATTTCACCAATACCTGGTATGCCAAATACATGCCTGTCATTTCAGGTTCCACTTATGACAACACTCAAAA
YPS138             : TGCTACCATGCCTCTATTTCACCAACACCTGGTATGCCAAATACATGCCTGTCATTTCAGGTTCCACTTATGACAACACTCAAAA
UWOPS91-917.1      : TGCTACCATGCCTCTATTTCACCAACACCTGGTATGCCAAATACATGCCTGTCATTTCAGGTTCCACTTATGACAACACTCAAAA
N44                : TCCTACCATGCCTCTATTTCACCAACACCTGGTACGCCAAATACATGCCTGTCATTTCAGGTTCCACTTATGACAACACTCAAAA
CBS432             : TCCTACCATGCCTCTATTTCACCAACACCTGGTATGCCAAATACATGCCTGTCATTTCAGGTTCCACTTATGACAACGCTCAAAA
CEY647             : TGCTACCATGCCTCTATTTCACCAATACCTGGTATGCCAAATACATGCCTGTCATTTCAGGTTCCACTTATGACAACACTCAAAA
CLQCA_20-060       : TGCTACCATGTCTCTATTTCACCAACACCTGGTATGCCAAATACATGCCTGTCATTTCAGGTTCCACTTATGACAACACTCAAAA
CEY650             : TGCTACCATGTCTCTATTTCACCAACACCTGGTATGCCAAATACATGCCTGTCATTTCAGGTTCCACTTATGACAACACTCAAAA
CEY653             : TGCTACCATGTCTCTATTTCACCAACACCTGGTATGCCAAATACATGCCTGTCATTTCAGGTTCCACTTATGACAACACTCAAAA
CEY649             : TGCTACCATGTCTCTATTTCACCAACACCTGGTATGCCAAATACATGCCTGTCATTTCAGGTTCCACTTATGACAACACTCAAAA
YJM1250            : TGCTACCATGTCTCTATTTCACCAACACCTGGTATGCCAAATACATGCCTGTCATTTCAGGTTCCACTTATGACAACACTCAAAA
SA.9.4.BR2         : TGCTACCATGCCTCTATTTCACCAACACCTGGTATGCCAAATACATGCCTGTCATTTCAGGTTCCACTTATGACAACACTCAAAA
YJM1444            : TGCTGCCATGTCTTTATTTTACGAATACCTGGTATGCCAAATACATGCCGGTCATTTCAGGTTCTACTTATGACAACACTCAAAA
GLBRCY22_3         : TGCTGCCATGTCTTTATTTTACGAATACCTGGTATGCCAAATACATGCCGGTCATTTCAGGTTCTACTTATGACAACACTCAAAA
UWOPS87-2421       : TGCTGCCATGTCTTTATTTTACGAATACCTGGTATGCCAAATACATGCCGGTCATTTCAGGTTCTACTTATGACAACACTCAAAA
YJM653_1b          : TGCTGCCATGTCTTTATTTTACGAATACCTGGTATGCCAAATACATGCCCGTCATTTCAGGTTCTACTTATGACAACACTCAAAA
YJM681             : TGCTGCCATGTCTTTATTTTACGAATACCTGGTATGCCAAATACATGCCCGTCATTTCAGGTTCTACTTATGACAACACTCAAAA
EC1118             : TGCTGCCATGTCTTTATTTTACGAATACCTGGTATGCCAAATACATGCCCGTCATTTCAGGTTCTACTTATGACAACACTCAAAA
S288c              : TGCTGCCATGTCTTTATTTTACGAATACCTGGTATGCCAAATACATGCCCGTCATTTCAGGTTCTACTTATGACAACACTCAAAA

                           1200          *        1220          *        1240          *        1260          *
H4                 : TAAATACAACGTAACGAAGATTCTTAACGAGGATTATTCCATTAATCTCGAGAAATATAAAGAATACTCACCAGTATTTGTTCCA
RP11.4.14          : TAAATACAACGTAACGAAGATTCTTAACGAGGATTATTCCATTAATCTCGAGAAATATAAAGAATACTCACCAGTATTTGTTCCA
UFRJ50816          : TAAATACAACGTAACGAAGATTCTTAACGAGGATTATTCCATTAATCTCGAGAAATATAAAGAATACTCACCAGTATTTGTTCCA
YPS138             : TAAATACAACGTAACGAAGATTCTTAACGAGGATTATTCCATTAATCTCGAGAAATATAAAGAATACTCACCAGTATTTGTTCCA
UWOPS91-917.1      : TAAATACAACGTAACGAAGATTCTTAACGAGGATTATTCCATTAATCTCGAGAAATATAAAGAATATTCACCAGTATTTGTTCCA
N44                : CAAATACAACGTAACGAAGATTCTTAACGAGGATTATTCCATTAATCTAGAGAAATATAAAGAATATTCGCCAGTATTCGTTCCA
CBS432             : CAAATACAACGTAACGAAGATTCTTAACGAGGATTACTCCATTAATCTCGAGAAATATAAAGAATACTCACCAGTATTTGTTCCA
CEY647             : TAAATACAACGTAACGAAGATTCTTAACGAGGATTATTCCATTAATCTCGAGAAATATAAAGAATACTCACCAGTATTTGTTCCA
CLQCA_20-060       : TAAATACAACGTAACGAAGATTCTTAACGAGGATTATTCCATTAATCTCGAGAAATATAAAGAATACTCACCAGTATTTGTTCCA
CEY650             : TAAATACAACGTAACGAAGATTCTTAACGAGGATTATTCCATTAATCTCGAGAAATATAAAGAATACTCACCAGTATTTGTTCCA
CEY653             : TAAATACAACGTAACGAAGATTCTTAACGAGGATTATTCCATTAATCTCGAGAAATATAAAGAATACTCACCAGTATTTGTTCCA
CEY649             : TAAATACAACGTAACGAAGATTCTTAACGAGGATTATTCCATTAATCTCGAGAAATATAAAGAATACTCACCAGTATTTGTTCCA
YJM1250            : TAAATACAACGTAACGAAGATTCTTAACGAGGATTATTCCATTAATCTCGAGAAATATAAAGAATACTCACCAGTATTTGTTCCA
SA.9.4.BR2         : TAAATACAACGTAACGAAGATTCTTAACGAGGATTATTCCATTAATCTCGAGAAATATAAAGAATACTCACCAGTATTTGTTCCA
YJM1444            : CAAATACAACGTAACAAAGATTCTAAACGAGGATTATTCCATTAATCTTGAGAAATATAAGGAATACTCACCGGTATTCGTTCCA
GLBRCY22_3         : CAAATACAACGTAACAAAGATTCTAAACGAGGATTATTCCATTAATCTTGAGAAATATAAGGAATACTCACCGGTATTCGTTCCA
UWOPS87-2421       : CAAATACAACGTAACAAAGATTCTAAACGAGGATTATTCCATTAATCTTGAGAAATATAAGGAATACTCACCGGTATTCGTTCCA
YJM653_1b          : CAAATACAACGTAACAAAGATTCTTAACGAGGATTATTCCATTAATCTTGAGAAATATAAGGAATACTCACCGGTATTCGTTCCA
YJM681             : CAAATACAACGTAACAAAGATTCTTAACGAGGATTATTCCATTAATCTTGAGAAATATAAGGAATACTCACCGGTATTCGTTCCA
EC1118             : CAAATACAACGTAACAAAGATTCTAAACGAGGATTATTCCATTAATCTTGAGAAATATAAGGAATACTCACCGGTATTCGTTCCA
S288c              : CAAATACAACGTAACAAAGATTCTTAACGAGGATTATTCCATTAATCTTGAGAAATATAAGGAATACTCACCGGTATTCGTTCCA
```

```
                      1280          *        1300          *        1320          *        1340          *        1360
H4              : TTTTCCTATCTTTTGTCGTATGCCTTGAATTTTGCCGCTGTTATTGCTGTGTTTGTTCACTGCTTCTTATACCACGGTAAAGATA
RP11.4.14       : TTTTCCTATCTTTTGTCGTATGCCTTGAATTTTGCCGCTGTTATTGCTGTGTTTGTTCACTGCTTCTTATACCACGGTAAAGATA
UFRJ50816       : TTTTCCTATCTTTTGTCGTATGCCTTGAATTTTGCCGCTGTTATCGCTGTGTTTGTTCACTGCTTCTTATACCACGGTAAAGATA
YPS138          : TTTTCCTATCTTTTGTCGTATGCCTTGAATTTTGCCGCTGTTATCGCTGTGTTTGTTCACTGCTTCTTATACCACGGTAAAGATA
UWOPS91-917.1   : TTTTCCTATCTTTTGTCGTATGCCTTGAATTTTGCCGCTGTTATCGCTGTGTTTGTTCACTGCTTCTTATACCACGGTAAAGATA
N44             : TTTTCCTATCTTTTGTCGTATGCCTTAAATTTTGCCGCTGTTATCGCTGTTTTTGTTCACTGCTTCTTATACCACGGTAAAGATA
CBS432          : TTTTCCTATCTTTTGTCGTATGCCTTGAATTTTGCCGCTGTTATCGCTGTTTTTGTTCACTGCTTCTTATACCATGGTAAAGATA
CEY647          : TTTTCCTATCTTTTGTCGTATGCCTTGAATTTTGCCGCTGTTATTGCTGTGTTTGTTCACTGCTTCTTATACCACGGTAAAGATA
CLQCA_20-060    : TTTTCCTATCTTTTATCGTATGCCTTGAATTTTGCCGCTGTTATCGCTGTGTTTGTTCACTGCTTCTTATACCACGGTAAAGATA
CEY650          : TTTTCCTATCTTTTATCGTATGCCTTGAATTTTGCCGCTGTTATCGCTGTGTTTGTTCACTGCTTCTTATACCACGGTAAAGATA
CEY653          : TTTTCCTATCTTTTATCGTATGCCTTGAATTTTGCCGCTGTTATCGCTGTGTTTGTTCACTGCTTCTTATACCACGGTAAAGATA
CEY649          : TTTTCCTATCTTTTATCGTATGCCTTGAATTTTGCCGCTGTTATCGCTGTGTTTGTTCACTGCTTCTTATACCACGGTAAAGATA
YJM1250         : TTTTCCTATCTTTTATCGTATGCCTTGAATTTTGCCGCTGTTATCGCTGTGTTTGTTCACTGCTTCTTATACCACGGTAAAGATA
SA.9.4.BR2      : TTTTCCTATCTTTTGTCGTATGCCTTGAATTTTGCCGCTGTTATTGCTGTGTTTGTTCACTGCTTCTTATACCACGGTAAAGATA
YJM1444         : TTTTCCTATCTTTTGTCGTATGCTTTAAATTTTGCCGCTGTTATCGCCGTTTTTGTTCACTGCATCTTATACCACGGTAAAGATA
GLBRCY22_3      : TTTTCCTATCTTTTGTCGTATGCTTTAAATTTTGCCGCTGTTATCGCCGTTTTTGTTCACTGCATCTTATACCACGGTAAAGATA
UWOPS87-2421    : TTTTCCTATCTTTTGTCGTATGCTTTAAATTTTGCCGCTGTTATCGCCGTTTTTGTTCACTGCATCTTATACCACGGTAAAGATA
YJM653_1b       : TTTTCCTATCTTTTGTCGTATGCTTTAAATTTTGCCGCTGTTATCGCCGTTTTTGTCCACTGCATCTTATACCACGGTAAAGATA
YJM681          : TTTTCCTATCTTTTGTCGTATGCTTTAAATTTTGCCGCTGTTATCGCCGTTTTTGTCCACTGCATCTTATACCACGGTAAAGATA
EC1118          : TTTTCCTATCTTTTGTCGTATGCTTTAAATTTTGCCGCTGTTATCGCCGTTTTTGTTCACTGCATCTTATACCACGGTAAAGATA
S288c           : TTTTCCTATCTTTTGTCGTATGCTTTAAATTTTGCCGCTGTTATCGCCGTTTTTGTCCACTGCATCTTATACCACGGTAAAGATA

                             *        1380          *        1400          *        1420          *        1440
H4              : TTGTCGCCAAGTTTAAAGACCGTAAAAATGGTGGTACCGATATTCACATGAGAATCTACTCAAAAAACTATAAGGACTGTCCCGA
RP11.4.14       : TTGTCGCCAAGTTTAAAGACCGTAAAAATGGTGGTACCGATATTCACATGAGAATCTACTCAAAAAACTATAAGGACTGTCCCGA
UFRJ50816       : TTGTCGCCAAGTTTAAAGACCGTAAAAATGGTGGTACCGATATTCACATGAGAATCTACTCAAAAAACTATAAGGACTGTCCCGA
YPS138          : TTGTCGCCAAGTTTAAAGACCGTAAAAATGGTGGTACCGATATTCACATGAGAATATACTCAAAAAACTATAAGGACTGTCCCGA
UWOPS91-917.1   : TTGTCGCCAAGTTTAAAGACCGTAAAAATGGTGGTACCGATATTCACATGAGAATCTACTCAAAAAACTATAAGGACTGTCCCGA
N44             : TTGTCGCCAAGTTTAAAGACCGTAAAAACGGTGGTACCGACATTCACATGAGAATCTACTCAAAAAACTATAAGGACTGTCCCGA
CBS432          : TTGTCGCCAAGTTTAAAGACCGTAAAAATGGTGGTACCGACATTCACATGAGAATCTACTCAAAAAACTACAAGGACTGTCCCGA
CEY647          : TTGTCGCCAAGTTTAAAGACCGTAAAAATGGTGGTACCGATATTCACATGAGAATCTACTCAAAAAACTATAAGGACTGTCCCGA
CLQCA_20-060    : TTGTCGCCAAGTTTAAAGACCGTAAAAATGGTGGTACCGATATTCACATGAGAATCTACTCAAAAAACTATAAGGACTGTCCCGA
CEY650          : TTGTCGCCAAGTTTAAAGACCGTAAAAATGGTGGTACCGATATTCACATGAGAATCTACTCAAAAAACTATAAGGACTGTCCCGA
CEY653          : TTGTCGCCAAGTTTAAAGACCGTAAAAATGGTGGTACCGATATTCACATGAGAATCTACTCAAAAAACTATAAGGACTGTCCCGA
CEY649          : TTGTCGCCAAGTTTAAAGACCGTAAAAATGGTGGTACCGATATTCACATGAGAATCTACTCAAAAAACTATAAGGACTGTCCCGA
YJM1250         : TTGTCGCCAAGTTTAAAGACCGTAAAAATGGTGGTACCGATATTCACATGAGAATCTACTCAAAAAACTATAAGGACTGTCCCGA
SA.9.4.BR2      : TTGTCGCCAAGTTTAAAGACCGTAAAAATGGTGGTACCGATATTCACATGAGAATCTACTCAAAAAACTATAAGGACTGTCCCGA
YJM1444         : TTGTCGCCAAGTTTAAAGACCGTAAAAATGGTGGCACTGACATTCACATGAGGATCTACTCCAAGAACTATAAGGATTGTCCCGA
GLBRCY22_3      : TTGTCGCCAAGTTTAAAGACCGTAAAAATGGTGGCACTGACATTCACATGAGGATCTACTCCAAGAACTATAAGGATTGTCCCGA
UWOPS87-2421    : TTGTCGCCAAGTTTAAAGACCGTAAAAATGGTGGCACTGACATTCACATGAGGATCTACTCCAAGAACTATAAGGATTGTCCCGA
YJM653_1b       : TTGTCGCCAAGTTAAAAGACCGTAAAAATGGTGGCACTGACATTCACATGAGAATCTACTCCAAGAACTATAAGGATTGTCCCGA
YJM681          : TTGTCGCCAAGTTAAAAGACCGTAAAAATGGTGGCACTGACATTCACATGAGAATCTACTCCAAGAACTATAAGGATTGTCCCGA
EC1118          : TTGTCGCCAAGTTTAAAGACCGTAAAAATGGTGGCACTGACATTCACATGAGGATCTACTCCAAGAACTATAAGGATTGTCCCGA
S288c           : TTGTCGCCAAGTTTAAAGACCGTAAAAATGGTGGCACTGACATTCACATGAGAATCTACTCCAAGAACTATAAGGATTGTCCCGA

                      *        1460          *        1480          *        1500          *        1520          *
H4              : TTGGTGGTATTTGCTCTTGCAAATTGTCATGATCGGTTTAGGGTTTGTAGCAGTATGCTGCTTCGATACTAAGTTCCCAGCCTGG
RP11.4.14       : TTGGTGGTATTTGCTCTTGCAAATTGTCATGATCGGTTTAGGGTTTGTAGCAGTATGCTGCTTCGATACTAAGTTCCCAGCCTGG
UFRJ50816       : TTGGTGGTATTTGCTCTTGCAAATTGTCATGATCGGTTTAGGGTTTGTAGCAGTATGCTGCTTTGATACTAAGTTCCCAGCCTGG
YPS138          : TTGGTGGTATTTGCTCTTGCAAATTGTCATGATCGGTTTAGGGTTTGTAGCAGTATGCTGCTTCGATACTAAGTTCCCAGCCTGG
UWOPS91-917.1   : TTGGTGGTATTTGCTCTTGCAAATTGTCATGATCGGTTTAGGGTTTGTAGCAGTATGCTGCTTCGATACTAAGTTCCCAGCCTGG
N44             : TTGGTGGTATTTGCTCTTGCAAATTGTCATGATCGGGTTAGGGTTTGTGGCAGTATGCTGCTTCGATACTAAATTCCCAGCCTGG
CBS432          : TTGGTGGTATTTGCTCTTGCAAATTGTCATGATCGGTTTAGGGTTTGTGGCAGTATGCTGCTTCGATACTAAGTTCCCAGCCTGG
CEY647          : TTGGTGGTATTTGCTCTTGCAAATTGTCATGATCGGTTTAGGGTTTGTAGCAGTATGCTGCTTCGATACTAAGTTCCCAGCCTGG
CLQCA_20-060    : TTGGTGGTATTTGCTCTTGCAAATTGTCATGATCGGTTTAGGGTTTGTAGCAGTATGCTGCTTCGATACTAAGTTCCCAGCCTGG
CEY650          : TTGGTGGTATTTGCTCTTGCAAATTGTCATGATCGGTTTAGGGTTTGTAGCAGTATGCTGCTTCGATACTAAGTTCCCAGCCTGG
CEY653          : TTGGTGGTATTTGCTCTTGCAAATTGTCATGATCGGTTTAGGGTTTGTAGCAGTATGCTGCTTCGATACTAAGTTCCCAGCCTGG
CEY649          : TTGGTGGTATTTGCTCTTGCAAATTGTCATGATCGGTTTAGGGTTTGTAGCAGTATGCTGCTTCGATACTAAGTTCCCAGCCTGG
YJM1250         : TTGGTGGTATTTGCTCTTGCAAATTGTCATGATCGGTTTAGGGTTTGTAGCAGTATGCTGCTTCGATACTAAGTTCCCAGCCTGG
SA.9.4.BR2      : TTGGTGGTATTTGCTCTTGCAAATTGTCATGATCGGTTTAGGGTTTGTAGCAGTATGCTGCTTCGATACTAAGTTCCCAGCCTGG
YJM1444         : TTGGTGGTATTTACTTTTGCAGATTGTCATGATCGGTTTAGGATTTGTAGCAGTGTGCTGTTTCGATACTAAGTTCCCAGCTTGG
GLBRCY22_3      : TTGGTGGTATTTACTTTTGCAGATTGTCATGATCGGTTTAGGATTTGTAGCAGTGTGCTGTTTCGATACTAAGTTCCCAGCTTGG
UWOPS87-2421    : TTGGTGGTATTTACTTTTGCAGATTGTCATGATCGGTTTAGGATTTGTAGCAGTGTGCTGTTTCGATACTAAGTTCCCAGCTTGG
YJM653_1b       : TTGGTGGTATTTACTTTTGCAGATTGTCATGATCGGTTTAGGATTTGTAGCAGTGTGCTGTTTCGATACTAAGTTCCCAGCTTGG
YJM681          : TTGGTGGTATTTACTTTTGCAGATTGTCATGATCGGTTTAGGATTTGTAGCAGTGTGCTGTTTCGACACTAAGTTCCCAGCTTGG
EC1118          : TTGGTGGTATTTACTTTTGCAGATTGTCATGATCGGTTTAGGATTTGTAGCAGTGTGCTGTTTCGATACTAAGTTCCCAGCTTGG
S288c           : TTGGTGGTATTTACTTTTGCAGATTGTCATGATCGGTTTAGGATTTGTAGCAGTGTGCTGTTTCGATACTAAGTTCCCAGCTTGG
```

```
                       1540         *        1560         *        1580         *        1600         *
H4                 : GCATTTGTTATTGCCATATTAATTTCCCTTGTAAATTTCATTCCACAAGGTATTTTGGAAGCAATGACTAATCAACACGTTGGCT
RP11.4.14          : GCATTTGTTATTGCCATATTAATTTCCCTTGTAAATTTCATTCCACAAGGTATTTTGGAAGCAATGACTAATCAACACGTTGGCT
UFRJ50816          : GCATTTGTTCTTGCCATATTAATTTCCCTTGTAAATTTCATTCCACAAGGTATTTTGGAAGCAATGACTAATCAACACGTTGGCT
YPS138             : GCATTTGTTATTGCCATATTAATTTCCCTTGTAAATTTCATTCCACAAGGTATTTTGGAAGCAATGACTAATCAACACGTTGGCT
UWOPS91-917.1      : GCATTTGTTATTGCCATATTAATTTCCCTTGTAAATTTCATTCCACAAGGTATTTTGGAAGCAATGACTAATCAACACGTTGGCT
N44                : GCATTTGTTATTGCCATATTAATTTCCCTTGTAAATTTCATTCCACAAGGTATTTTGGAAGCAATGACTAATCAACACGTTGGTT
CBS432             : GCATTTGTTATTGCCATATTAATTTCCCTTGTAAATTTCATTCCACAAGGTATATTGGAAGCAATGACTAATCAACACGTTGGTT
CEY647             : GCATTTGTTATTGCCATATTAATTTCCCTTGTAAATTTCATTCCACAAGGTATTTTGGAAGCAATGACTAATCAACACGTTGGCT
CLQCA_20-060       : GCATATGTTATTGCCATATTAATTTCCCTTGTAAATTTCATTCCACAAGGTATTTTGGAAGCAATGACTAATCAACACGTTGGCT
CEY650             : GCATATGTTATTGCCATATTAATTTCCCTTGTAAATTTCATTCCACAAGGTATTTTGGAAGCAATGACTAATCAACACGTTGGCT
CEY653             : GCATATGTTATTGCCATATTAATTTCCCTTGTAAATTTCATTCCACAAGGTATTTTGGAAGCAATGACTAATCAACACGTTGGCT
CEY649             : GCATATGTTATTGCCATATTAATTTCCCTTGTAAATTTCATTCCACAAGGTATTTTGGAAGCAATGACTAATCAACACGTTGGCT
YJM1250            : GCATATGTTATTGCCATATTAATTTCCCTTGTAAATTTCATTCCACAAGGTATTTTGGAAGCAATGACTAATCAACACGTTGGCT
SA.9.4.BR2         : GCATTTGTTATTGCCATATTAATTTCCCTTGTAAATTTCATTCCACAAGGTATTTTGGAAGCAATGACTAATCAACACGTTGGCT
YJM1444            : GCATTTGTTATTGCAATATTAATTTCCCTTGTAAATTTTATCCCGCAAGGTATCTTGGAAGCAATGACTAACCAACACGTAGGCT
GLBRCY22_3         : GCATTTGTTATTGCAATATTAATTTCCCTTGTAAATTTTATCCCGCAAGGTATCTTGGAAGCAATGACTAACCAACACGTAGGCT
UWOPS87-2421       : GCATTTGTTATTGCAATATTAATTTCCCTTGTAAATTTTATCCCGCAAGGTATCTTGGAAGCAATGACTAACCAACACGTAGGCT
YJM653_1b          : GCATTTGTTATTGCAATATTAATTTCCCTTGTAAATTTTATCCCGCAAGGTATCTTGGAAGCAATGACTAACCAACACGTAGGCT
YJM681             : GCATTTGTTATTGCAATATTAATTTCCCTTGTAAATTTCATCCCGCAAGGTATCTTGGAAGCAATGACTAACCAACACGTAGGTT
EC1118             : GCATTTGTTATTGCAATATTAATTTCCCTTGTAAATTTTATCCCGCAAGGTATCTTGGAAGCAATGACTAACCAACACGTAGGTT
S288c              : GCATTTGTTATTGCAATATTAATTTCCCTTGTAAATTTCATCCCGCAAGGTATCTTGGAAGCAATGACTAACCAACACGTAGGTT

                   1620         *        1640         *        1660         *        1680         *        1700
H4                 : TGAATATTATCACAGAATTGATTTGTGGTTATATGCTGCCTTTAAGACCAATGGCAAACTTGTTATTTAAACTATACGGATTCAT
RP11.4.14          : TGAATATTATCACAGAATTGATTTGTGGTTATATGCTGCCTTTAAGACCAATGGCAAACTTGTTATTCAAATTATACGGATTCAT
UFRJ50816          : TGAATATTATCACAGAATTGATTTGTGGTTATATGCTGCCTTTAAGACCAATGGCAAACTTGTTATTCAAACTATACGGATTCAT
YPS138             : TGAATATTATCACAGAATTGATTTGTGGTTATATGCTGCCTTTAAGACCAATGGCAAACTTGTTATTCAAACTATACGGATTCAT
UWOPS91-917.1      : TGAATATTATCACAGAATTGATTTGTGGTTATATGCTGCCTTTAAGACCAATGGCAAACTTGTTATTCAAACTATACGGATTCAT
N44                : TGAATATTATCACAGAATTGATTTGTGGTTATATGCTGCCTTTAAGGCCAATGGCAAACTTGTTATTCAAACTATACGGATTCAT
CBS432             : TGAATATTATCACAGAATTGATTTGTGGTTATATGCTGCCTTTAAGACCAATGGCAAACTTGTTATTCAAACTATACGGATTCAT
CEY647             : TGAATATTATCACAGAATTGATTTGTGGTTATATGCTGCCTTTAAGACCAATGGCAAACTTGTTATTCAAACTATACGGATTCAT
CLQCA_20-060       : TGAATATTATCACAGAATTGATTTGTGGTTATATGCTGCCTTTAAGACCAATGGCAAACTTGTTATTCAAATTATACGGATTCAT
CEY650             : TGAATATTATCACAGAATTGATTTGTGGTTATATGCTGCCTTTAAGACCAATGGCAAACTTGTTATTCAAATTATACGGATTCAT
CEY653             : TGAATATTATCACAGAATTGATTTGTGGTTATATGCTGCCTTTAAGACCAATGGCAAACTTGTTATTCAAATTATACGGATTCAT
CEY649             : TGAATATTATCACAGAATTGATTTGTGGTTATATGCTGCCTTTAAGACCAATGGCAAACTTGTTATTCAAATTATACGGATTCAT
YJM1250            : TGAATATTATCACAGAATTGATTTGTGGTTATATGCTGCCTTTAAGACCAATGGCAAACTTGTTATTCAAATTATACGGATTCAT
SA.9.4.BR2         : TGAATATTATCACAGAATTGATTTGTGGTTATATGCTGCCTTTAAGACCAATGGCAAACTTGTTATTTAAACTATACGGATTCAT
YJM1444            : TGAATATTATCACAGAATTGATCTGCGGTTATATGCTGCCTTTAAGACCAATGGCAAACTTATTATTCAAGCTATACGGATTTAT
GLBRCY22_3         : TGAATATTATCACAGAATTGATCTGCGGTTATATGCTGCCTTTAAGACCAATGGCAAACTTATTATTCAAGCTATACGGATTTAT
UWOPS87-2421       : TGAATATTATCACAGAATTGATCTGCGGTTATATGCTGCCTTTAAGACCAATGGCAAACTTATTATTCAAGCTATACGGATTTAT
YJM653_1b          : TGAATATTATCACAGAATTGATCTGCGGTTATATGCTGCCTTTAAGACCAATGGCAAACTTATTATTCAAGCTATACGGATTTAT
YJM681             : TGAATATTATCACAGAATTGATCTGCGGTTATATGCTGCCTTTAAGACCAATGGCAAACTTATTATTCAAGCTATACGGATTTAT
EC1118             : TGAATATTATCACAGAATTGATCTGCGGTTATATGCTGCCTTTAAGACCAATGGCAAACTTATTATTCAAGCTATACGGATTTAT
S288c              : TGAATATTATCACAGAATTGATCTGCGGTTATATGCTGCCTTTAAGACCAATGGCAAACTTATTATTCAAGCTATACGGATTTAT

                           *        1720         *        1740         *        1760         *        1780
H4                 : CGTCATGAGACAAGGTTTAAATCTAAGTAGAGATTTGAAATTGGCTATGTATATGAAAGTTTCCCCCCGTCTGATTTTTGCCGTT
RP11.4.14          : CGTCATGAGACAAGGTTTAAATCTAAGTAGAGATTTGAAATTGGCTATGTATATGAAAGTTTCCCCCCGTCTGATTTTTGCCGTT
UFRJ50816          : CGTCATGAGACAAGGTTTAAATCTAAGTAGAGATTTGAAATTGGCTATGTACATGAAAGTTTCCCCCCGTCTGATTTTTGCCGTT
YPS138             : CGTCATGAGACAAGGTTTAAATCTAAGTAGAGATTTGAAATTGGCTATGTACATGAAAGTTTCCCCCCGTCTGATTTTTGCCGTT
UWOPS91-917.1      : CGTCATGAGACAAGGTTTAAATCTAAGTAGAGATTTGAAATTGGCTATGTACATGAAAGTTTCCCCCCGTCTGATTTTTGCCGTT
N44                : CGTCATGAGACAAGGTTTAAACCTGAGTAGAGATTTGAAATTGGCTATGTACATGAAAGTTTCCCCTCGTCTGATTTTTGCCGTT
CBS432             : CGTCATGAGACAAGGTTTAAATCTGAGTAGAGATTTGAAATTGGCTATGTACATGAAAGTTTCCCCCCGTCTGATTTTTGCCATT
CEY647             : CGTCATGAGACAAGGTTTAAATCTAAGTAGAGATTTGAAATTGGCTATGTATATGAAAGTTTCCCCTCGTTTGATCTTTGCCGTT
CLQCA_20-060       : CGTCATGAGACAAGGTTTAAATCTAAGTAGAGATTTGAAATTGGCTATGTATATGAAAGTTTCCCCCCGTCTGATTTTTGCCGTT
CEY650             : CGTCATGAGACAAGGTTTAAATCTAAGTAGAGATTTGAAATTGGCTATGTATATGAAAGTTTCCCCCCGTCTGATTTTTGCCGTT
CEY653             : CGTCATGAGACAAGGTTTAAATCTAAGTAGAGATTTGAAATTGGCTATGTATATGAAAGTTTCCCCCCGTCTGATTTTTGCCGTT
CEY649             : CGTCATGAGACAAGGTTTAAATCTAAGTAGAGATTTGAAATTGGCTATGTATATGAAAGTTTCCCCCCGTCTGATTTTTGCCGTT
YJM1250            : CGTCATGAGACAAGGTTTAAATCTAAGTAGAGATTTGAAATTGGCTATGTATATGAAAGTTTCCCCCCGTCTGATTTTTGCCGTT
SA.9.4.BR2         : CGTCATGAGACAAGGTTTAAATCTAAGTAGAGATTTGAAATTGGCTATGTATATGAAAGTTTCCCCCCGTCTGATTTTTGCCGTT
YJM1444            : TGTCATGAGACAAGGCTTGAATTTGAGTAGAGATTTGAAATTAGCCATGTACATGAAGGTTTCCCCTCGTTTGATCTTTGCCGTT
GLBRCY22_3         : TGTCATGAGACAAGGCTTGAATTTGAGTAGAGATTTGAAATTAGCCATGTACATGAAGGTTTCCCCTCGTTTGATCTTTGCCGTT
UWOPS87-2421       : TGTCATGAGACAAGGCTTGAATTTGAGTAGAGATTTGAAATTAGCCATGTACATGAAGGTTTCCCCTCGTTTGATCTTTGCCGTT
YJM653_1b          : TGTCATGAGACAAGGCTTGAATTTGAGTAGAGATTTGAAATTAGCCATGTACATGAAGGTTTCCCCTCGTTTGATCTTTGCCGTT
YJM681             : TGTCATGAGACAAGGCTTGAATTTGAGTAGAGATTTGAAATTAGCCATGTACATGAAGGTTTCCCCTCGTTTGATCTTTGCCGTT
EC1118             : TGTCATGAGACAAGGCTTGAATTTGAGTAGAGATTTGAAATTAGCCATGTACATGAAGGTTTCCCCTCGTTTGATCTTTGCCGTC
S288c              : TGTCATGAGACAAGGCTTGAATTTGAGTAGAGATTTGAAATTAGCCATGTACATGAAGGTTTCCCCTCGTTTGATCTTTGCCGTT
```

```
                     *      1800       *      1820       *      1840       *      1860       *
H4             : CAAATCTATGCCACTATAATATCTGGTATGGTTAACGTCGGTGTCCAAGAATGGATGATGCATAATATCGATGGCTTGTGTACCA
RP11.4.14      : CAAATCTATGCCACTATAATATCTGGTATGGTTAACGTCGGTGTCCAAGAATGGATGATGCATAATATCGATGGCTTGTGTACCA
UFRJ50816      : CAAATCTATGCCACTATAATATCTGGTATGGTTAACGTCGGTGTCCAAGAATGGATGATGCATAATATCGATGGCTTGTGTACCA
YPS138         : CAAATCTATGCCACTATAATATCTGGTATGGTTAACGTCGGTGTCCAAGAATGGATGATGCATAATATCGATGGCTTGTGTACCA
UWOPS91-917.1  : CAAATCTATGCCACTATAATATCTGGTATGGTTAACGTTGGTGTCCAAGAATGGATGATGCATAATATCGATGGCTTGTGTACCA
N44            : CAAATCTATGCCACTATCATATCTGGTATGGTTAACGTTGGTGTCCAGGAATGGATGATGCATAATATCGATGGCTTGTGTACCA
CBS432         : CAAATTTATGCCACTATCATATCTGGTATGGTTAACGTTGGTGTACAGGAATGGATGATGCATAATATCGATGGCTTGTGTACCA
CEY647         : CAAATCTATGCCACTATCATATCAGGTATGGTTAACGTTGGTGTCCAGGAATGGATGATGCATAATATCGATGGCTTGTGTACCA
CLQCA_20-060   : CAAATCTATGCCACTATAATATCTGGTATGGTTAACGTCGGTGTCCAAGAATGGATGATGCATAATATCGATGGCTTGTGTACCA
CEY650         : CAAATCTATGCCACTATAATATCTGGTATGGTTAACGTCGGTGTCCAAGAATGGATGATGCATAATATCGATGGCTTGTGTACCA
CEY653         : CAAATCTATGCCACTATAATATCTGGTATGGTTAACGTCGGTGTCCAAGAATGGATGATGCATAATATCGATGGCTTGTGTACCA
CEY649         : CAAATCTATGCCACTATAATATCTGGTATGGTTAACGTCGGTGTCCAAGAATGGATGATGCATAATATCGATGGCTTGTGTACCA
YJM1250        : CAAATCTATGCCACTATAATATCTGGTATGGTTAACGTCGGTGTCCAAGAATGGATGATGCATAATATCGATGGCTTGTGTACCA
SA.9.4.BR2     : CAAATCTATGCCACTATAATATCTGGTATGGTTAACGTCGGTGTCCAAGAATGGATGATGCATAATATCGATGGCTTGTGTACCA
YJM1444        : CAAATCTATGCCACTATCATATCAGGTATGGTTAACGTTGGTGTCCAGGAATGGATGATGCATAATATCGATGGCTTATGTACCA
GLBRCY22_3     : CAAATCTATGCCACTATCATATCAGGTATGGTTAACGTTGGTGTCCAGGAATGGATGATGCATAATATCGATGGCTTATGTACCA
UWOPS87-2421   : CAAATCTATGCCACTATCATATCAGGTATGGTTAACGTTGGTGTCCAGGAATGGATGATGCATAATATCGATGGCTTATGTACCA
YJM653_1b      : CAAATCTATGCCACTATCATATCAGGTATGGTTAACGTTGGTGTCCAGGAATGGATGATGCATAATATCGATGGCTTATGTACCA
YJM681         : CAAATCTATGCCACTATCATATCAGGCATGGTTAACGTTGGTGTCCAGGAATGGATGATGCATAATATCGATGGCTTATGTACCA
EC1118         : CAAATCTATGCCACTATCATATCAGGTATGGTTAACGTTGGTGTCCAGGAATGGATGATGCATAATATCGATGGCTTATGTACCA
S288c          : CAAATCTATGCCACTATCATATCAGGCATGGTTAACGTTGGTGTCCAGGAATGGATGATGCATAATATCGATGGCTTATGTACCA

                   1880       *      1900       *      1920       *      1940       *
H4             : CTGATCAACCAAATGGGTTCACTTGTGCAAATGGTCGCACAGTTTTCAACGCCTCCATCATCTGGTCTTTGCCAAAGTATCTTTT
RP11.4.14      : CTGATCAACCAAATGGGTTCACTTGTGCAAATGGTCGCACAGTTTTCAACGCCTCCATCATCTGGTCTTTGCCAAAGTATCTTTT
UFRJ50816      : CTGATCAACCAAATGGGTTCACTTGTGCAAATGGTCGCACAGTTTTCAACGCCTCCATCATCTGGTCTTTGCCAAAGTATCTTTT
YPS138         : CTGATCAACCAAATGGGTTCACTTGTGCAAATGGTCGCACAGTTTTCAACGCCTCCATCATCTGGTCTTTGCCAAAGTATCTTTT
UWOPS91-917.1  : CTGATCAACCAAATGGGTTCACTTGTGCAAATGGTCGCACAGTTTTCAACGCCTCCATCATCTGGTCTTTGCCAAAGTATCTTTT
N44            : CTGATCAGCCAAACGGGTTCACTTGTGCAAATGGTCGCACAGTTTTCAATGCCTCCATCATATGGTCTTTGCCAAAGTATCTGTT
CBS432         : CTGATCAGCCAAACGGGTTCACTTGTGCAAATGGTCGCACAGTTTTCAACGCCTCTATCATCTGGTCTTTGCCAAAGTATCTTTT
CEY647         : CTGATCAACCAAATGGGTTCACTTGTGCAAATGGTCGCACAGTTTTCAACGCCTCCATCATCTGGTCTTTGCCAAAGTATCTTTT
CLQCA_20-060   : CTGATCAACCAAATGGGTTCACTTGTGCAAATGGTCGCACAGTTTTCAATGCCTCCATCATCTGGTCTTTGCCAAAGTATCTTTT
CEY650         : CTGATCAACCAAATGGGTTCACTTGTGCAAATGGTCGCACAGTTTTCAATGCCTCCATCATCTGGTCTTTGCCAAAGTATCTTTT
CEY653         : CTGATCAACCAAATGGGTTCACTTGTGCAAATGGTCGCACAGTTTTCAATGCCTCCATCATCTGGTCTTTGCCAAAGTATCTTTT
CEY649         : CTGATCAACCAAATGGGTTCACTTGTGCAAATGGTCGCACAGTTTTCAATGCCTCCATCATCTGGTCTTTGCCAAAGTATCTTTT
YJM1250        : CTGATCAACCAAATGGGTTCACTTGTGCAAATGGTCGCACAGTTTTCAATGCCTCCATCATCTGGTCTTTGCCAAAGTATCTTTT
SA.9.4.BR2     : CTGATCAACCAAATGGGTTCACTTGTGCAAATGGTCGCACAGTTTTCAACGCCTCCATCATCTGGTCTTTGCCAAAGTATCTTTT
YJM1444        : CCGATCAACCAAATGGCTTCACTTGTGCTAATGGTCGCACGGTTTTCAATGCTTCCATTATCTGGTCTTTGCCAAAGTATCTTTT
GLBRCY22_3     : CCGATCAACCAAATGGCTTCACTTGTGCTAATGGTCGCACGGTTTTCAATGCTTCCATTATCTGGTCTTTGCCAAAGTATCTTTT
UWOPS87-2421   : CCGATCAACCAAATGGCTTCACTTGTGCTAATGGTCGCACGGTTTTCAATGCTTCCATTATCTGGTCTTTGCCAAAGTATCTTTT
YJM653_1b      : CCGATCAACCAAATGGCTTCACTTGTGCTAATGGTCGCACGGTTTTCAATGCTTCCATTATCTGGTCTTTGCCAAAGTATCTTTT
YJM681         : CCGATCAACCAAATGGCTTCACTTGTGCTAATGGTCGCACGGTTTTCAATGCTTCCATTATCTGGTCTTTGCCAAAGTATCTTTT
EC1118         : CCGATCAACCAAATGGCTTCACTTGTGCTAATGGTCGCACGGTTTTCAATGCTTCCATTATCTGGTCTTTGCCAAAGTATCTTTT
S288c          : CCGATCAACCAAATGGCTTCACTTGTGCTAATGGTCGCACGGTTTTCAATGCTTCCATTATCTGGTCTTTGCCAAAGTATCTTTT

                    1960       *      1980       *      2000       *      2020       *      2040
H4             : CTCATCAGGGCGTATTTATAGTCCGCTAATGTGGTTCTTCTTGATTGGTTTGCTATTCCCACTAGTGATTTATGCCATTCAATGG
RP11.4.14      : CTCATCAGGGCGTATTTATAGTCCGCTAATGTGGTTCTTCTTGATTGGTTTGCTATTCCCACTAGTGATTTATGCCATTCAATGG
UFRJ50816      : CTCATCAGGGCGTATTTATAGTCCGCTAATGTGGTTCTTTTTGATTGGTTTGCTATTCCCACTAGTGATTTATGCCATTCAATGG
YPS138         : CTCATCAGGGCGTATTTATAGTCCGCTAATGTGGTTCTTTTTGATTGGTTTGCTATTCCCACTAGTGATTTATGCCATTCAATGG
UWOPS91-917.1  : CTCATCAGGGCGTATTTATAGTCCGCTAATGTGGTTCTTCTTGATTGGTTTGCTATTCCCACTAGTGATTTATGCCATTCAATGG
N44            : CTCATCAGGGCGCATTTATAGTCCGCTAATGTGGTTTTTCTTGATTGGTTTGCTATTCCCATTAGTGGTTTATGCAATTCAATGG
CBS432         : CTCATCAGGGCGTATTTATAGTCCGCTAATGTGGTTCTTCTTGATTGGTTTGCTATTTCCACTAGTGGTTTATGCCATTCAATGG
CEY647         : CTCATCAGGGCGTATTTATAGTCCGCTAATGTGGTTCTTCTTGATTGGTTTGCTATTCCCACTAGTGATTTATGCCATTCAATGG
CLQCA_20-060   : CTCATCAGGGCGTATTTATAGTCCGCTAATGTGGTTCTTCTTGATTGGTTTGCTATTCCCACTAGTGATTTACGCCATTCAATGG
CEY650         : CTCATCAGGGCGTATTTATAGTCCGCTAATGTGGTTCTTCTTGATTGGTTTGCTATTCCCACTAGTGATTTACGCCATTCAATGG
CEY653         : CTCATCAGGGCGTATTTATAGTCCGCTAATGTGGTTCTTCTTGATTGGTTTGCTATTCCCACTAGTGATTTACGCCATTCAATGG
CEY649         : CTCATCAGGGCGTATTTATAGTCCGCTAATGTGGTTCTTCTTGATTGGTTTGCTATTCCCACTAGTGATTTACGCCATTCAATGG
YJM1250        : CTCATCAGGGCGTATTTATAGTCCGCTAATGTGGTTCTTCTTGATTGGTTTGCTATTCCCACTAGTGATTTACGCCATTCAATGG
SA.9.4.BR2     : CTCATCAGGGCGTATTTATAGTCCGCTAATGTGGTTCTTCTTGATTGGTTTGCTATTCCCACTAGTGATTTATGCCATTCAATGG
YJM1444        : CTCATCAGGGCGCATTTATAATCCGCTGATGTGGTTCTTCTTGATTGGTTTGCTATTCCCACTAGCCGTTTATGCTGTTCAATGG
GLBRCY22_3     : CTCATCAGGGCGCATTTATAATCCGCTGATGTGGTTCTTCTTGATTGGTTTGCTATTCCCACTAGCCGTTTATGCTGTTCAATGG
UWOPS87-2421   : CTCATCAGGGCGCATTTATAATCCGCTGATGTGGTTCTTCTTGATTGGTTTGCTATTCCCACTAGCCGTTTATGCTGTTCAATGG
YJM653_1b      : CTCATCAGGGCGCATTTATAATCCGCTGATGTGGTTCTTCTTGATTGGTTTGCTATTCCCACTAGCCGTTTATGCTGTTCAATGG
YJM681         : CTCATCAGGGCGCATTTATAATCCGCTGATGTGGTTCTTCTTGATTGGTTTGCTATTCCCACTAGCCGTTTATGCTGTTCAATGG
EC1118         : CTCATCAGGGCGCATTTATAATCCGCTGATGTGGTTCTTCTTGATTGGTTTGCTATTCCCACTAGCCGTTTATGCTGTTCAATGG
S288c          : CTCATCAGGGCGCATTTATAATCCGCTGATGTGGTTCTTCTTGATTGGTTTGCTATTCCCACTAGCCGTTTATGCTGTTCAATGG
```

```
                        *        2060         *        2080         *        2100         *        2120
H4              : AAATTCCCTAACTTTAAATTTGCCAAGCACATCCACACTCCCGTATTTTTCACAGGGCCAGGTAATATCCCCCCAAGTACACCTT
RP11.4.14       : AAATTCCCTAACTTTAAATTTGCCAAGCACATCCACACTCCCGTATTTTTCACAGGGCCAGGTAATATCCCCCCAAGTACACCTT
UFRJ50816       : AAATTCCCTAACTTTAAATTTGCCAAGCACATCCACACTCCCGTATTTTTCACAGGGCCAGGTAATATCCCCCCAAGTACACCTT
YPS138          : AAATTCCCTAACTTTAAATTTGCCAAGCACATCCACACTCCCGTATTTTTCACAGGGCCAGGTAATATCCCCCCAAGTACACCTT
UWOPS91-917.1   : AAATTCCCTAACTTTAAATTTGCCAAGCACATCCACACTCCCGTATTTTTCACAGGACCAGGTAATATCCCCCCAAGTACACCTT
N44             : AAGTTCCCTAACTTTAAATTCGCTAAGCACATACACACTCCCGTATTTTTCACAGGCCCAGGTAATATCCCCCCAAGCACACCTT
CBS432          : AAATTCCCTAACTTTAAATTCGCTAAGCACATTCACACTCCCGTATTTTTCACAGGCCCAGGTAATATCCCCCCAAGCACACCTT
CEY647          : AAATTCCCTAACTTTAAATTTGCCAAGCACATCCACACTCCCGTATTTTTCACAGGGCCAGGTAATATTCCCCCAAGTACACCTT
CLQCA_20-060    : AAATTCCCTAACTTTAAATTTGCCAAGCACATCCACACTCCCGTATTTTTCACAGGGCCAGGTAGTATCCCCCCAAGTACACCTT
CEY650          : AAATTCCCTAACTTTAAATTTGCCAAGCACATCCACACTCCCGTATTTTTCACAGGGCCAGGTAATATCCCCCCAAGTACACCTT
CEY653          : AAATTCCCTAACTTTAAATTTGCCAAGCACATCCACACTCCCGTATTTTTCACAGGGCCAGGTAATATCCCCCCAAGTACACCTT
CEY649          : AAATTCCCTAACTTTAAATTTGCCAAGCACATCCACACTCCCGTATTTTTCACAGGGCCAGGTAATATCCCCCCAAGTACACCTT
YJM1250         : AAATTCCCTAATTTTAAATTTGCCAAGCACATCCACACTCCCGTATTTTTCACAGGGCCAGGTAATATCCCCCCAAGTACACCTT
SA.9.4.BR2      : AAATTCCCTAACTTTAAATTTGCCAAGCACATCCACACTCCCGTATTTTTCACAGGGCCAGGTAATATCCCCCCAAGTACACCTT
YJM1444         : AAATTCCCTAAATTTAAATTTGCTAAGCACATTCATACTCCTGTATTTTTCACAGGCCCAGGTAATATTCCACCAAGCACACCTT
GLBRCY22_3      : AAATTCCCTAAATTTAAATTTGCTAAGCACATTCATACTCCTGTATTTTTCACAGGCCCAGGTAATATTCCACCAAGCACACCTT
UWOPS87-2421    : AAATTCCCTAAATTTAAATTTGCTAAGCACATTCATACTCCTGTATTTTTCACAGGCCCAGGTAATATTCCACCAAGCACACCTT
YJM653_1b       : AAATTCCCTAAATTTAAATTTGCTAAGCACATTCATACTCCTGTATTTTTCACAGGCCCAGGTAATATTCCACCAAGCACACCTT
YJM681          : AAATTCCCTAAATTTAAATTTGCTAAGCACATTCATACTCCTGTATTTTTCACAGGCCCAGGTAATATTCCACCAAGCACACCTT
EC1118          : AAATTCCCTAAATTTAAATTTGCTAAGCACATTCATACTCCTGTATTTTTCACAGGCCCAGGCAATATTCCACCAAGCACACCTT
S288c           : AAATTCCCTAAATTTAAATTTGCTAAGCACATTCATACTCCTGTATTTTTCACAGGCCCAGGTAATATTCCACCAAGCACACCTT

                       *        2140         *        2160         *        2180         *        2200         *
H4              : ATAACTACTCATTATTTTTCGCAGTGTCATTTTGCCTAACTTTATAAGAAAAAGATGGAGAGCTTGGTTCAATAAATATAATTT
RP11.4.14       : ATAACTACTCATTATTTTTCGCAGTGTCATTTTGCCTAACTTTATAAGAAAAAGATGGAGAGCTTGGTTCAATAAATATAATTT
UFRJ50816       : ATAACTACTCATTATTTTTCGCAGTGTCATTTTGCCTAACTTTATAAGAAAAAGATGGAGAGCTTGGTTCAATAAATATAATTT
YPS138          : ATAACTACTCATTATTTTTCGCAGTGTCATTTTGCCTAACTTTATAAGAAAAAGATGGAGAGCTTGGTTCAATAAATATAATTT
UWOPS91-917.1   : ATAACTACTCATTGTTTTTCGCAGTGTCATTTTGCCTAACTTTATCAGAAAAAGATGGAGAGCTTGGTTCAATAAATATAATTT
N44             : ATAACTACTCATTGTTTTTCGCAATGTCATTTTGCCTAACTTTATAAGGAAAAGATGGAGAGCCTGGTTCAATAAGTATAATTT
CBS432          : ATAACTACTCATTATTTTTCGCAATGTCATTCTGCCTAACTTTATAAGGAAAAGATGGAGAGCTTGGTTCAATAAGTATAATTT
CEY647          : ATAACTACTCATTATTTTTCGCAATGTCATTTTGCCTAACTTTATAAGAAAAAGATGGAGAGCTTGGTTCAATAAATATAATTT
CLQCA_20-060    : ATAACTACTCATTATTTTTCGCAGTGTCATTTTGCCTAACTTTATAAGAAAAAGATGGAGAGCTTGGTTCAATAAATATAATTT
CEY650          : ATAACTACTCATTATTTTTCGCAGTGTCATTTTGCCTAACTTTATAAGAAAAAGATGGAGAGCTTGGTTCAATAAATATAATTT
CEY653          : ATAACTACTCATTATTTTTCGCAGTGTCATTTTGCCTAACTTTATAAGAAAAAGATGGAGAGCTTGGTTCAATAAATATAATTT
CEY649          : ATAACTACTCATTATTTTTCGCAGTGTCATTTTGCCTAACTTTATAAGAAAAAGATGGAGAGCTTGGTTCAATAAATATAATTT
YJM1250         : ATAACTACTCATTATTTTTCGCAGTGTCATTTTGCCTAACTTTATAAGAAAAAGATGGAGAGCTTGGTTCAATAAATATAATTT
SA.9.4.BR2      : ATAACTACTCATTATTTTTCGCAGTGTCATTTTGCCTAACTTTATAAGAAAAAGATGGAGAGCTTGGTTCAATAAATATAATTT
YJM1444         : ATAACTACTCATTATTTTTTGCAATGTCATTTTGCCTAACTTGATAAGAAAAAGATGGAGAGCTTGGTTCAATAAGTACAATTT
GLBRCY22_3      : ATAACTACTCATTATTTTTTGCAATGTCATTTTGCCTAACTTGATAAGAAAAAGATGGAGAGCTTGGTTCAATAAGTACAATTT
UWOPS87-2421    : ATAACTACTCATTATTTTTTGCAATGTCATTTTGCCTAACTTGATAAGAAAAAGATGGAGAGCTTGGTTCAATAAGTACAATTT
YJM653_1b       : ATAACTACTCATTATTTTTTGCAATGTCATTTTGCCTAACTTGATAAGAAAAAGATGGAGAGCTTGGTTCAATAAGTACAATTT
YJM681          : ATAACTACTCATTATTTTTTGCAATGTCATTCTGCCTAACTTGATAAGAAAAAGATGGAGAGCTTGGTTCAATAAGTACAATTT
EC1118          : ATAACTACTCATTATTTTTTGCAATGTCATTTTGCCTAACTTGATAAGAAAAAGATGGAGAGCTTGGTTCAATAAGTACAATTT
S288c           : ATAACTACTCATTATTTTTTGCAATGTCATTCTGCCTAACTTGATAAGAAAAAGATGGAGAGCTTGGTTCAATAAGTACAATTT

                        2220         *        2240         *        2260         *        2280         *
H4              : TGTCATGGGGGCCGGTGTTGAAGCGGGCGTGGCAATCTCTGTCGTTATTATCTTTCTGTGTGTACAATATCCAGGTGGCAAACTC
RP11.4.14       : TGTCATGGGGGCCGGTGTTGAAGCGGGCGTGGCAATCTCTGTCGTTATTATCTTTCTGTGTGTACAATATCCAGGTGGCAAACTC
UFRJ50816       : TGTCATGGGGGCCGGTGTTGAAGCGGGCGTGGCAATCTCTGTCGTTATTATCTTTCTGTGTGTACAATATCCAGGTGGCAAACTC
YPS138          : TGTCATGGGGGCCGGTGTTGAAGCGGGCGTGGCAATCTCTGTCGTTATTATCTTTCTGTGTGTACAATATCCAGGTGGCAAACTC
UWOPS91-917.1   : TGTCATGGGGGCCGGTGTTGAAGCGGGCGTGGCAATCTCTGTCGTTATTATCTTTCTGTGTGTACAATATCCAGGTGGCAAACTC
N44             : TGTCATGGGGGCCGGTGTTGAAGCAGGAGTGGCAATCTCTGTCGTTATTATCTTTCTGTGTGTTCAATATCCAGGTGGCAAACTG
CBS432          : TGTCATGGGGGCTGGTGTTGAAGCAGGCGTGGCAATCTCTGTCGTTATCATATTTCTGTGTGTACAATATCCAGGTGGTAAACTC
CEY647          : TGTCATGGGGGCCGGTGTTGAAGCGGGCGTGGCAATCTCTGTCGTTATTATCTTTCTGTGTGTACAATATCCAGGTGGCAAACTC
CLQCA_20-060    : TGTCATGGGGGCCGGTGTTGAAGCGGGCGTGGCAATCTCTGTCGTTATTATCTTTCTGTGTGTACAATATCCAGGTGGCAAACTC
CEY650          : TGTCATGGGGGCCGGTGTTGAAGCGGGCGTGGCAATCTCTGTCGTTATTATCTTTCTGTGTGTACAATATCCAGGTGGCAAACTC
CEY653          : TGTCATGGGGGCCGGTGTTGAAGCGGGCGTGGCAATCTCTGTCGTTATTATCTTTCTGTGTGTACAATATCCAGGTGGCAAACTC
CEY649          : TGTCATGGGGGCCGGTGTTGAAGCGGGCGTGGCAATCTCTGTCGTTATTATCTTTCTGTGTGTACAATATCCAGGTGGCAAACTC
YJM1250         : TGTCATGGGGGCCGGTGTTGAAGCGGGCGTGGCAATCTCTGTCGTTATTATCTTTCTGTGTGTACAATATCCAGGTGGCAAACTC
SA.9.4.BR2      : TGTCATGGGGGCCGGTGTTGAAGCGGGCGTGGCAATCTCTGTCGTTATTATCTTTCTGTGTGTACAATATCCAGGTGGCAAACTC
YJM1444         : CGTCATGGGGGCCGGTGTTGAAGCAGGTGTGGCAATCTCCGTCGTCATCATCTTCTTGTGTGTACAGTACCCAGGTGGTAAACTC
GLBRCY22_3      : CGTCATGGGGGCCGGTGTTGAAGCAGGTGTGGCAATCTCCGTCGTCATCATCTTCTTGTGTGTACAGTACCCAGGTGGTAAACTC
UWOPS87-2421    : CGTCATGGGGGCCGGTGTTGAAGCAGGTGTGGCAATCTCCGTCGTCATCATCTTCTTGTGTGTACAGTACCCAGGTGGTAAACTC
YJM653_1b       : CGTCATGGGGGCCGGTGTTGAAGCAGGTGTGGCAATCTCCGTCGTCATCATCTTCTTGTGTGTACAGTACCCAGGTGGTAAACTC
YJM681          : CGTCATGGGGGCCGGTGTTGAAGCAGGTGTGGCAATCTCCGTCGTCATCATCTTCTTGTGTGTACAGTACCCAGGTGGTAAGCTC
EC1118          : CGTCATGGGGGCCGGTGTTGAAGCAGGCGTGGCAATCTCCGTCGTCATCATCTTCTTGTGTGTACAGTACCCAGGTGGTAAGCTC
S288c           : CGTCATGGGGGCCGGTGTTGAAGCAGGTGTGGCAATCTCCGTCGTCATCATCTTCTTGTGTGTACAGTACCCAGGTGGTAAGCTC
```

```
                      2300          *        2320          *        2340          *        2360          *        2380
H4                : AGCTGGTGGGGAAACAACGTTTGGAAAAGAACTTATGATAATGATTATATAAAAAATTCTACACCTTAAAGAAAGGTGAGACATTCG
RP11.4.14         : AGCTGGTGGGGAAACAACGTTTGGAAAAGAACTTATGATAATGATTATATAAAAAATTCTACACCTTAAAGAAAGGTGAGACATTCG
UFRJ50816         : AGCTGGTGGGGAAACAACGTTTGGAAAAGAACTTATGATAATGATTATATAAAAAATTCTACACCTTAAAGAAAGGTGAGACATTCG
YPS138            : AGCTGGTGGGGAAACAACGTTTGGAAAAGAACTTATGATAATGATTATATAAAAAATTCTACACCTTAAAGAAAGGTGAGACATTCG
UWOPS91-917.1     : AGCTGGTGGGGAAACAACGTTTGGAAAAGAACTTATGATAATGATTATATAAAAAATTCTACACCTTAAAGAAAGGTGAGACATTCG
N44               : AGCTGGTGGGGAAACAACGTTTGGAAAAGAACGTATGATAATGATTATATAAAAAATTCTATACCTTAAAGAAAGGTGAGACATTCG
CBS432            : AGCTGGTGGGGAAACAACGTTTGGAAAAGAACGTATGATAATGATTATATAAAAAATTCTATACCTTAAAGAAAGGTGAGACATTCG
CEY647            : AGCTGGTGGGGAAACAACGTTTGGAAAAGAACTTATGATAATGATTATATAAAAAATTCTACACCTTAAAGAAAGGTGAGACATTCG
CLQCA_20-060      : AGCTGGTGGGGAAACAACGTTTGGAAAAGAACTTATGATAATGATTATATAAAAAATTCTACACCTTAAAGAAAGGTGAGACATTCG
CEY650            : AGCTGGTGGGGAAACAACGTTTGGAAAAGAACTTATGATAATGATTATATAAAAAATTCTACACCTTAAAGAAAGGTGAGACATTCG
CEY653            : AGCTGGTGGGGAAACAACGTTTGGAAAAGAACTTATGATAATGATTATATAAAAAATTCTACACCTTAAAGAAAGGTGAGACATTCG
CEY649            : AGCTGGTGGGGAAACAACGTTTGGAAAAGAACTTATGATAATGATTATATAAAAAATTCTACACCTTAAAGAAAGGTGAGACATTCG
YJM1250           : AGCTGGTGGGGAAACAACGTTTGGAAAAGAACTTATGATAATGATTATATAAAAAATTCTACACCTTAAAGAAAGGTGAGACATTCG
SA.9.4.BR2        : AGCTGGTGGGGAAACAACGTTTGGAAAAGAACTTATGATAATGATTATATAAAAAATTCTACACCTTAAAGAAAGGTGAGACATTCG
YJM1444           : AGCTGGTGGGGAAACAACGTTTGGAAAAGAACGTATGATAATGATTATATAAAAAATTCTACACCTTAAAGAAAGGTGAGACATTCG
GLBRCY22_3        : AGCTGGTGGGGAAACAACGTTTGGAAAAGAACGTATGATAATGATTATATAAAAAATTCTACACCTTAAAGAAAGGTGAGACATTCG
UWOPS87-2421      : AGCTGGTGGGGAAACAACGTTTGGAAAAGAACGTATGATAATGATTATATAAAAAATTCTACACCTTAAAGAAAGGTGAGACATTCG
YJM653_1b         : AGCTGGTGGGGAAACAACGTTTGGAAAAGAACGTATGATAATGATTATATAAAAAATTCTACACCTTAAAGAAAGGTGAGACATTCG
YJM681            : AGCTGGTGGGGAAACAACGTTTGGAAAAGAACGTATGATAATGATTATATAAAAAATTTTATACCTTAAAGAAAGGTGAGACATTTG
EC1118            : AGCTGGTGGGGAAACAACGTTTGGAAAAGAACGTATGATAATGATTATATAAAAAATTTTATACCTTAAAGAAAGGTGAGACATTTG
S288c             : AGCTGGTGGGGAAACAACGTTTGGAAAAGAACGTATGATAATGATTATATAAAAAATTTTATACCTTAAAGAAAGGTGAGACATTTG
```

```
                              *        2400
H4                : GTTATGATAAATGGTGGTAG
RP11.4.14         : GTTATGATAAATGGTGGTAG
UFRJ50816         : GTTATGATAAATGGTGGTAG
YPS138            : GTTATGATAAATGGTGGTAG
UWOPS91-917.1     : GTTATGATAAATGGTGGTAG
N44               : GTTATGATAAATGGTGGTAG
CBS432            : GTTATGATAAATGGTGGTAG
CEY647            : GTTATGATAAATGGTGGTAG
CLQCA_20-060      : GTTATGATAAATGGTGGTAG
CEY650            : GTTATGATAAATGGTGGTAG
CEY653            : GTTATGATAAATGGTGGTAG
CEY649            : GTTATGATAAATGGTGGTAG
YJM1250           : GTTATGATAAATGGTGGTAG
SA.9.4.BR2        : GTTATGATAAATGGTGGTAG
YJM1444           : GTTATGATAAATGGTGGTAG
GLBRCY22_3        : GTTATGATAAATGGTGGTAG
UWOPS87-2421      : GTTATGATAAATGGTGGTAG
YJM653_1b         : GTTATGATAAATGGTGGTAG
YJM681            : GTTATGATAAATGGTGGTAA
EC1118            : GTTATGATAAATGGTGGTAA
S288c             : GTTATGATAAATGGTGGTAA
```

**SUPPLEMENTARY FIGURE 4|** Multiple sequence alignment of *OPT1* nucleotide sequences displaying gene chimerism. Strains names in red indicate *OPT1* sequences from the respective *S. cerevisiae* isolates, while blue labeled strains represent *OPT1* sequences from *S. paradoxus* isolates. The color scheme for nucleotide sequences indicate in red capital letters (and dark blue background) nucleotide sequences related to S288c form of the *OPT1* gene, whereas in white capital letters (and light blue background) are nucleotides related to the *S. paradoxus OPT1* form. Note that in many *S. cerevisiae* strains there are different recombination points in which the *S. paradoxus* sequence starts to prevail along the sequence. Strain names and accession numbers are described on the **Supplementary Table 12**.

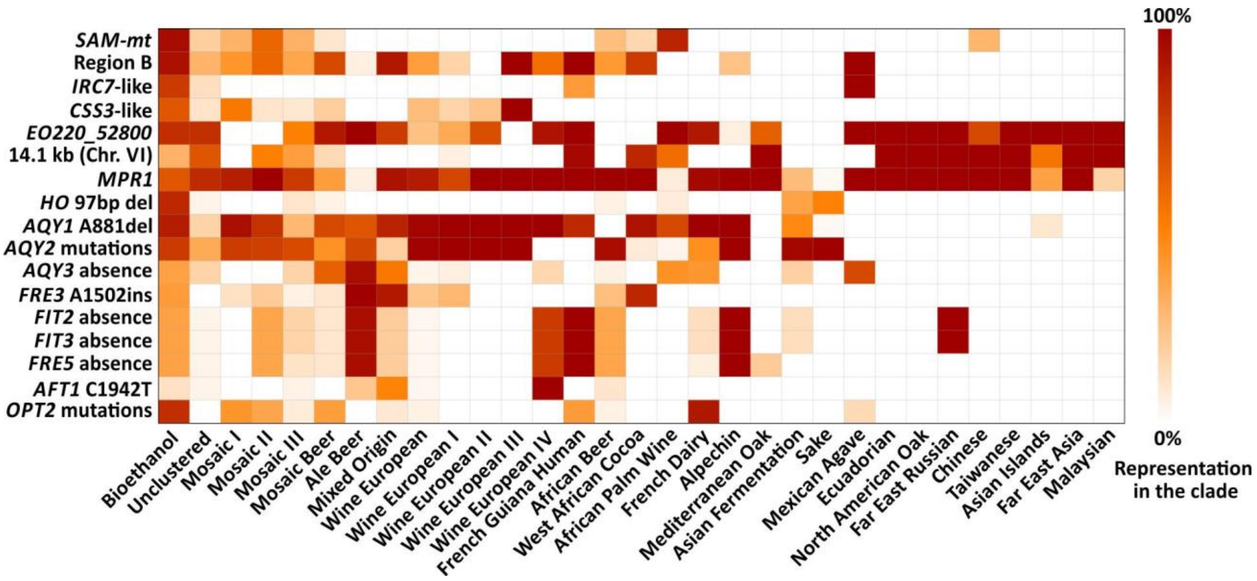


**SUPPLEMENTARY FIGURE 5|** Heatmap depicting the representation (%) across 1,020 yeasts of selected features observed in bioethanol strains. The presence or lack of the analyzed features (listed in the y-axis and representing specific genes, mutated alleles, or absence of genes) was estimated for each clade from the BLASTN hits. For each group, the fraction (%) of representation is schematically depicted by a color gradient from 0% to 100% (right bar). Yeasts are grouped along the x-axis in 31 clades, according to Peter et al., 2018 (the "Chinese" group, as labeled here, combines the clades CHNI, II, III, and V). BLASTN parameters, query sequences, and defined criteria to estimate the presence or absence of alleles are shown in the **Supplementary Table 2**. For *AQY3*, *FIT2*, *FIT3*, and *FRE5*, the heatmap shows the extent of their absence within yeast clades. *AQY2* mutations are the 11bp del, G25del, and C424T (see **Figure 3A** on the main text). *OPT2* mutations are T560ins, AA777ins, T1020del, and other variants (A41del, C1224A, 1499 17bp ins, and 1519 17bp ins).

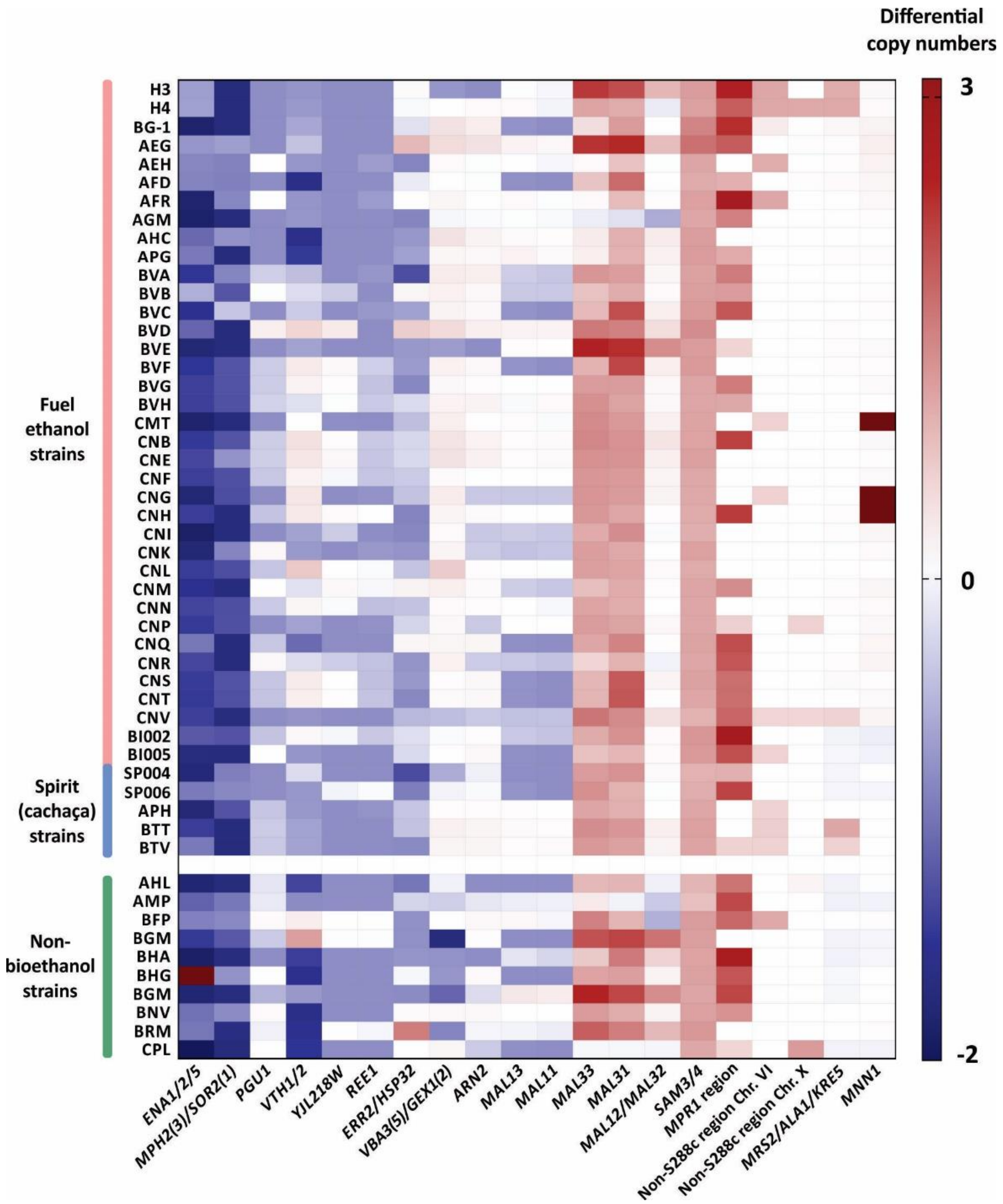


**SUPPLEMENTARY FIGURE 6|** Genes/regions with similar CNVs in both bioethanol and non-bioethanol strains. CNVs of key genes/regions (labeled at the bottom) were estimated in bioethanol and non-bioethanol yeasts based on read depth and discounted from the copy numbers deduced from the S288c reference genome. Only genes/regions in which CNVs tend to be similar in both bioethanol and non-bioethanol strains are shown. This suggests that most variability lies within the S288c lineage, or that the differences observed are not exclusive to the bioethanol group. A colored scheme is applied in which extra copies in bioethanol strains are quantitatively expressed in a red gradient (up to three copies). Equal number of copies in both bioethanol and S288c strains is represented in white, while surplus of copy numbers in S288c are displayed as a blue gradient. A group of ten non-bioethanol yeasts were included for comparison. Dark-red boxes outside the red gradient range represent a few cases when more than three extra copies of the probed gene/region were counted. This exception is the case of the *MNN1* gene, which is represented by about 6, 10, and 11 copies in the CNH, CNG, and CMT strains, respectively.

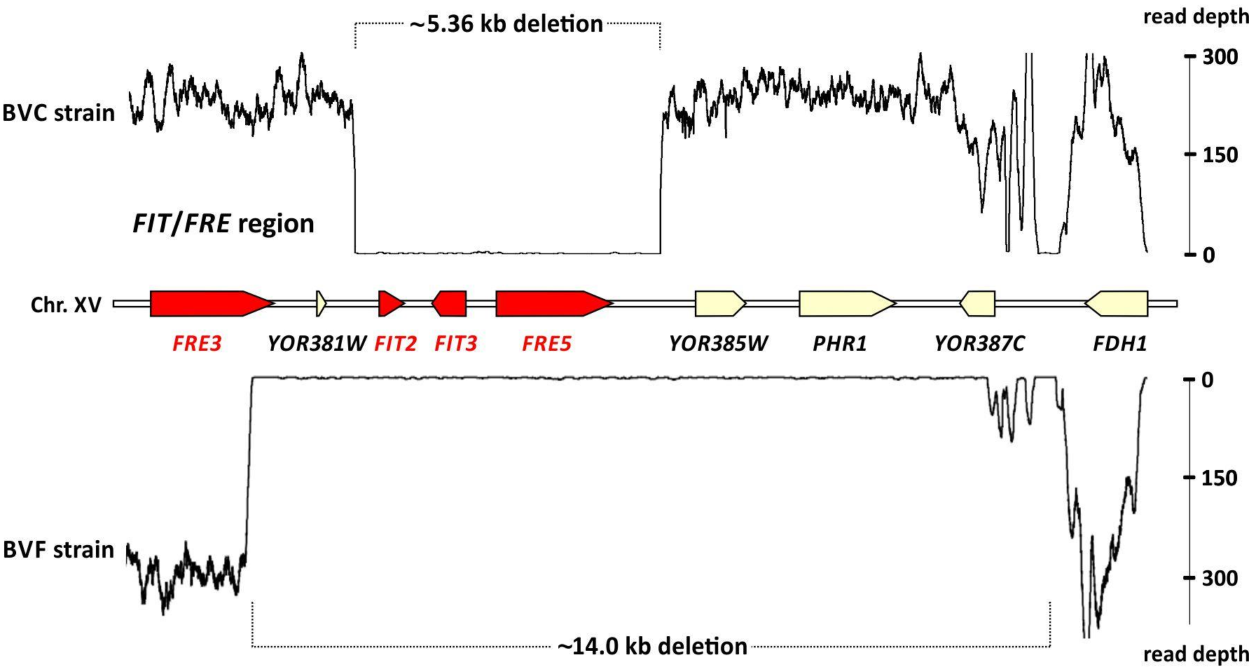


**SUPPLEMENTARY FIGURE 7|** Deletion of the *FIT*/*FRE* region in bioethanol strains. The *FIT*/*FRE* region on the Chr. XV of *S. cerevisiae* S288c is shown in the middle with the *FRE3*, *FIT2*, *FIT3*, and *FRE5* genes highlighted in red. Illumina reads from the genome sequence of strain BVC were mapped against the S288c genome. A read depth plot above the Chr. XV segment reveals a gap of about ~5.36 kb. This denotes a deletion of the genes *FIT2*, *FIT3*, and *FRE5*. The same ~5.36 kb deletion is observed in other nine bioethanol strains. Similarly, when Illumina reads from the genome sequencing of the strain BVF are mapped against the S288c genome a major deletion of ~14.0 kb is seen on the read depth plot below the chromosomal segment that includes the genes *FIT2*, *FIT3*, and *FRE5*. The 14.0 kb deletion has a left breaking point at the 3' region of *FRE3*. The same deletion is observed in other 21 bioethanol strains (**Figure 4**, main text).

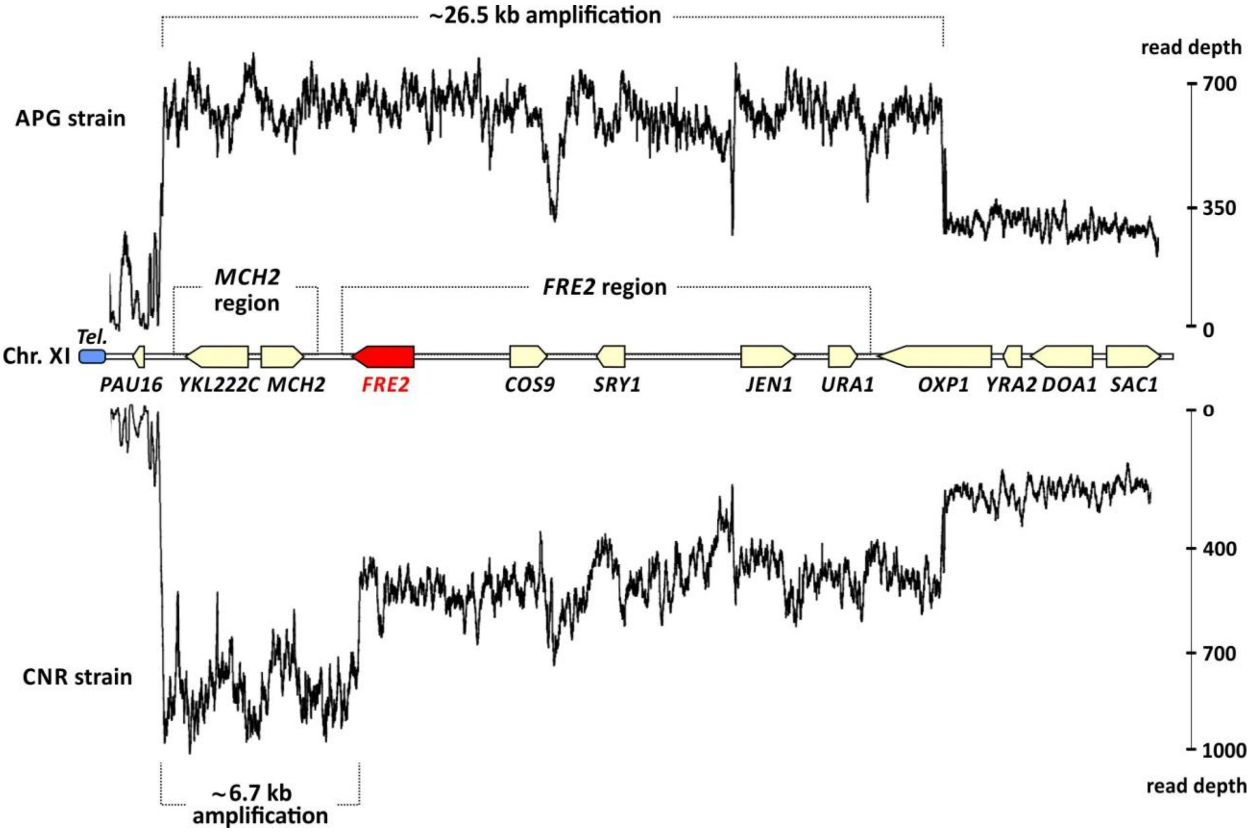


**SUPPLEMENTARY FIGURE 8|** Two patterns of amplification over the *MCH2* and *FRE2* regions in bioethanol yeasts. The *MCH2*/*FRE2* regions on the Chr. XI of *S. cerevisiae* S288c are shown in the middle, with *FRE2* highlighted in red. Illumina reads from the genome sequence of strain APG were mapped against the S288c genome. Above the chromosome diagram, a read depth plot of the Chr. XI left-end reveals a sudden raise in coverage, indicating a segmental duplication of about 26.5 kb that involves *FRE2*. In another example, depicted below the chromosome diagram, the mapping of Illumina reads from the genome sequence of strain CNR against the genome of S288c also recapitulates the same ~26.5 kb amplification. However, an increase in read depth indicates a further round of amplification over the ~6.7 Kb *MCH2* region.

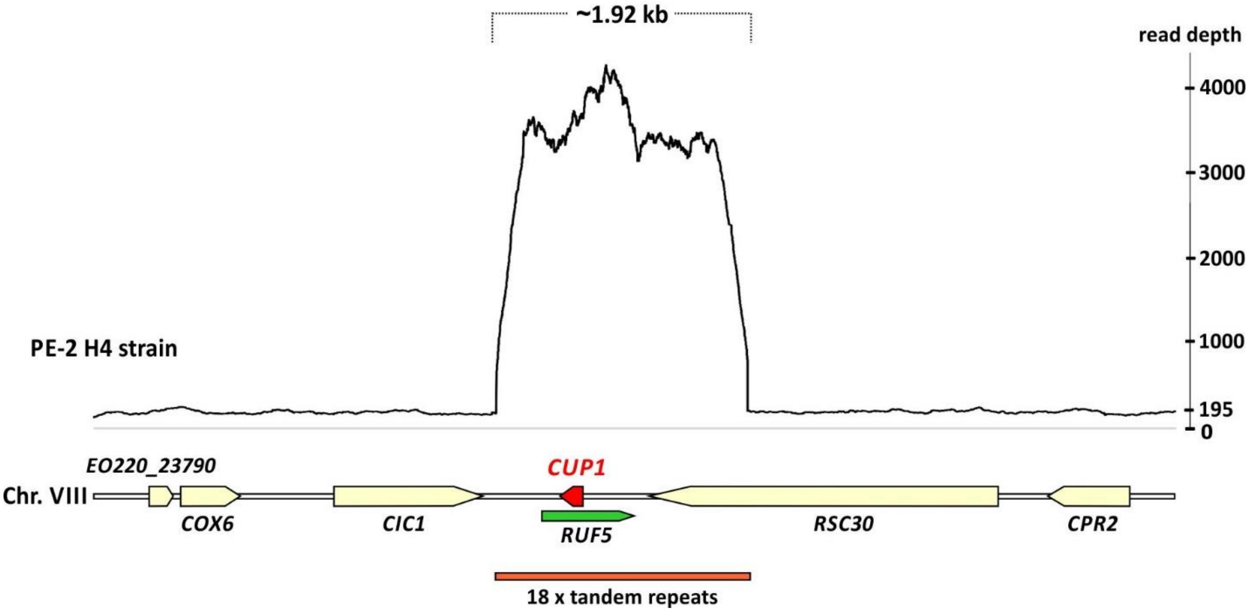


**SUPPLEMENTARY FIGURE 9|** Amplification of *CUP1*. Read depth plot over a segmental region of H4 Chr. VIII indicates about 18 copies of *CUP1*, represented in red. The type-IV tandem repeats of ~1.9 kb in length (Zhao et al., 2014) is supported by PacBio long reads. *RUF5* in green represents a non-coding RNA associated with the *CUP1* locus.

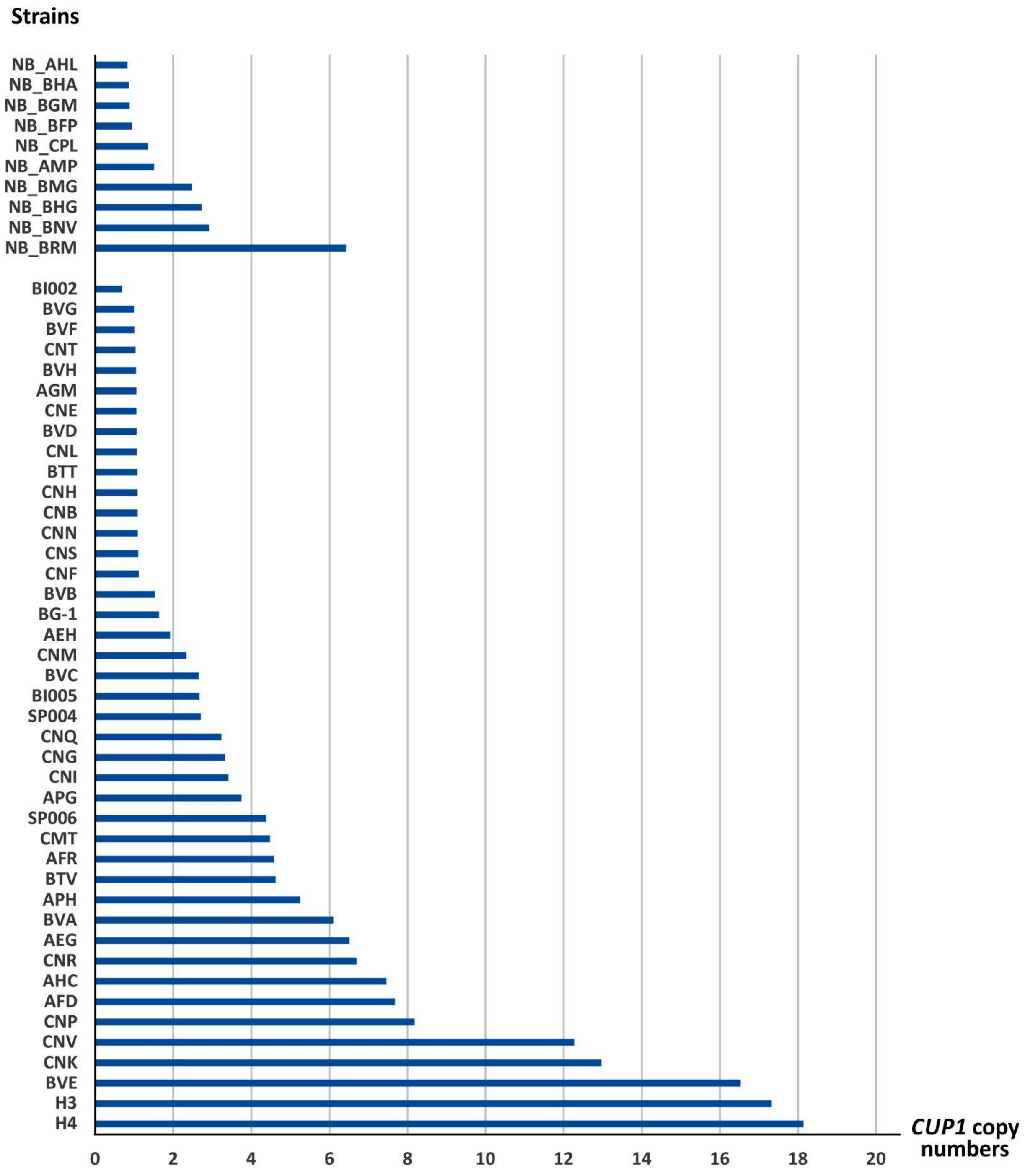


**SUPPLEMENTARY FIGURE 10|** *CUP1* copy numbers in strains from the bioethanol group. A graphical representation shows the estimated *CUP1* absolute copy numbers across 42 strains of the bioethanol group (bottom) and, for comparison, 10 non-bioethanol (NB_) yeasts (top). Copy numbers were calculated for each yeast based on Illumina read depth over the region length, and normalized for the strain specific genomic coverage.

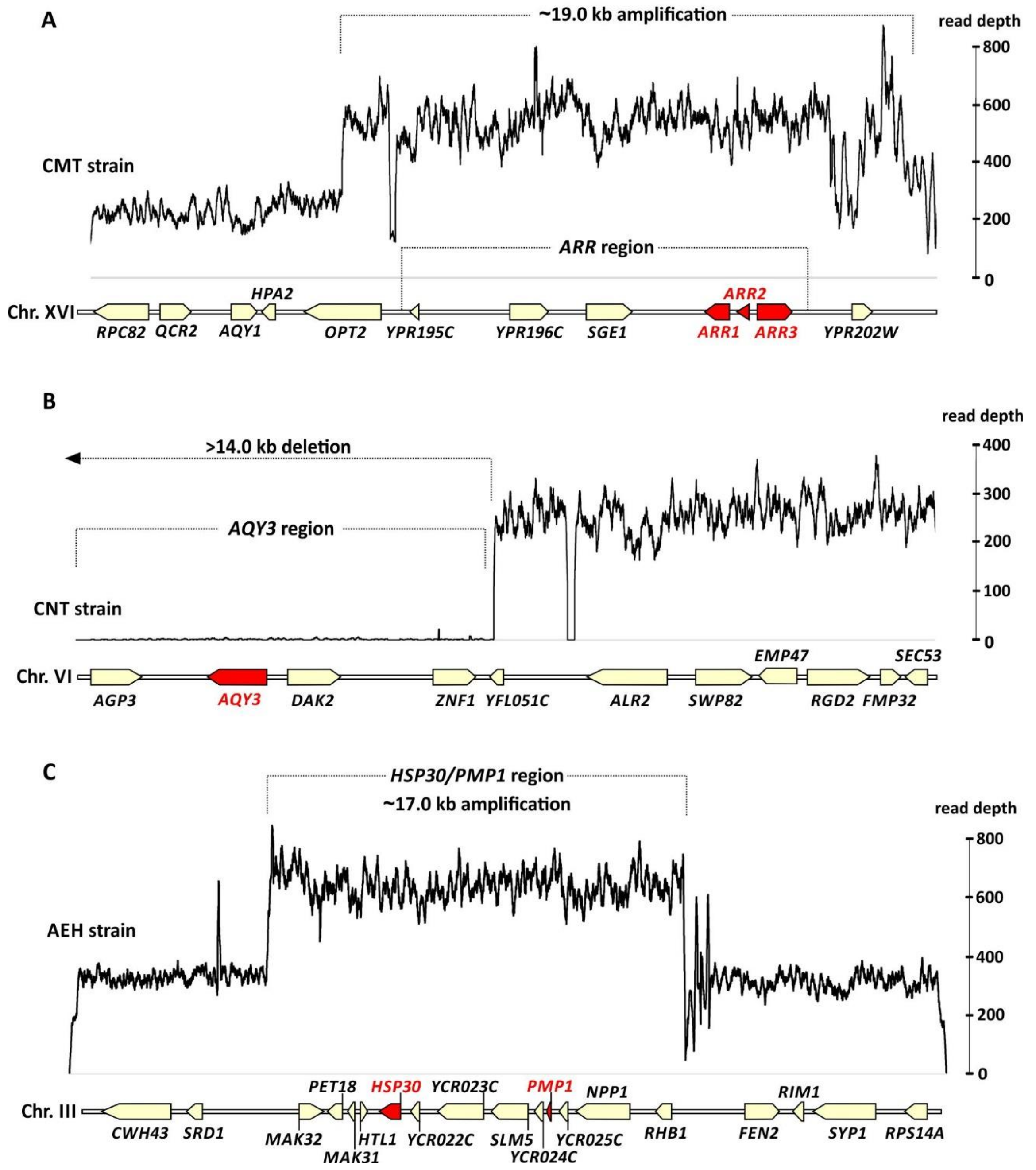


**SUPPLEMENTARY FIGURE 11|** CNVs over larger genomic regions of bioethanol strains. Read depth plots for three different bioethanol strains are shown over three respective S288c chromosomic regions. Genes putatively under selection are depicted in red. **(A)** When reads from the strain CMT are mapped against the S288c genome an amplification is observed over a ~19.0 kb region of Chr. XVI encompassing the *ARR* genes (in red) for arsenic metabolism. **(B)** *AQY3* region in S288c. Complete absence of mapped reads over the ~14.0 kb region indicates its deletion in the CNT strain when compared to S288c. A similar pattern is observed in a further 17 strains from the bioethanol group. It is possible that the deletion extends beyond the 14.0 kb region. **(C)** Read depth plot of strain AEH over the S288c Chr. III indicates a ~17.0 kb amplification involving 13 genes.

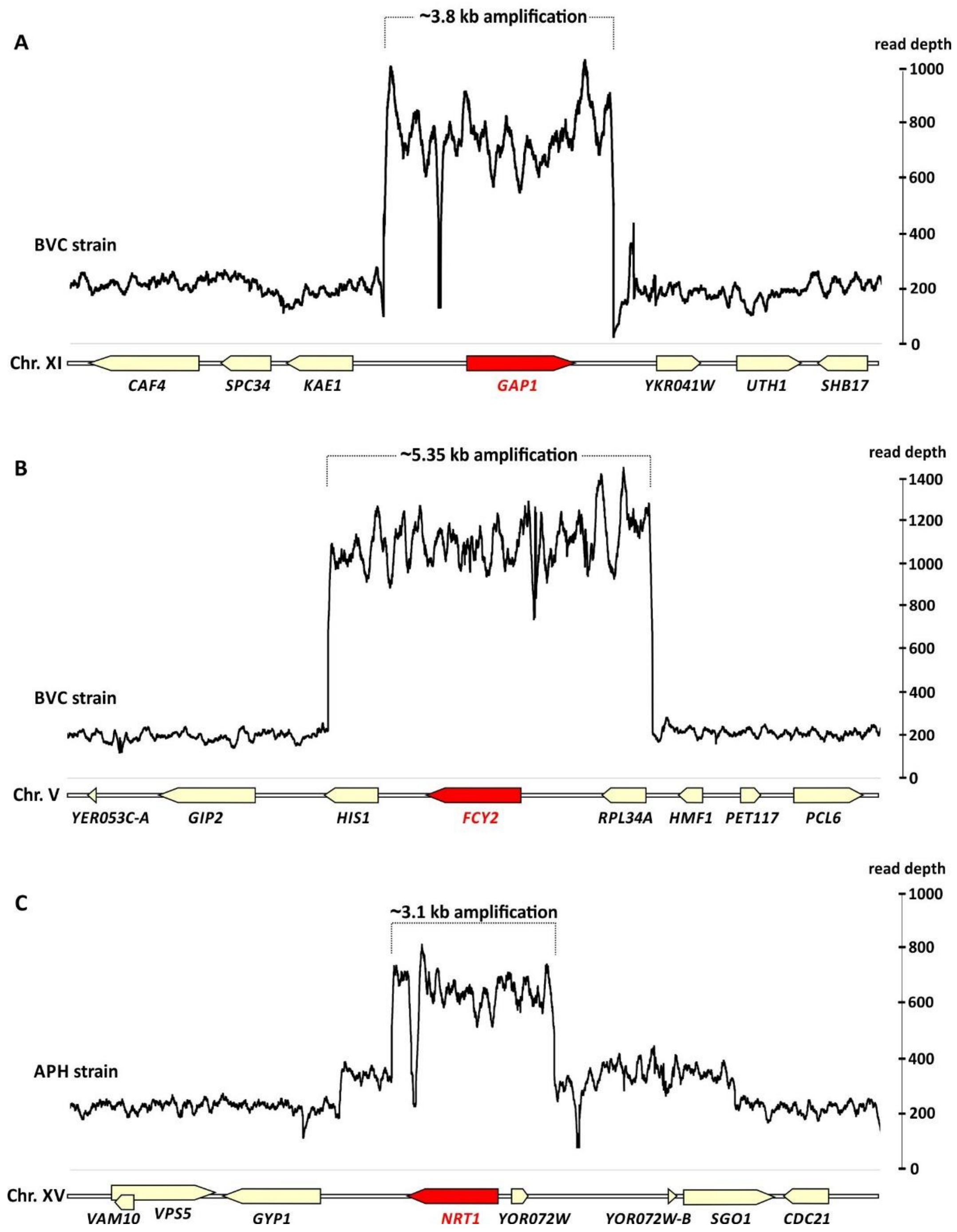


**SUPPLEMENTARY FIGURE 12|** Amplification of nutrient-acquisition genes in bioethanol and cachaça strains. Read mapping plots of the strain BVC against the genome of S288c reveal (**A**) a ~3.8 kb amplification encompassing the general amino acid permease *GAP1* (in red) at the Chr. XI that is significantly enriched among strains of the bioethanol group, and (**B**) the amplification of *FCY2* (encoding a purine-cytosine permease) within a ~5.35 kb region of Chr. V. **(C)** The APH cachaça strain has a 3.1 kb amplificated region on Chr. XV when compared with the S288c reference. This amplification encompasses *NRT1* (in red) and is shared by other cachaça yeasts in our dataset.
